# Spatial proximity to AD pathology defines cell-type and genotype-specific characteristics in human AD hippocampus

**DOI:** 10.64898/2026.10.15.759523

**Authors:** Christina Huan Shi, Bing Zhu, Juan Pina-Crespo, Win Ning Chen, Giau Van Vo, Huijie Huang, Kevin Y. Yip, Timothy Y. Huang

**Affiliations:** Center for Neurologic Diseases, Sanford Burnham Prebys Medical Discovery Institute, La Jolla, CA 92037, USA; Center for Data Science and Artificial Intelligence, Sanford Burnham Prebys Medical Discovery Institute, La Jolla, CA 92037, USA; Cancer Genome and Epigenetics Program, NCI-Designated Cancer Center, Sanford Burnham Prebys Medical Discovery Institute, La Jolla, CA 92037, USA

**Keywords:** Alzheimer’s disease, hippocampus, spatial transcriptomics, eQTL, MERFISH, APOE

## Abstract

How amyloid-β (Aβ) plaques and tau phosphorylation spatially relate to cell response and genetic variation in Alzheimer’s disease (AD) brain is currently unclear. We established a pipeline to concurrently annotate Aβ and AT8 ptau pathology and cell-specific MERFISH expression profiles in human control and AD hippocampus. We characterized expression signatures in glia and neurons proximal or distal to Aβ plaques, AT8 ptau-enriched neurons, or Aβ and AT8 ptau in combination, and used spatially-defined transcriptional signatures to infer pathology-associated states in large snRNA-seq datasets lacking spatial context. We characterized variants from 120 gene loci associated with AD risk that affect cell-specific gene expression within local pathological contexts. Astrocyte APOE4/E4 expression was specifically suppressed in proximity to Aβ plaques, whereas other APOE genotypes were unaffected by Aβ proximity. Together, this analysis pipeline couples expression profiles in proximity to Aβ and tau pathology, and implicates AD cis-gene variants with spatial sensitivity to AD pathology.

## Introduction

Alzheimer’s disease is a neurological disorder that manifests cognitive decline with age, and is defined by the deposition of amyloid-β (Aβ) plaques and appearance of neurofibrillary tangles comprising pathological tau phosphoforms during progression of the disease (DeTure and Dickson 2019, Rawat, Sehar et al. 2022). Aβ accumulation typically accumulates in neocortical regions around two decades prior to signs of memory dysfunction (Grothe, Barthel et al. 2017, Quiroz, Sperling et al. 2018), whereas tau spreading originates within transentorhinal and entorhinal cortices and subsequently spreads into association and unimodal cortex regions (Braak, Thal et al. 2011, Kaufman, Del Tredici et al. 2018, Vogels, Leuzy et al. 2020). Tau pathology can manifest independently of Aβ during aging; age-related accumulation of neurofibrillary tangles can occur in the medial temporal lobe (MTL) comprising the entorhinal cortex, hippocampus and amygdala regions in individuals around 50 or 60 years of age (Braak and Braak 1997, Braak, Thal et al. 2011). Tau tangles that coincide with aging appear early in life (Braak and Del Tredici 2011), and tau pathology in the absent of Aβ in non-neuropathological “primary age-related tauopathy” (PART) is often not associated with dementia or significant cognitive impairment (Crary, Trojanowski et al. 2014). Interestingly, the appearance of Aβ within the MTL is thought to potentiate the tau spreading beyond the MTL (Jacobs, Hedden et al. 2018), which supports a model where Aβ can induce tau spreading (Musiek and Holtzman 2012, Jagust 2018). Thus, spatial convergence of Aβ and tau pathology could potentially have effects on the cellular features with significant consequences during AD onset (Shi, Pina-Crespo et al. 2026).

Decreased hippocampal volume and progressive atrophy is associated with cognitive impairment and clinical dementia (Barnes, Bartlett et al. 2009, Fjell and Walhovd 2010, Fraser, Shaw et al. 2015, Peng, Feng et al. 2015, Nobis, Manohar et al. 2019). Hippocampal atrophy appears to correlate with the initiation of AD onset (Li, Coyle et al. 2011) and sustained atrophy occurs along the progression of AD (Whitwell, Przybelski et al. 2007). Neurofibrillary tangle burden is inversely correlated with antemortem hippocampal volume measured near the end of life (Silbert, Quinn et al. 2003). Hippocampal Aβ pathology or accumulation also correlates with hippocampal atrophy in AD (Mattsson, Insel et al. 2014, Nosheny, Insel et al. 2015, Huang, Shan et al. 2023), yet shows poor correlation with atrophy or cognitive impairment in control subjects (Insel, Mattsson et al. 2015). This suggests that hippocampal changes in normal aging and clinical AD onset can share pathological features such as Aβ deposition and accumulation, yet result in distinct functional consequences.

Numerous genetic variants have been associated with increased AD risk, which include rare familial AD variants in APP, PS1 and PS2 which consequently result in early onset AD (prior to age 60) with high penetrance (Tanzi 2012), as well as numerous variants associated with sporadic late onset AD (age 65 and over) (Khani, Gibbons et al. 2022). Various genome-wide association studies (GWAS) have identified numerous gene loci and potential risk variants linked to AD (Eadb, Eadi et al. 2026), including genes such as *TREM2*, *SORL1* (encoding SORLA), *CD33*, *ABCA7*, *MS4A6A* and *MS4A4A* with enriched expression in microglia (McQuade and Blurton-Jones 2019). Variants in the lipid/cholesterol carrier APOE comprise three major alleles of which APOE3 is considered to be risk-neutral, while APOE2 is associated with reduced AD risk and APOE4 confers increased risk of AD onset (Zhao, Liu et al. 2018). AD risk genes also functionally converge through shared molecular pathways; for example, receptor-ligand interactions have been described for trafficking receptors such as SORL1 and immune signaling receptors such as TREM2 and APOE (Yajima, Tokutake et al. 2015, Yeh, Wang et al. 2016). Moreover, sporadic risk genes such as TREM2 and APOE have been implicated as induced disease associated microglia (DAM) signatures in 5xFAD (Keren-Shaul, Spinrad et al. 2017) as well as neurodegenerative microglia (MGnD) signatures in APP/PS1 AD mouse models (Krasemann, Madore et al. 2017). The impact of various AD-associated risk genes on Aβ and tau pathology and spreading has also been described. For example, murine *Trem2* deletion ameliorates Aβ pathology in young APP/PS1 animals (Jay, Miller et al. 2015, Jay, Hirsch et al. 2017), yet exacerbates Aβ in aged APP/PS1 (Jay, Hirsch et al. 2017) and 5xFAD mice (Wang, Cella et al. 2015). Deletion of murine *Trem2* can also enhance exosome-mediated tau spreading in young mice (Zhu, Liu et al. 2022), and reduce tau dispersion in older animals (Lee-Gosselin, Jury-Garfe et al. 2023). *Trem2* deletion also enhances tau pathology in the presence of Aβ pathology in 5xFAD (Gratuze, Chen et al. 2021) and TauPS2APP (Lee, Meilandt et al. 2021) mouse brain. SORLA deletion can exacerbate Aβ levels (Andersen, Reiche et al. 2005) and PS19 tau pathology (Huang, Shi et al. 2026) in mice, while SORLA overexpression can suppress Aβ (Caglayan, Takagi-Niidome et al. 2014) as well as tau pathology in mouse models of AD/tauopathy (Huang, Shi et al. 2026). Although these results together implicate potential relationships between the modulation of genes related to AD risk and pathological outcome, whether spatial proximity to Aβ or tau pathology is consequential to the expression of AD-associated gene transcripts and whether this correlates with functional readouts in various cell types is unclear.

Here, we characterize cell-type-specific expression of variants in 120 potential AD-risk genes as well as genes related to cellular function in response to their proximity to Aβ plaques or AT8 ptau pathology individually or in combination in control and AD hippocampus (Table S1) using a probe panel for the MERFISH platform, which provides subcellular resolution. In establishing this high-definition expression atlas (neuro-degeneration.org/admerfish), we relate and compare spatially-defined transcriptomic templates to AD snRNA-seq datasets and define genotype-specific cellular expression profiles of AD-risk genes such as *MS4A4A* and *APOE* in relation to Aβ pathology in microglia and astrocytes.

## Results

### Annotating cell types and comparing cell expression profiles in control and AD hippocampus

Current single cell sequencing and characterization methods can catalogue high resolution transcriptomic features in numerous cell types in brain, and have been used extensively to define differences in cell expression profiles in control and AD brain (Ma, Xia et al. 2025). However, these methods are unable to define direct spatial relationships between enriched pathological features such as amyloid plaques and tau phosphorylation and gene expression profiles in various cell types. We therefore employed a MERFISH panel comprising 120 genes related to AD risk (Table S2), as well as probes related to annotate various cell types and to define glial/neuronal activation, function or signatures previously observed in mouse or AD brain (Figure 1A; Figure S1A, Table S3). These cell-type-specific expression profiles are then characterized in relation to Aβ plaques and AT8 ptau (pS202/pT205) pathology individually or in combination (Figure 1A). We also characterized the genomic sequence profiles from these individuals, and called variants of 120 potential AD-risk genes (Figure 1A). We identified postmortem hippocampal samples from 6 control (age ranging from 53 to 79, all Braak stage 1) and 7 AD individuals (ranging from 60 to 89, Braak stage 4-6) (Figure 1B, Table S1) with RIN values ranging from 5 to 7.1. Sections were processed for immunohistological staining and MERSCOPE analysis to characterize Aβ plaque and AT8 ptau pathology multiplexed with concurrent analysis of transcriptomic features using the MERFISH probe panel *in situ* (Figure 1A).

**Figure 1.**
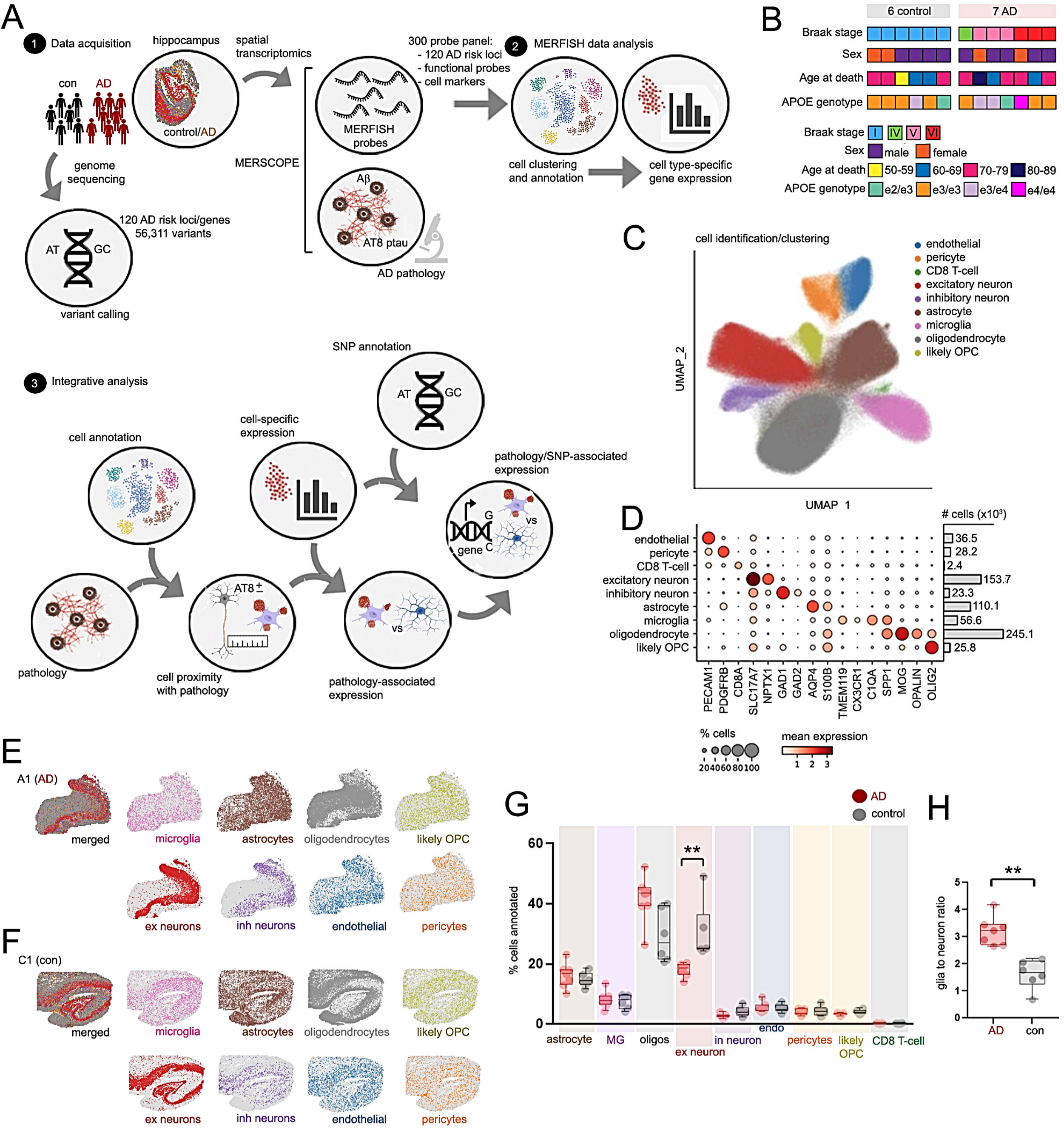
Characterizing expression profiles in relation to pathology in control and AD hippocampus. (A) Study design and analysis pipeline. (B) Patient information of control and AD individuals; color-coded maps indicate Braak stage, sex, APOE genotype and age. (C) UMAP visualization of annotated cell types. Colored indicators are shown for various cell types. (D) Dot plot indicating expression of markers for annotated hippocampal cell types. Size scales indicate percentage expression, and color scales depict mean expression values; total number of cells are indicated in the adjacent bar graph. (E, F) Representative rendered images of annotated cell types in AD (E) and control (F) hippocampus. (G) Percentage of cell types in AD (red) and control (gray) hippocampus. (H) Glia to neuron ratio in individual control (“con”, gray) and AD (red) samples. Unpaired Student’s t-tests were used to determine statistically significant differences between cells proportions of each cell type in (G), or glia to neuron ratio in (H) (*p<0.05).

Following filtering for cell quality, we annotated a total of 681,854 cells using established cell type markers, identifying excitatory/inhibitory neurons, glia (oligodendrocytes, microglia and astrocytes), endothelial cells, pericytes, and CD8 T-cells (Figure 1C-F; Figure S1B-D). The annotated cell types predominantly comprised oligodendrocytes and excitatory neurons (Figure S1D), and we observed decreased numbers of excitatory neurons in AD compared to control hippocampus, potentially due to late-stage neuronal degeneration (Figure 1G, H). We further examined two samples from donors with relatively younger and older ages at death, and confirmed that cells from these samples were evenly intermixed with cells from other samples, suggesting that effects were not driven by age differences (Figure S1E-F).

We compared cell-type-specific expression changes in control and AD hippocampus; a total of 133 differentially-expressed genes (DEGs) were observed across glia, neurons, pericytes and endothelial cells (Figure 2A). Among these DEGs were cell-type-specific changes in various AD-risk genes (Figure S2A, B). For example, *APOE* was found to be significantly upregulated overall in AD microglia and was significantly reduced in AD astrocytes (Figure S2A, B). Some AD vs control DEGs were shared between excitatory and inhibitory neurons; genes including *SLC24A4*, *PLEKHA1*, *NFIC*, *WWOX*, *ANK3* and *NYAP1* were upregulated in both excitatory and inhibitory neurons in AD compared with controls (Figure S2C, D). The neurexin-family component *CNTNAP2* was increased in excitatory neurons, and conversely downregulated in inhibitory neurons (Figure S2C, D) in AD compared with controls. *APP* (amyloid precursor protein) was downregulated specifically in AD oligodendrocytes and inhibitory neurons compared with controls (Figure S2A, C). Gene ontology (GO) pathway analysis of AD vs control DEGs revealed enrichment of genes associated with KEGG “Cholesterol metabolism”, “Lysosome” and “Alzheimer’s disease” categories (Figure 2B-D, and S2E), as well as Molecular Function (MF) categories such as “amyloid-binding”, “apolipoprotein binding” and “tau protein binding” (Figure S2F, G).

**Figure 2.**
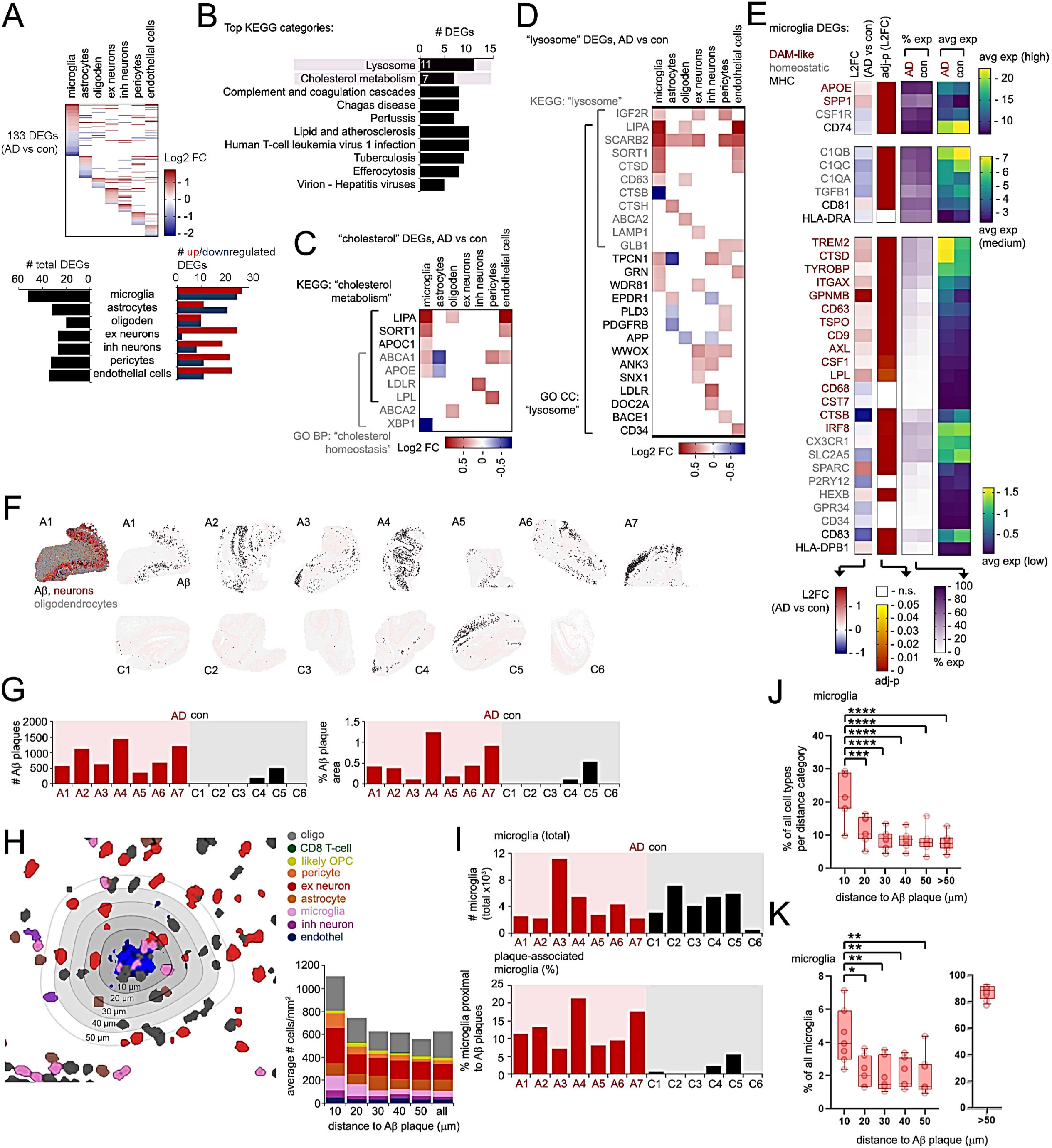
Cell-type-specific DEG analysis, annotation of Aβ plaques and cells proximal to Aβ plaques in AD and control hippocampus. (A) Heatmap of 133 DEGs identified comparing AD vs con (Log2 fold-change, FC) hippocampus in various cell types. Bar graphs indicate number of total (black bars) or upregulated (red) or downregulated (blue bars) DEGs identified for each cell type. (B) Top KEGG categories enriched in DEGs identified in (A). (C, D) Heatmap depicting Log2 fold-change AD vs control DEGs in KEGG “cholesterol metabolism” (C) and “lysosome” (D) categories in the hippocampal cell types indicated. (E) Heatmaps depicting Log2 fold-change (L2FC, red/blue heatmaps), adj-p (red/yellow heatmaps), and average percentage (purple heatmaps) and average expression (yellow, green, blue) values for AD vs control DAM-like (red), homeostatic (gray) and MHC-associated (black) microglia DEGs. DEGs are split according to relatively high, medium and low average expression values. (F) Annotation of Aβ plaques in AD and control hippocampus. Aβ plaques identified in AD (A1-A7) or control (C1-C6) hippocampus are indicated in black. (G) Quantification of Aβ plaque number, and percentage area in AD and control hippocampus in (F). (H-K) Annotation of various cell types in proximity and distal to Aβ plaques. (H) Representative MERSCOPE image of an Aβ plaque (blue, image background in white); cells were stratified according to their distance to the nearest Aβ plaque within different distance intervals: 10 μm (<10 μm, plaque-proximal), 20 μm (10-20 μm), 30 μm (20-30μm), 40 μm (30-40μm) and 50 μm (40-50 μm). Adjacent graph depicts average number of various cell types (normalized per mm^2^) within different distance intervals of Aβ plaques in AD hippocampal sections. (I) Bar graphs depict total number of microglia and percentage of microglia in proximity to plaques (within 50 μm of the plaque boundaries in individual AD (red) and control (black) hippocampal sections. (J, K) Box and whisker plots indicate percentage of microglia in different distance intervals to Aβ plaques in AD samples; percentages were normalized for all cell types within a region category (J), or normalized as a percentage of microglia (K). Statistical significance was determined by one-way ANOVA (**p<0.01, ***p<0.001, ****p<0.0001).

Statistically significant DEGs comparing AD vs control hippocampus in annotated cell types here also reveal upregulation of DAM genes such as *TREM2* and *APOE* in AD vs control microglia (Figure 2E and S2A, B) as well as various DEGs shared and unique to varying cell types (Table S4). In addition to *TREM2* and *APOE*, microglia DAM-like genes such as *SPP1*, *CTSD*, *TYROBP*, *GPNMB* and *CD9* were upregulated, while homeostatic genes such as *CSF1R*, *TGFB1*, *P2RY12* and *CXCR1* were downregulated in AD vs control brain (Figure 2E). These results detail cell-type-specific transcriptional changes in AD hippocampus using spatial transcriptomic profiles of cells measured *in situ*.

### Cell-specific expression profiles in association with Aβ pathology

Aβ plaques were annotated and spatially mapped in control and AD hippocampus (Figure 2F). Although Aβ plaques were enriched in AD samples, we observed a significant number of Aβ plaques in two controls (C4, C5) (Figure 2G). We then mapped annotated cell types in 10-μm distance intervals from the Aβ boundaries up to 50 μm, with the 0-10 μm interval also including cells overlapping the plaques, and classified cells beyond 50 μm separately (Figure 2H). We further assigned a binary annotation of cells in proximity and distal to Aβ plaques, where cells within a 50 μm periphery were annotated as Aβ-proximal (Aβ+), and cells beyond 50 μm were distal to Aβ plaques (Aβ-). Interestingly, we observed enrichment of microglia in proximity to Aβ (Figure 2I-K); microglia comprised on average 22.1% of cells within 10 μm of Aβ plaques in AD hippocampus across all cell types in the same distance interval (Figure 2J), and on average 4.4% of AD microglia were located within 10 μm of Aβ plaques as a proportion of microglia in all distance categories (Figure 2K). We did not observe enrichment of any other cell types within 50 μm regions within Aβ plaques in AD hippocampus (Figure S3A, B), or microglia from the two Control samples with some plaques (Figure S3C).

Given that microglia interactions with Aβ plaques have been shown to be important in limiting Aβ plaque expansion and containing the plaque perimeter (Condello, Yuan et al. 2015), we further clustered microglia based on gene expression profiles and associated microglia clusters to spatial deposition of Aβ. Using expression-based profiles, we identified 6 microglia clusters (clusters 0 to 5) within our dataset (Figure 3A, B and S4A). Homeostatic genes such as *TMEM119* were significantly enriched in cluster 0, whereas DAM-like genes such as *GPNMB* were significantly enriched in cluster 4 (Figures 3B-D, S4A and Table S5). DAM-like genes were generally upregulated while homeostatic genes were downregulated in AD vs control hippocampus, in most clusters (Figure 3C), with specific enrichment of DAM-like gene signatures in microglia cluster 4 (Figure 3D), indicating that microglia expression profiles are affected both generally by AD status and specifically by cell states as defined by the clusters. However, segregation of homeostatic and DAM-like markers appeared to be somewhat partial; DAM-like genes such as *TREM2*, *TYROBP* (*DAP12*) and *SPP1* were also expressed in cluster 0 microglia, and homeostatic genes such as *TMEM119* were likewise expressed in microglia cluster 4 (Figure 3B). Despite intermixed expression of some DAM-like and homeostatic genes, we observed modest enrichment of DAM-like (cluster 4) microglia in direct proximity to Aβ in AD hippocampus normalized to all microglia clusters within a distance interval (Figure 3E), and observed a more drastic enrichment of cluster 4 microglia in direct proximity (10 μm) of Aβ plaques as a percentage of cells in all distance intervals within a specific cluster (Figure 3F). Trending depletion of cluster 0 (potentially homeostatic) microglia was observed in close proximity to Aβ plaques (Figure 3E, F).

**Figure 3.**
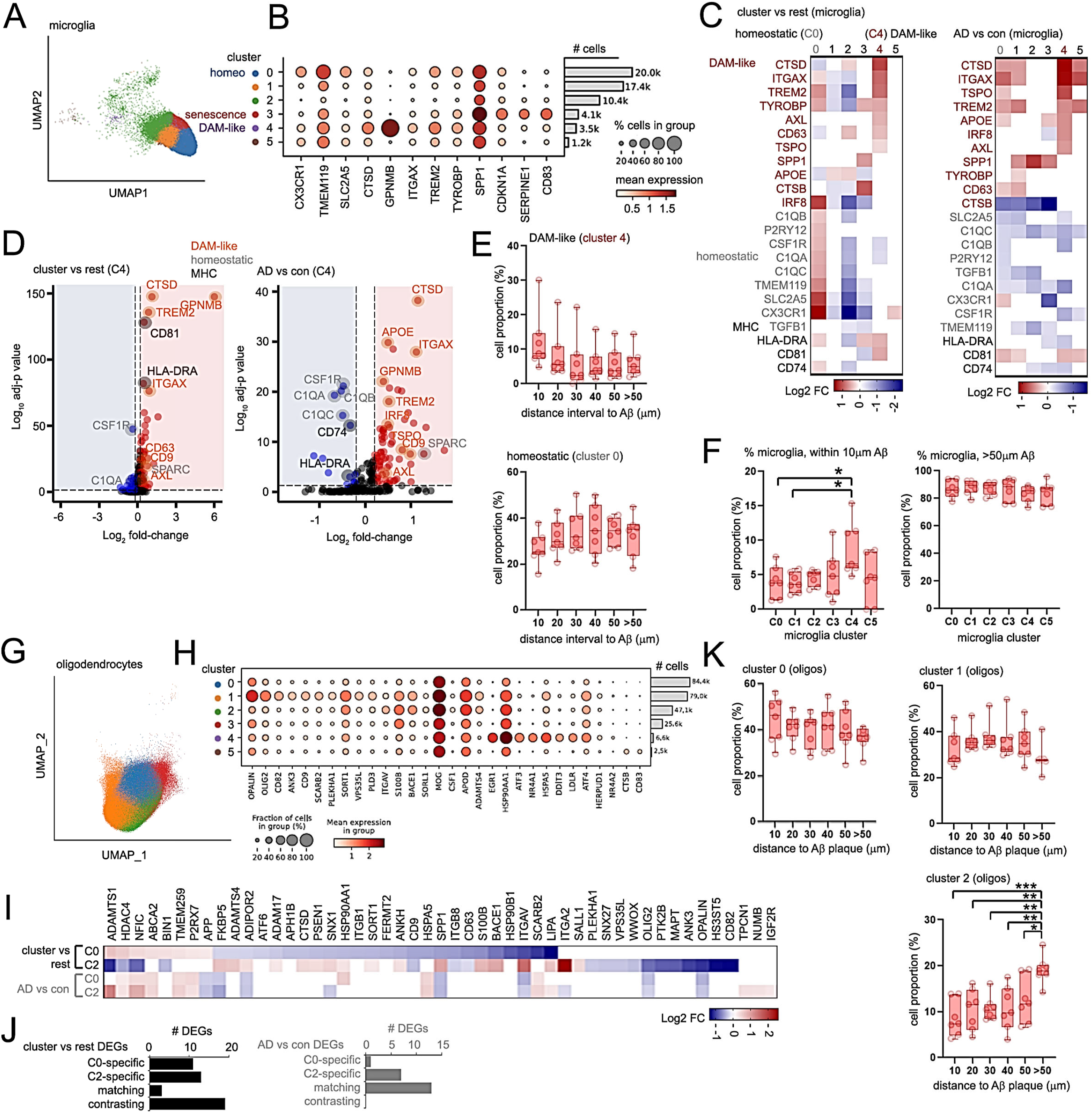
Subclustering and characterizing microglia and oligodendrocytes. (A) UMAP representation of 56,615 microglia in control and AD hippocampus. A total of 6 microglia subclusters (0 to 5) were identified. (B) Dot plot indicating fraction of cells (%) and level of expression of various microglia transcripts in clusters indicated. Percentage of cells expressing microglia transcripts (scaled dot size) or mean expression (red heat scale) are indicated. Bar graph indicates number of cells in each cluster. (C) Heatmaps depicting Log2 fold-change in microglia DAM-like (red), homeostatic (gray) or MHC-associated DEGs in microglia clusters 0-5 unique to each cluster (“cluster vs rest”, left heatmap) or AD vs control (right heatmap) comparisons. (D) Volcano plots for microglia cluster 4 enriched for DAM-like DEGs in “cluster vs rest” or “AD vs control” comparisons. Upregulated DAM-like (orange), homeostatic (gray) and MHC (black) genes are indicated (adj-p<0.05, absolute L2FC ≥ 0.2). (E) Box and whisker plots indicating percentage of microglia identified in different distance intervals surrounding Aβ plaques in cluster 4 (top) or cluster 0 (bottom) microglia. Percentages were calculated as a fraction of all microglia within a distance category from all clusters. (F) Box and whisker plots indicating percentage of microglia identified within 10 μm of Aβ-plaques (left) and > 50 μm away (right) in microglia clusters 0-5. Percentages were normalized to all microglia in all distance categories within a cluster designation. (G) UMAP subcluster representation of 245,101 oligodendrocytes in control and AD hippocampus. A total of 6 oligodendrocyte subclusters (0 to 5) were identified as represented by colors indicated in the legend. (H) Dot plot indicating fraction of cells (%) and level of expression of various oligodendrocyte transcripts in clusters indicated. Percentage of cells expressing oligodendrocyte transcripts (scaled dot size) or mean expression (red heat scale) are indicated. Bar graph indicates number of cells identified in each cluster. (I, J) (I) Heatmap depicting Log2 fold-change in cluster-specific (cluster vs rest) or AD vs control DEGs in C0 or C2 oligodendrocyte clusters. (J) Graphs indicating number of C0 or C2-specific, or matching/contrasting cluster vs rest (left) or AD vs con oligodendrocyte DEGs (right). (K) Box and whisker plots depicting percentage proportion of cells within different distance intervals to Aβ plaques in AD for oligodendrocyte clusters 0 to 2 as indicated. Percentages were calculated from a total number of cells from all clusters within a distance interval. Statistical significance in (E), (F) and (K) was determined by one-way ANOVA (*p<0.05, **p<0.01, ***p<0.001).

Astrocytes were clustered into 7 clusters (clusters 0 to 6) based on expression profiles (Figure S4B-D). We observed enrichment of reactive astrocyte genes such as *CD44* and *SERPINA3* in cluster 0 (Figures S4D-F and Table S5). Cluster 3 astrocytes featured marked expression of several transcription factors, including *NR4A1*, *NR4A2*, *EGR1* and *ATF3*, and cluster 5 showed enrichment of genes such as *CD83*, *C5AR1* and *DOC2A* (Figure S4C). While we observed enrichment of genes related to astrocyte activation such as *CD44*, *SERPINA3*, *OSMR* and *VIM* in astrocyte cluster 0 (Figure S4E, F), we did not see enrichment of cluster 0 or other clusters in proximity to Aβ plaques (Figure S4G).

Oligodendrocytes, excitatory and inhibitory neurons were also clustered according to transcriptional profiles and annotated for proximity to Aβ plaques. Oligodendrocytes were clustered into 6 transcriptional subclusters (clusters 0 to 5) (Figure 3G, H and S4H, Table S5). Clusters 0 and 2 featured opposing DEG profiles (Figure S4D). A comparison of cluster-specific (“cluster vs rest”) DEG profiles of clusters 0 and 2 revealed a high number of contrasting DEGs between these two clusters (Figure 3I, J). Interestingly, cluster 0 showed a trending enrichment, while cluster 2 was significantly depleted in proximity to Aβ plaques (Figure 3K). Excitatory neurons were clustered into 6 transcriptional clusters (clusters 0 to 5) (Figure S5A, B) and inhibitory neurons were clustered into 4 subclusters (clusters 0 to 3) (Figure S5C, D). For both excitatory and inhibitory neurons, clusters 0 and 1 showed distinct, opposing DEG expression profiles (Figure S5B, D). These results indicate that transcriptional clustering using our MERSCOPE probe panel can segregate expression profiles in various cell subtypes, and while microglia subcluster 4 and oligodendrocyte subcluster 0 show trending enrichment in proximity to Aβ plaques, transcription-based subclusters in other cell types do not show any spatial association with plaques.

We therefore applied a direct approach to define expression profiles associated with Aβ plaques by characterizing cell expression profiles within 50 μm (Aβ+ cells) and outside 50 μm (Aβ-cells) distance interval from plaque boundaries (Figure 4A). Using this method, we characterized microglial Aβ+ vs Aβ-DEGs in AD and control hippocampus (Figure 4B and Table S6), or AD vs control DEGs in Aβ-proximal (Aβ+) or Aβ-distal (Aβ-) microglia (Table S6). We observed enrichment of some DAM-like genes such as *CTSD*, *SPP1* and *APOE* in Aβ-proximal microglia, and downregulation of some homeostatic genes such as *CX3CR1* and *SLC2A5* (Figure 4C); while *SLC2A5* expression was reduced in all or AD Aβ+ microglia, its expression was conversely upregulated in control Aβ+ microglia (Figure 4C). Moreover, some Aβ-associated microglia genes such as *C1QB* and *CSF1R* were upregulated in AD, yet downregulated in control microglia (Figure 4C).

**Figure 4.**
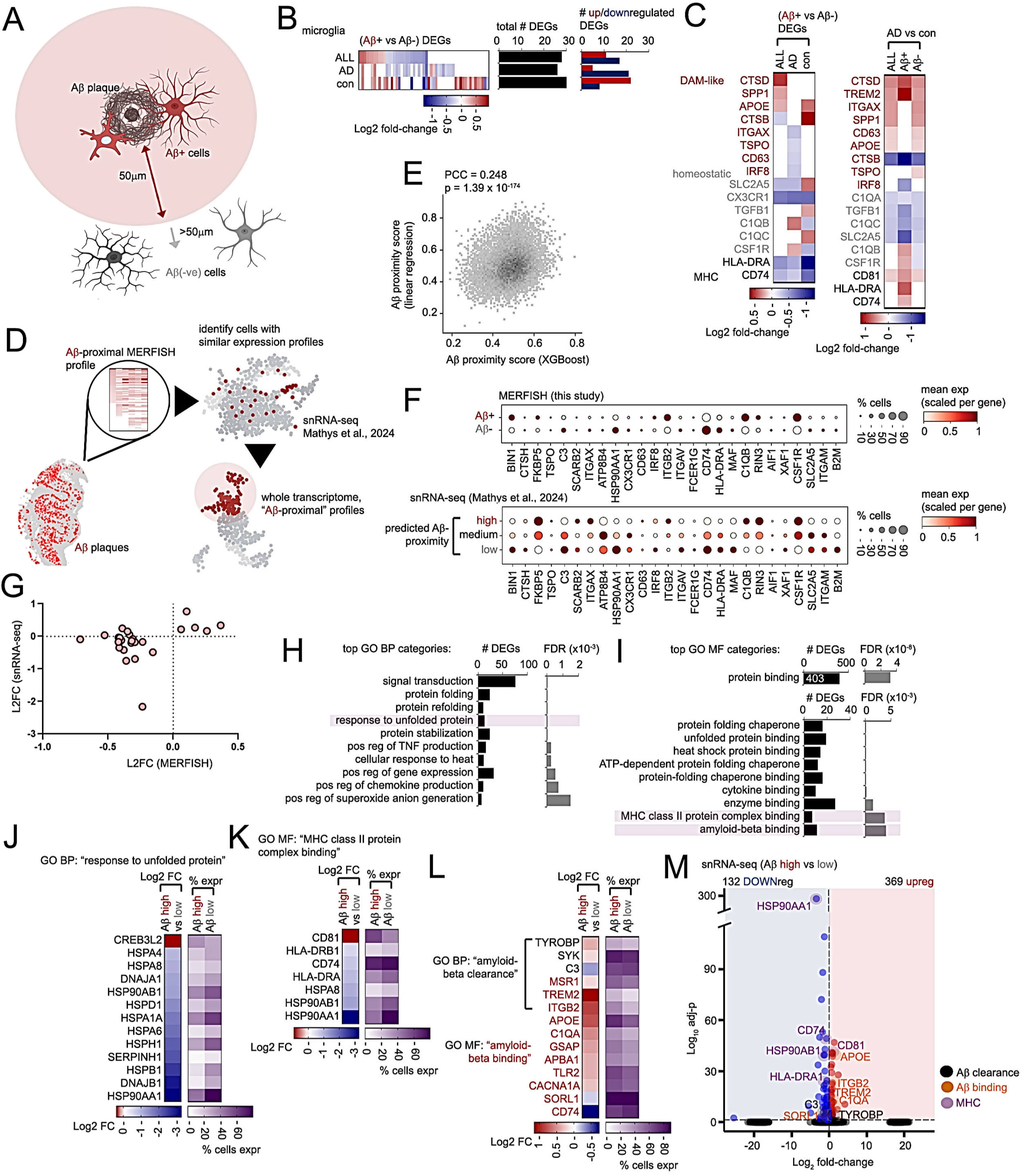
Characterizing expression profiles in Aβ plaque-proximal microglia and oligodendrocytes. (A) Schematic depicting annotation of Aβ plaque-associated (Aβ+) and Aβ-plaque distal (Aβ-) cell types. Aβ+ cells reside within 50 μm, while Aβ-cells fall outside 50 μm of Aβ plaque boundaries. (B) Heatmap depicting Aβ-plaque proximal (Aβ+ vs Aβ-) DEGs identified in all, AD or control microglia. Adjacent graphs show total number of DEGs (black bars) or up (red) and down-regulated (blue) DEGs. (C) Heatmaps depicting Log2 fold-change in microglial DAM-like, homeostatic or MHC DEGs from all, AD or control Aβ+ vs Aβ-microglia comparisons (left) or AD vs con DEGs in all, Aβ+ or Aβ-microglia (right). (D) Schematic depicting use of pathology-proximity-defined MERFISH cells to predict pathology proximal/distal profiles in snRNA-seq datasets. Pathology-associated MERFISH gene expression profiles are used to identify/predict cells with similar profiles in snRNA-seq datasets (Mathys et al., 2024); whole transcriptome profiles of “pathology-proximal” cells from snRNA-seq datasets are then characterized. (E) Scatterplot depicting Aβ-proximal scores for microglia in snRNA-seq data predicted by linear regression (LR) (y-axis) and XGBoost (x-axis) methods. Pearson’s correlation coefficient (PCC) and p-value are shown. (F) Dot plots depicting mean and percentage expression of genes enriched or depleted in Aβ-proximal (Aβ+) or distal microglia (Aβ-) identified by MERFISH analysis (top dot plot), with similar expression profiles in snRNA-seq datasets with high, medium or low Aβ pathology (Mathys et al., 2024; bottom dot plot). (G) Scatterplot showing Log2 fold-change, high vs low pathology in snRNA-seq (Mathys et al., 2024; y-axis) or Aβ+ vs Aβ-MERFISH comparisons (x-axis). (H, I) Top GO BP (H) and MF (I) categories identified for Aβ-proximal microglia DEGs mapped in snRNA-seq datasets. (J-L) Heatmaps depicting Log2 fold-change (left heatmap) and percent expression (right heatmap) in Aβ proximal and distal (Aβ high and low) microglia of DEGs (Aβ high vs Aβ low) in GO BP “response to unfolded protein” (J), GO MF “MHC class II protein complex binding” (K) and GO BP “amyloid-beta clearance” and “amyloid-beta binding” categories (L). (M) Volcano plot of mapped Aβ-proximal vs distal (high vs low) DEGs in microglia from snRNA-seq datasets (Mathys et al., 2024). DEGs related to Aβ clearance (black), Aβ-binding (orange) and MHC (purple) are highlighted.

We then aligned our Aβ-associated microglia MERSCOPE expression profiles with microglia clusters identified using snRNA-seq datasets from AD and control human brain (Figure S6A) (Gerrits, Brouwer et al. 2021). We observed some alignment of MERSCOPE Aβ+ microglia DEGs with phagocytic/activated “AD1” snRNA-seq microglia clusters (Figure S6B, D), as well as “AD2” homeostatic snRNA-seq clusters (Figure S6C, D). Interestingly, we also found that AD2 snRNA-seq microglia featured contrasting expression profiles with Aβ+ microglia in both AD and control brain, with higher contrast in control brain (Figure S6C, D).

We compared astrocyte Aβ+ vs Aβ-DEGs (Figure S6E) with DEGs from comparing snRNA-seq astrocyte profiles among individuals with no, early or late AD pathology in prefrontal cortex (Brodmann area 10) (Mathys, Davila-Velderrain et al. 2019), and revealed some overlap in DEGs (Figure S6F, G). In comparing Aβ+ oligodendrocyte profiles (Figure 4H) to early/late AD oligodendrocyte profiles from snRNA-seq datasets, we find very good alignment of Aβ+ MERSCOPE oligodendrocyte DEGs enriched or depleted in both early and late AD pathology (Figure S6I, J), especially in DEGs depleted in AD pathology such as *CD9* and *SPP1* (Figure 4I).

Having established expression profiles in Aβ-proximal microglia, we used Aβ-associated expression templates to characterize potential Aβ-proximal profiles in snRNA-seq datasets. To this end, we trained machine learning models using MERFISH-based Aβ-proximal/distal expression profiles in microglia to identify microglia of similar profiles in snRNA-seq datasets from human control and AD hippocampus (Figure 4D-G) (Mathys, Boix et al. 2024). This approach allowed us to find potential Aβ-proximal/distal cells in snRNA-seq data even if they do not form cell clusters naturally. Whole-transcriptome coverage of snRNA-seq then allowed us to compare Aβ-proximal and Aβ-distal cells for genes not profiled by our MERFISH panel. We characterized DEG expression profiles in predicted “Aβ-proximal” compared to “Aβ-distal” microglia from snRNA-seq data (Table S7, Figure 4H-M), and identified a total of 501 DEGs, which showed enrichment in GO Biological Process (BP) categories such as “response to unfolded protein” and GO MF categories such as “MHC class II protein complex binding” and “amyloid-beta binding” (Figure 4H-M). We also observed enrichment of GO KEGG pathways related to “cellular senescence” (Figure S7A, B) and GO BP “positive regulation of phagocytosis” and GO MF “cytokine binding” categories (Figure S7C, D). We also observed upregulation of DAM and some homeostatic DEGs in Aβ-associated microglia snRNA-seq profiles, with marked upregulation of *GPNMB* (Figure S7E). Similarly, GPNMB was found as the key marker of a GPNMB+ microglia population associated with lipid processing, cell migration and vascular reorganization from a snRNA-seq study (Luquez, Algoo et al. 2026).

Similarly, we used machine learning models to predict potential Aβ-proximal/distal excitatory neurons in snRNA-seq datasets (Table S9, Figure S8A-C), and observed a total of 2,960 DEGs in predicted Aβ-proximal vs Aβ-distal excitatory neurons (Figure S8D, E) (Mathys, Boix et al. 2024). Aβ-associated DEGs were enriched in KEGG pathways related to “glutamatergic synapse”, “GABAergic synapse”, and “Alzheimer’s disease” (Figure S8F-I), as well as in the GO MF terms “amyloid-beta binding” and “tau protein binding” (Figure S8J). We observed downregulation of Aβ-binding DEGs such as *LDLR* and *SORL1* in Aβ-proximal excitatory neurons (Figure S10J), as well as strong downregulation of tau-binding DEGs with an exception that *APOE* was upregulated in Aβ-proximal excitatory neurons (Figure S10J). In addition, upregulated DEGs in predicted Aβ-proximal excitatory neurons were enriched for the GO BP terms “positive regulation of amyloid-beta formation” including *GSAP* and *APOE* (Figure S8K). In contrast, downregulated DEGs were enriched in “negative regulation of amyloid-beta formation” including *PIN1* and *SORL1*, as well as “amyloid-beta clearance” including *CD36* and *LDLR* (Figure S8L).

### Cell-type-specific expression profiles in association with AT8 ptau pathology

Given that tau pathology is an intracellular feature enriched primarily in neurons (Luo, Yu et al. 2026), we quantified AT8 ptau (pS202/T205) staining signals in areas covering neuron cell bodies as ptau levels for excitatory and inhibitory neurons. Based on the quantified staining scores, we further identified AT8+ excitatory (Figure 5A, Table S7) and inhibitory neurons (Figure S9A, Table S8) in AD and control hippocampus, and also neurons with little/no AT8 ptau (AT8-) staining (Figure 5B, C). Non-neuronal cells were classified into AT8+ or AT8-if proximal or distal to AT8+ neurons respectively (Figure 5B). We then mapped AT8+ DEG profiles to cell-specific transcriptional clusters in neurons and glia (Figure 5B).

**Figure 5.**
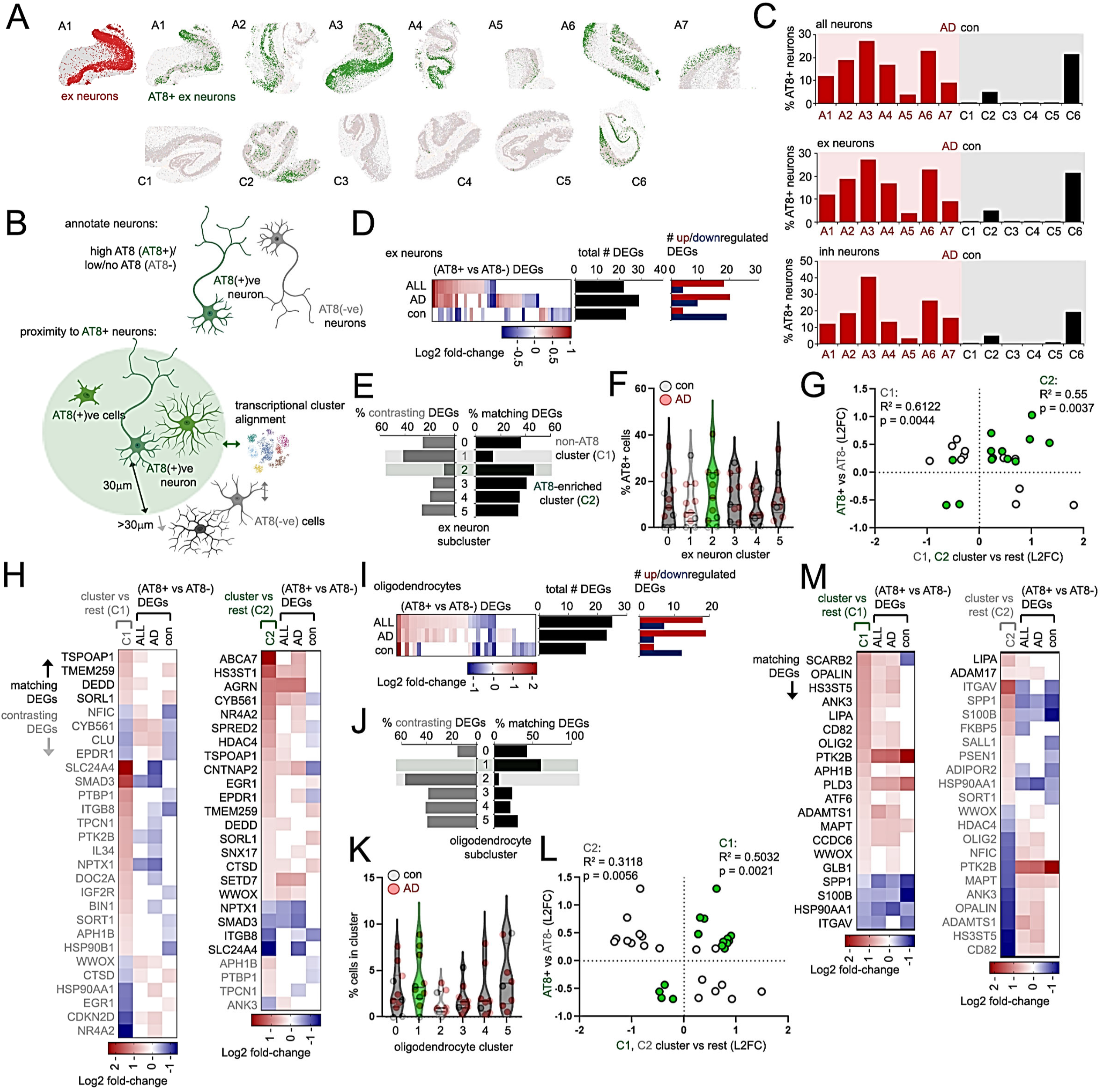
Characterizing tau-associated expression profiles in neurons and oligodendrocytes. (A-C) Annotation of AT8+ excitatory (“ex”) neurons in AD and control hippocampus. (A) AT8-enriched excitatory neurons (green) identified in AD (A1-A7) or control (C1-C6) hippocampus. (B) Schematic, annotation of AT8+ neurons and (AT8+)-proximal cell types. Neurons identified with high AT8 levels are AT8+, while neurons with little or no AT8 ptau are AT8-(top panel). For non-neuronal cell types, cells with AT8+ neurons within 30 μm are AT8+, while cells with neither high nor medium-level neurons within 30 μm are AT8-. To determine whether transcriptional cell clusters for various cell types are enriched for AT8+ (neurons) or proximity to AT8+ neurons, transcriptional cluster profiles are aligned to AT8+ vs AT8-DEGs. (C) Graphs depicting percentage of AT8+ neurons in all, excitatory (“ex”) or inhibitory (“inh”) neurons in AD (red bars) or control hippocampus (black bars). (D) Heatmap showing Log2 fold-change of DEGs in all, AD or control AT8+ vs AT8-excitatory neurons. Adjacent graphs depict number of total (black bars) and up (red) or down-regulated DEGs. (E-G) Alignment of AT8-enriched DEGs to transcriptional ex neuron clusters. (E) Graph depicting alignment of % matching (black bars) or contrasting (gray bars) excitatory neuron DEGs in clusters 0 through 5 in comparison to AT8+ vs AT8-DEGs. Cluster 1 features a high number of contrasting and reduced number of overlapping AT8+ DEGs (potential AT8-depleted cluster), while cluster 2 shows good alignment and reduced number of contrasting AT8+ vs AT8-DEGs (potential AT8-enriched cluster). (F) Percentage of AT8+ excitatory neurons in each ex neuron cluster; potential AT8-depleted (light gray) and AT8-enriched clusters (green) are indicated. Plots representing percentages for ex neurons in each AD (red) and control (gray) hippocampus are shown. (G) Scatterplot depicting Log2 fold-change (L2FC) in ex neuron cluster 1 (C1, gray) and C2 (green) DEGs; cluster vs rest (x-axis) and AT8+ vs AT8-(“ALL” DEGs, y-axis) L2FC values for individual DEGs, and R^2^/p-values for C1 and C2 trends are shown. (H) Heatmap showing Log2 fold-change profiles of ex neuron cluster 1 and 2 profiles aligned with AT8+ vs AT8-ex neuron DEGs. (I) Heatmap showing Log2 fold-change of DEGs in all, AD or control AT8+ vs AT8-oligodendrocytes. Adjacent graphs depict number of total (black bars) and up (red) or down-regulated DEGs. (J-M) Alignment of AT8-enriched DEGs to transcriptional oligodendrocytes clusters. (J) Graph depicting alignment of % matching (black bars) or contrasting (gray bars) oligodendrocyte DEGs in clusters 0 through 5 in comparison to AT8+ vs AT8-DEGs. Cluster 1 features enrichment of overlapping, and reduced contrasting AT8+ vs AT8-DEGs (potential AT8-enriched cluster), while cluster 2 shows a high number of contrasting/reduced number of overlapping AT8+ oligodendrocyte DEGs (potential AT8-depleted cluster). (K) Percentage of AT8+ oligodendrocytes in each cluster; potential AT8-enriched clusters (green) and AT8-depleted (light gray) are indicated. Plots representing percentages for oligodendrocytes in individual AD (red) and control (gray) hippocampal sections are shown. (L) Scatterplot depicting Log2 fold-change (L2FC) in oligodendrocyte cluster 1 (C1, green) and C2 (gray) DEGs; cluster vs rest (x-axis) and AT8+ vs AT8-(“ALL” DEGs, y-axis) L2FC values for individual DEGs, and R^2^/p-values for C1 and C2 trends are shown. (K) Heatmap showing Log2 fold-change profiles of oligodendrocyte cluster 1 and 2 profiles aligned with AT8+ vs AT8-oligodendrocyte DEGs. Statistical significance in (G) and (L) were determined by simple linear regression analysis using Prism.

We defined AT8+ DEG profiles in all, AD or control excitatory neurons (Figure 5D) and inhibitory neurons (Figure S9B), and aligned AT8-associated DEGs with transcriptionally-defined excitatory (Figure 5E-G) or inhibitory neuron clusters (Figure S9C-E). Interestingly, we observed a high number of contrasting DEGs and low number of matching AT8+ DEGs in excitatory neuron cluster 1 (i.e., higher in C1 as compared to other clusters but lower in AT8+ as compared to AT8-, or vice versa) suggesting that this may represent a AT8-depleted cluster (Figure 5E-H); conversely, excitatory neuron cluster 2 featured a high number of matching and low number of contrasting AT8+ DEGs (Figure 5E-H), implicating cluster 2 as an AT8-enriched excitatory neuron cluster. In quantifying percentage of AT8+ cells within each excitatory neuron cluster, we indeed observed depletion of AT8+ neurons in cluster 1, and enrichment of AT8+ neurons in cluster 2 (Figure 5F). These results suggest that cells in these two clusters have widespread expression changes due to absence/presence of AT8, causing them to form clusters separated from other excitatory neurons. We also observed enrichment of contrasting and depleted number of matching AT8+ DEGs in inhibitory neuron cluster 2 (Figure S9C-F), as well as a reduced percentage of AT8+ neurons in cluster 2 (Figure S9D), suggesting that cluster 2 represents an AT8-depleted cluster.

Transcriptional-based clusters were aligned to glia proximal (AT8+) and distal (AT8-) to AT8-enriched neurons. Although we observed a high number of matching/reduced number of contrasting AT8+ DEGs in microglia cluster 4 (Figure S9G, H), we did not observe an increased proportion of AT8+ microglia in cluster 4 (Figure S9I). However, we observed increased expression of DAM-like genes in AT8+ microglia (Figure S9J). Homeostatic microglia genes with the exception of *C1QA* were not downregulated in AT8+ microglia; however, we observed reduced levels of *C1QA*, *C1QC*, *SLC2A5* in AT8-, reductions of *C1QB*, *CSF1R* in AT8+ and *TGFB1* in both AT8+ and AT8-microglia in AD vs control comparisons (Figure S9J). Astrocyte cluster 0 was identified as a potential AT8-depleted cluster, while clusters 1 and 4 were potential AT8+ astrocyte clusters (Figure S9K-O). In agreement with this, the percentage of AT8+ was reduced in astrocyte cluster 0, and somewhat increased in astrocyte cluster 4 (Figure S9M). Oligodendrocyte cluster 1 was identified as a potential AT8-enriched, while cluster 2 appeared to be a potential AT8-depleted oligodendrocyte cluster (Figure 5I-M). Although cluster 1 only showed slight enrichment in the percentage of AT8+ oligodendrocytes, oligodendrocyte cluster 2 showed a reduced number of AT8+ oligodendrocytes (Figure 5K). Overall, these results indicate that cell-type-specific transcriptional clusters such as excitatory neuron cluster 2 (AT8-enriched), and inhibitory neuron cluster 2, astrocyte cluster 0 and oligodendrocyte cluster 2 (AT8-depleted) are spatially responsive to AT8 pathology.

To align AT8-enriched MERFISH profiles to cells from AD and control human hippocampus snRNA-seq datasets for excitatory neurons, we trained machine learning models using MERIFSH data and predicted potential AT8+ and AT8-cells in the snRNA-seq data (Table S9, Figure S10A-C) (Mathys, Boix et al. 2024). Predicted AT8+/-excitatory neurons showed good expression alignment with MERFISH profiles (Figure S10B, C), and produced total of 3,620 significant AT8+ vs AT8-DEGs in excitatory neurons (Figure S10D). DEGs were enriched in KEGG pathways such as “ABC transporters” and “glutamatergic synapse” pathways (Figure S10E-G). In comparing snRNA-seq DEG profiles from predicted Aβ+ or AT8+ excitatory neurons, we observed a high degree of overlap between Aβ+ and AT8+ profiles (Figure S10H). Interestingly, DEGs such as *HSP90AA1*, *NEFL* (neurofilament) and *CDKN2D* (p19^INK4d^) were depleted in Aβ+, yet enriched in AT8+ excitatory neurons (Figure S10H), indicating that corresponding components may be specific to Aβ-proximity or AT8 ptau enrichment. GO MF “tau-binding” and GO BP: “negative regulation of amyloid-beta formation” DEGs such as *APOE* were upregulated in both Aβ+ and AT8+ excitatory neurons, while *CLU* (ApoJ) was specifically upregulated in AT8+ neurons (Figure S10J, K). We also observed 32 DEGs related to GO MF “amyloid-beta binding” in Aβ+ and AT8+ excitatory neurons (Figure S10L). Together, this suggests that Aβ+ and AT8+ profiles show a high degree of overlap in excitatory neurons.

### Characterizing cell-type-specific profiles related to convergence of Aβ and AT8 ptau pathology

We next characterized regional expression profiles in cells spatially associated with both Aβ pathology and AT8+ neurons. Based on Aβ and Tau pathological annotation of MERFISH cells, we performed systematic comparisons including AT8+ vs AT8-when proximal (Aβ+) or distal to Aβ (Aβ-), Aβ+ vs Aβ-when Tau annotation is AT8+ or AT8-, and Aβ+/AT8+ vs Aβ-/AT8-(Figure 6A-C, Table S10). Our prior analysis showed that DAM-like microglia cluster 4 is enriched in close proximity to Aβ (Figure 3E, F), and has little association with AT8+ tau (Figure S9H). In further categorization based on combined proximity to Aβ and AT8 Tau, we also found that cluster 4 was nominally significantly enriched specifically in Aβ+/AT8-category (*p*=0.02, Wilcoxon rank-sum test comparing the proportions of cluster 4 cells in Aβ+/AT8-category with the overall proportions of cluster 4 cells among microglia across AD samples). This suggests that cluster 4 microglia may be preferentially associated with Aβ pathology in the absence of AT8 Tau pathology.

**Figure 6.**
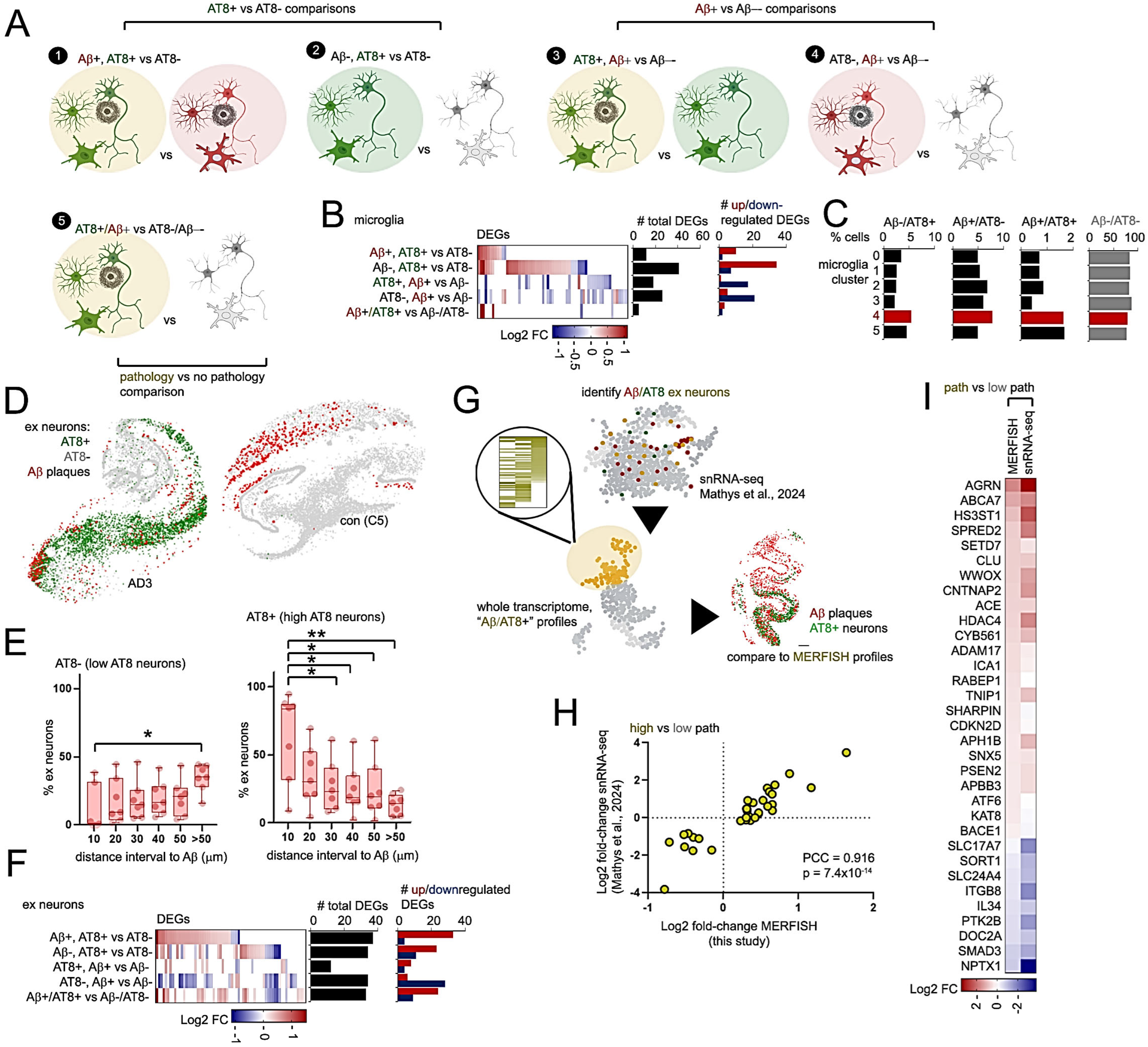
Characterizing cell-specific expression profiles associated with combined Aβ plaques and AT8 ptau pathology. (A) Cell-specific MERSCOPE AT8+ vs AT8-comparisons with or without Aβ (“1”, “2”), Aβ+ vs Aβ-comparisons with or without AT8 pathology (“3”, “4”), or with combined Aβ/AT8 pathology vs no pathology (“5”). (B) Microglia DEGs from the five AT8+ vs AT8- and Aβ+ vs Aβ-comparisons described in (A). Adjacent graphs depict number of total (black bars) and up (red) or down-regulated microglial DEGs. (C) Graphs depicting percentage of Aβ-/AT8+, Aβ+/AT8-, Aβ+/AT8+ or Aβ-/AT8-microglia in clusters 0 through 5, normalized as a percentage of total cells from each cluster. Microglia “DAM-like” cluster 4 is highlighted in red. (D-H) Characterization of Aβ and AT8-associated profiles in ex neurons. (D) Representative hippocampal tissue sections from AD (AD3) and control (C5) individuals. Aβ plaques (red), AT8+ (green) and AT8-excitatory neurons (gray) are indicated within hippocampal sections. (E) Graphs representing percentages of AT8-(left) and AT8+ excitatory neurons (right graph) in different distance intervals to Aβ plaques. (F) Heatmap depicting Log2 fold-change in ex neuron DEGs from the five AT8+ vs AT8- and Aβ+ vs Aβ-comparisons described in (A). Adjacent graphs depict number of total (black bars) and up (red) or down-regulated (blue) ex neuron DEGs. (G) Schematic: DEG profiles from excitatory neurons with high predicted Aβ and AT8 pathology from snRNA-seq analysis (Mathys et al., 2024) were compared to expression profiles obtained by MERFISH analysis. (H) Scatterplot showing Log2 fold-change, high vs low pathology in snRNA-seq (Mathys et al., 2024; y-axis) or Aβ+/AT8+ vs Aβ-/AT8-MERFISH comparisons (x-axis). Pearson’s correlation co-efficient (PCC) and p-value is shown. (I) Heatmap depicting Log2 fold-change in ex neuron DEGs from MERFISH and snRNA-seq datasets shown in (H). Statistical significance in (E) was determined by one-way ANOVA with Tukey’s multiple comparisons (*p<0.05, **p<0.01).

We observed convergence of Aβ plaques and AT8+ excitatory neurons (Figure 6D, E) and inhibitory neurons (Figure S11A, B) in AD hippocampus (Figure 6D, E and S11A, B). More specifically, we observed some enrichment of AT8-excitatory neurons in regions distal to Aβ plaques, as well as an enrichment of AT8+ excitatory and inhibitory neurons within a 10 μm distance-interval of Aβ plaques in AD hippocampus (Figure 6E and S11B). In comparison to microglia, we observed a greater number of Aβ+/AT8+ vs Aβ-/AT8-DEGs in excitatory neurons (Figure 6F). In contrast to excitatory neurons, inhibitory neurons featured fewer DEGs in combined Aβ/AT8 ptau comparisons; no significant DEGs were found for AT8-, Aβ+ vs Aβ-comparisons (Figure S11C). Based on predicted Aβ+/- and AT8+/-excitatory neurons from snRNA-seq datasets (Mathys, Boix et al. 2024), we identified cells predicted to be Aβ+/AT8+ or Aβ-/AT8-in combination, and compared DEG profiles of Aβ+/AT8+ vs Aβ-/AT8-excitatory neurons between snRNA-seq and MERFISH datasets (Table S9, Figure 6G). We observed a high correlation in gene expression changes from Aβ+/AT8+ vs Aβ-/AT8-between snRNA-seq and MERFISH datasets (Figure 6H, I), indicating a good convergence of combined Aβ/AT8 profiles in snRNA-seq and MERFISH analyses. Comparing among Aβ+ (Aβ+ vs Aβ-), AT8+ (AT8+ vs AT8-) and combined Aβ+/AT8+ (Aβ+/AT8+ vs Aβ-/AT8-) DEGs from snRNA-seq excitatory neurons, we observed a high degree of overlap, and also DEGs unique to Aβ+, AT8+ and Aβ+/AT8+ excitatory neurons in combination (Figure S11D, E). DEGs unique to Aβ-proximal excitatory neurons include downregulation of genes related to KEGG “oxidative phosphorylation” and “ribosome” pathways (Figure S11F, G), while DEGs unique to AT8+ neurons showed enrichment in GO BP “positive regulation of myelination” and “positive regulation of type II interferon production” categories (Figure S11H, I). DEGs unique to Aβ+/AT8+ excitatory neurons showed enrichment in GO BP “fatty acid biosynthetic process” with predominant upregulation of DEGs in this category (Figure S11J, K).

### Identifying potential cis-eQTLs associated with spatial proximity to AD pathology

By integrating spatial transcriptomics, pathology staining and genetic data, we performed cis-eQTL analysis coupled to Aβ-proximity to identify cell-type/genotype-specific transcriptional changes within local pathological microenvironments in human AD hippocampus (Figure 7A). From whole genome sequencing data, we called 56,311 cis-variants with primarily single nucleotide polymorphisms (SNPs) within 10 kb upstream and downstream of gene body regions of the 120 AD risk genes. We also validated allele-associated, pathology-dependent eQTL expression features using coupled genotype (De Jager, Ma et al. 2018) and snRNA-seq expression profiles (Fujita, Gao et al. 2024) from external AD studies including 424 individuals stratified by overall Aβ and tau pathology burden (“Fujita et al.”) to mirror Aβ/AT8 ptau annotations in our MERFISH analysis. Overall, major allele frequency profiles were comparable between our cohort and the Fujita et al. snRNA-seq dataset (Fujita, Gao et al. 2024) (Figure 7B). A significant number of eQTL-associated genes (eGenes) when being proximal to (Aβ+) or distal from (Aβ-) Aβ pathology was observed between our cohort and the external cohort across glial and neuronal cell types (Figure 7C). We then identified eQTLs specific to Aβ+ or Aβ-microenvironments with overlapping cis-eQTL expression profiles in MERFISH and snRNA-seq cohorts (Figure 7D-G, Table S11).

**Figure 7.**
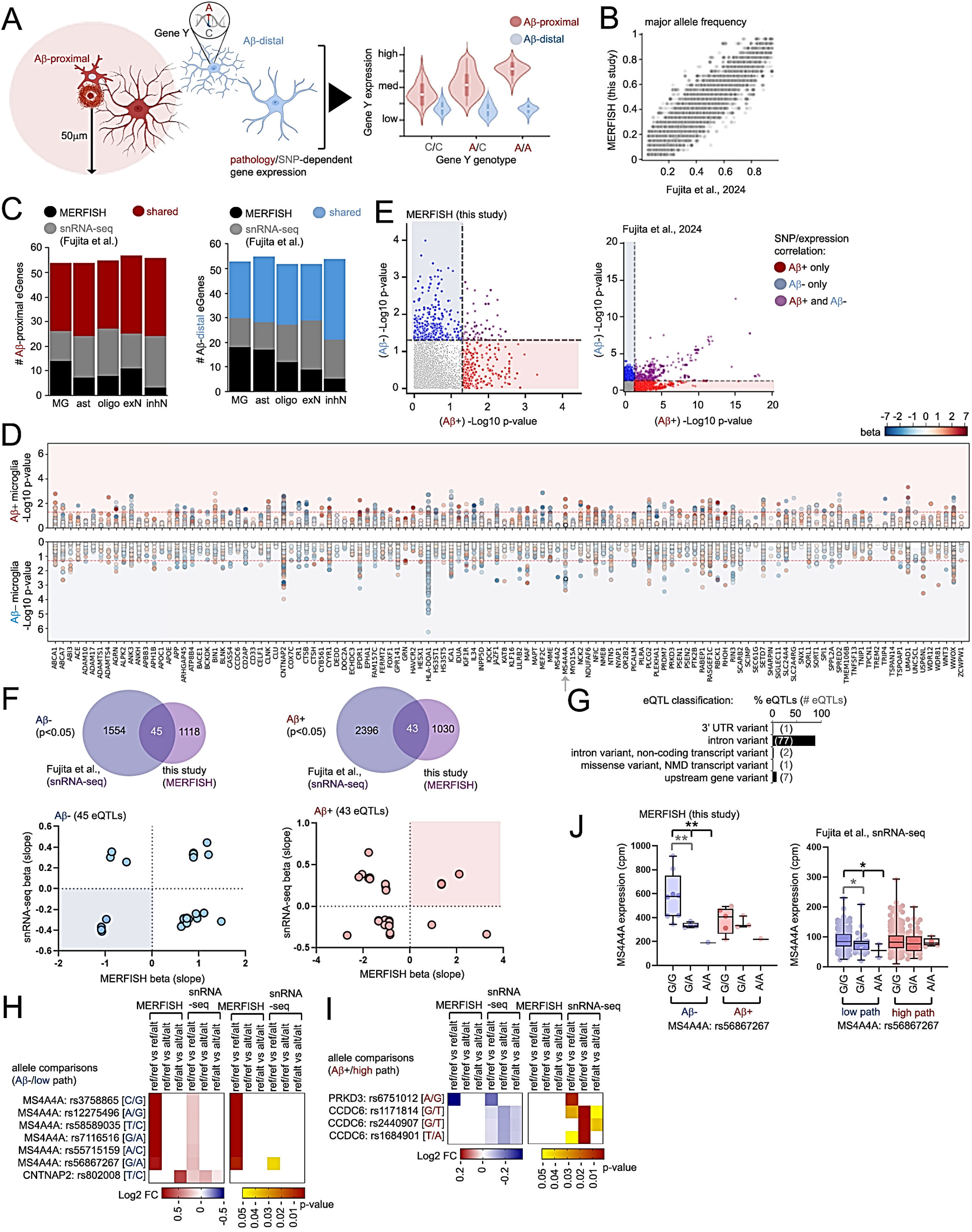
Use of spatial profiling to identify potential microglial SNP variants responsive to spatial proximity to Aβ pathology. (A) Schematic of pathology-dependent eQTL analysis: Gene expression profiles of various cell types in proximity (red) or distal to Aβ pathology (blue) are characterized according to gene variant within a gene locus. Potential gene variants responsive or refractory to Aβ are then identified. (B) Scatterplot comparing major allele frequency of variants called in the MERSCOPE cohort (this study) compared to a published cohort of 424 aged individuals (Fujita et al., 2024). (C) Number of shared eGenes derived from Aβ-proximal (Aβ+, top) or Aβ-distal cells (Aβ-, bottom graph) identified in MERFISH and snRNA-seq datasets (highlighted in red and blue). Unique eGenes for MERSCOPE and snRNA-seq datasets are shown in black and gray. (D) Manhattan plot of Aβ+ (top) or Aβ-eQTLs (bottom plots) results in microglia, -log10(p-value) and heat-scaled color indicators for beta (slope) are shown. (E) Scatterplot showing -log10(p-value) of eQTLs responsive in either Aβ- or Aβ+ spatial categories in MERFISH and snRNA-seq datasets (Fujita et al.). Non-significant hits are shown in gray. Aβ+ (red) or Aβ-(blue) or Aβ+/Aβ-responsive eQTLs (purple) are indicated. (F-I) Pathology-dependent eQTL prioritization and characterization of allele-specific microglia expression profiles in proximity/distal to Aβ plaques. (F) Venn diagrams (top panels) depicting overlapping, statistically significant eQTLs identified by snRNA-seq (Fujita et al.) and MERFISH analysis in Aβ-(left) and Aβ+ (right) microglia. Scatterplot depicting beta (slope) values in snRNA-seq datasets (Fujita et al., y-axis) and MERFISH analysis (this study). Blue and red highlights indicate eQTLs with negative or positive slope values in both datasets in Aβ-and Aβ+ microglia. (G) Graph depicting percentage (bars) and number of eQTLs (numbers in brackets) with different effects. (H, I) Heatmaps depicting Log2 fold-change (left heatmaps) and p-values (right heatmaps) for reference (“ref”) and alternative (“alt”) allele combinations comparisons in MERFISH and snRNA-seq (Fujita et al.) datasets in Aβ- or low pathology (H) or Aβ+/high pathology (I) microglia as indicated. (J) Boxplots showing *MS4A4A* expression in microglia from individuals comprising G/G, G/A or A/A genotypes at MS4A4A rs56867267 in MERFISH (left) and snRNA-seq datasets (right graph). Statistical significance in (J) was determined using a linear model in MatrixEQTL (see method section) and Wilcoxon rank-sum tests for pairwise comparisons between allele combinations. Black indicators represent statistical significance for eQTL analysis, and gray indicators depict statistical significant differences between allele combinations by pairwise analysis (*p<0.05, **p<0.01).

For microglia, we identified 45 and 43 eQTLs specific to Aβ+ or Aβ-microenvironments in both cohorts respectively (Figure 7D-F, Table S11), with most of the eQTLs involving intronic variants (Figure 7G). Among them, 6 eQTLs are associated with *MS4A4A* (Figure 7H-J and S12), with a particular enrichment of Aβ-responsive eQTLs residing between the *MS4A4A* promoter and enhancer region (Figure S12A, B). *MS4A4A* eQTL alleles such as rs56867267-G (Figure 7J) and rs3758865-C (Figure S12B) showed significant enhancement in *MS4A4A* expression in microglia distal to Aβ-plaques by MERFISH analysis, with similar trends observed in snRNA-seq datasets (Figure 7J, S12B) (Fujita, Gao et al. 2024), but with non-significant trends in microglia proximal to Aβ-plaques. Previously, a MS4A4A variant rs1582763 was found to enhance *MS4A4A* expression, and was associated with soluble TREM2 levels in AD patient CSF (Deming, Filipello et al. 2019). Conversely, MS4A4A deletion or inhibition was shown to reduce *TREM2* expression in iPSC-derived microglia (Rosner, Sun et al. 2026). In our cohort, we also observed a positive correlation between *MS4A4A* and *TREM2* expression in both control and AD microglia from human hippocampus (Figure S12D). We also observed eQTLs such as *CCDC6* rs1171814-T/T which featured increased *CCDC6* expression in Aβ-proximal microglia as compared to the G/T genotype; and although no pairwise significance between different rs1171814 genotypes within the MERFISH cohort, rs1171814-T/T showed significant effects in enhancing *CCDC6* expression in individuals with high Aβ pathology burden based on analysis of snRNA-seq data (Figure S12C). We also characterized other eQTLs that can potentially suppress gene expression in Aβ-distal (Aβ-) (Figure S13A-C) or enhance expression in Aβ-proximal (Aβ+) microglia (Figure S13D-F). For example, CNTNAP2 rs12540655-G/G suppressed *CNTNAP2* expression in Aβ-microglia in both MERFISH and snRNA-seq datasets, similar to RASGEF1C rs12516993-A/A (Figure S13C). Conversely, ANK3 rs7069191-C/C and CCDC6 rs1709345-T/T featured enhanced *ANK3* expression in Aβ+/high pathology microglia (Figure S13F).

For excitatory neurons, we identified and characterized cis-eQTLs that can potentially modulate expression in excitatory neurons proximal to Aβ plaques (Figures S14 and S15), as well as in AT8+ excitatory neurons (Figures S16 and S17). We identified eQTLs with significant effects in modulating expression at cis-loci in MERFISH (Figure S14A, B) and snRNA-seq datasets (Figure S14B), and characterized eQTLs with similar directional changes in both MERFISH and snRNA-seq analysis (Figure S14C, D). Pairwise comparisons of expression levels between different genotypes of the eQTL variants (Figure S14E and S15A, B) implicated eQTLs such as ACE rs4575595-G/G which enhanced ACE expression in Aβ-distal (MERFISH) or low pathology (snRNA-seq) excitatory neurons with similar trends in Aβ+ neurons (Figure S14F). Conversely, HS3ST5 rs13217813-G/G suppressed *HS3ST5* expression in Aβ-distal/low pathology neurons (Figure S14F). Additional Aβ-responsive eQTLs include RASGEF1C rs34351120-T/T which suppressed *RASGEF1C* expression, BACE1 rs2445939-T/T or MAPT rs63750222-C/C which enhanced *BACE1* expression in Aβ+/high pathology excitatory neurons (Figure S15C).

In addition to Aβ-microenvironment, we also characterized AT8 ptau-dependent cis-eQTLs for excitatory neurons (Figure S16A), and further prioritizated by overlapping with a parallel analysis using an external cohort with 424 individuals (Figure S16B-D and S17A). We identified eQTLs such as ACE rs4575595-G/G enhancing *ACE* expression in AT8-neurons, with similar trends in AT8+ neurons (Figure S17E, F); thus implicating effects of rs4575595-G in elevating *ACE* expression in both Aβ-(Figure S14F) and AT8-excitatory neurons (Figure S16F). Analysis of eQTLs that enhanced or suppressed cis-expression in AT8+ excitatory neurons (Figure S17A-D) implicated SNPs such as TNIP1 rs8177449-C/C and ZCWPW1 rs117078929-A/A which suppressed *TNIP1* and *ZCWPW1* expression in AT8+ excitatory neurons (Figure S17C), as well as ADAM17 rs12329030-G/G and TNIP1 rs3792789-G/G which enhanced *ADAM17* or *TNIP1* expression in AT8+ neurons (Figure S17E).

Together, these results describe a pathological-microenvironment-dependent cis-eQTL library comprising matching expression trends in both MERFISH and snRNA-seq cohorts (Fujita, Gao et al. 2024). Highlights include identification of a rs4575595-G that potentially increases *ACE* expression in both Aβ-and AT8-excitatory neurons, as well as MS4A4A rs56867267-G which increases *MS4A4A* expression in Aβ-distal microglia (and with weaker effects in Aβ-proximal microglia).

### APOE4/E4 expression is suppressed in astrocytes in proximity to Aβ-plaques

Similar to RNA-seq analysis in human brain (Zhang, Sloan et al. 2016), our MERSCOPE results indicate that *APOE* is most highly expressed in astrocytes, with some enrichment in microglia (Figure S18A). Having mapped and annotated astrocytes and microglia in proximity and distal to Aβ plaques, we then determined whether spatial distance to Aβ pathology can affect *APOE* expression in APOE2/E3, E3/E3, E3/E4 and E4/E4 carriers within our cohort (Figure 8A, B). We observed a slight reduction in *APOE* expression in astrocytes distal (> 50 μm) to Aβ plaques in APOE2/E3 and APOE3/E3 carriers (Figure 8C); conversely, we observed marked suppression of astrocyte *APOE* expression in proximity to Aβ plaques in a homozygous APOE4/E4 carrier within our cohort (Figure 8C). No variation in *APOE* expression was observed in APOE3/E4 astrocytes in proximity to Aβ (Figure 8C), suggesting that attenuation of *APOE* expression in Aβ-proximal astrocytes may be exclusive to homozygous APOE4/E4 carriers. We also observed slight reduction of microglial *APOE* expression in regions without Aβ pathology in APOE3/E3 and APOE3/E4 carriers, with no significant variation in microglial *APOE* expression in relation to Aβ in APOE4/E4 hippocampus (Figure 8E).

**Figure 8.**
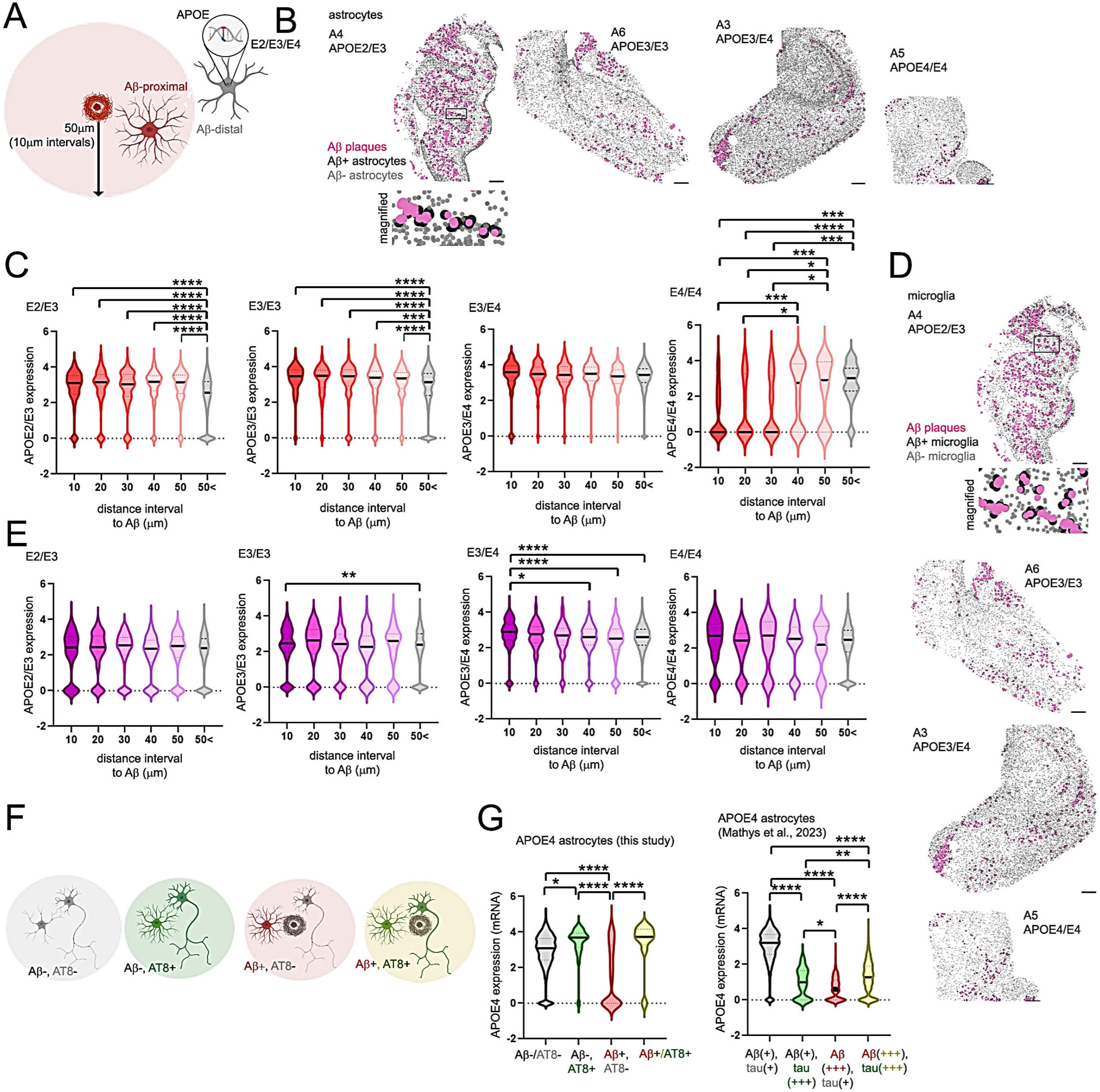
Astrocyte *APOE* expression in response to spatial proximity to pathology is defined by APOE genotype. (A) *APOE* expression in proximity (within 50 μm), or distal to (outside 50 μm) of Aβ plaques is quantified in AD hippocampus of varying APOE genotypes. (B) Aβ plaques (pink), Aβ+ (black) and Aβ-astrocytes (gray) in representative APOE2/E3 (A4), E3/E3 (A6), E3/E4 (A3) and E4/E4 (A5) hippocampus, scaler bar=1mm. (C) *APOE* mRNA expression in APOE2/E3, E3/E3, E3/E4 or E4/E4 astrocytes in 10 μm distance intervals to Aβ as indicated. (D) Image of Aβ plaques (pink), Aβ+ (black) and Aβ-microglia (gray) in representative APOE2/E3 (A4), E3/E3 (A6), E3/E4 (A3) and E4/E4 (A5) hippocampus, scaler bar=1mm. (E) *APOE* mRNA expression in APOE2/E3, E3/E3, E3/E4 or E4/E4 microglia in 10 μm distance intervals to Aβ as indicated. (F, G) Astrocyte *APOE* expression in APOE4/E4 samples is spatially defined by proximity to both Aβ and AT8 ptau pathology. (F) Schematic: *APOE* expression is quantified in astrocytes in Aβ-/AT8-, Aβ-/AT8+, Aβ+/AT8-or Aβ+/AT8+ regions in APOE4/E4 hippocampus (AD5). (G) *APOE* expression quantified in APOE4/E4 astrocytes in proximity to Aβ or AT8+ neurons as described in (F) from MERFISH (left) or snRNA-seq analysis (Mathys et al., 2023; right graph). In the snRNA-seq dataset, intermediate (+) or high (+++) pathology for Aβ and tau pathology are indicated. Statistical significance was determined in (C), (E) and (G) by one-way ANOVA with Tukey’s multiple comparison (*p<0.05, **p<0.01, ***p<0.001, ****p<0.0001).

We also determined whether proximity to AT8+ neurons can potentially influence astrocyte or microglia *APOE* expression; we found that astrocytes of all APOE genotypes featured slightly enhanced *APOE* expression in proximity to AT8+ neurons (Figure S18B). We also observed slight reduction in microglial *APOE* expression in AT8+ proximal APOE3/E3 and APOE3/E4 carriers (Figure S18C). Given that proximity to AT8+ pathology can potentially elevate *APOE* expression in astrocytes, we next determined whether Aβ and AT8 tau pathology in combination can affect astrocyte *APOE* expression in APOE4/E4 hippocampus (Figure 8F). Interestingly, APOE expression appeared to be specifically suppressed in astrocytes proximal to Aβ but distal to AT8+ neurons (Aβ+/AT8-); *APOE* expression in Aβ+/AT8-astrocytes significantly decreased compared to Aβ+/AT8+ astrocytes in APOE4/E4 hippocampus (Figure 8G). In APOE4/E4 microglia, we did not observe any significant alterations in APOE expression in proximity to Aβ or AT8+ neurons alone, or in combination (Aβ+/AT8+ microglia) (Figure S18D).

Given that these results implicate spatial sensitivity to Aβ and AT8+ pathology in APOE4/E4 astrocytes, we next determined whether spatial *APOE* expression trends could also be observed in snRNA-seq datasets in AD brain (Mathys, Peng et al. 2023). To this end, 8 AD APOE4/E4 individuals were categorized according to the presence of moderate (Cerad 2, Aβ+) to high (Cerad 3, Aβ+++) Aβ, as well as moderate (Braak III/IV, tau+) to high (Braak V/VI, tau+++) tau pathology in human prefrontal cortex (Mathys, Peng et al. 2023). Similar to our MERFISH results, *APOE* expression was attenuated in Aβ+ astrocytes from APOE4/E4 individuals with moderate (tau+) tau pathology (Figure 8G); however, APOE4/E4 individuals with moderate Aβ pathology and high AT8+ pathology (Aβ+/tau+++) showed significantly reduced *APOE* expression compared to astrocytes from Aβ+/tau+ individuals (Figure 8G). Interestingly, APOE4/E4 carriers clinically characterized with high Aβ and tau pathology (Aβ+++/tau+++) featured elevated astrocyte *APOE* expression compared to Aβ+++/tau+ or Aβ+/tau+++ astrocytes (Figure 8G), further suggesting that APOE4/E4 astrocytes may exhibit sensitivity to both Aβ and tau pathology.

## Discussion

Here we define cell-type-specific transcriptional profiles in spatial proximity and distal to Aβ and AT8 ptau pathology in human control and AD hippocampus (neuro-degeneration.org/admerfish). Transcriptome-based clustering identified potential microglia clusters (cluster 4) that were enriched in proximity to Aβ plaques, as well as expression-based cluster profiles in neurons and glia that were enriched or depleted in AT8+ neurons, or in glia in proximity to AT8+ neurons. We also observed a general enrichment of AT8+ neurons in proximity to Aβ plaques, which suggests that cumulative Aβ and tau pathology may be spatially related in the hippocampus. As tau and Aβ originate and spread from differing brain regions during AD onset (Braak and Braak 1995, Thal, Rub et al. 2002), tau and Aβ may initially develop independently (Mesulam 1999, Small and Duff 2008). According to the “amyloid hypothesis”, it is believed that Aβ is upstream of tau aggregation and dispersion which is consequential to AD pathology and onset (Hardy and Higgins 1992, Selkoe and Hardy 2016). So far, how Aβ influences neuronal tau pathology remains largely unknown. However, Aβ has been shown to induce and/or promote tau pathology in various mouse models. For example, Aβ has been shown to enhance neuritic dystrophy and tau phosphorylation in cultured neurons (Jin, Shepardson et al. 2011), and injection of Aβ in transgenic P301L tau mice can promote the formation of tau aggregates (Gotz, Chen et al. 2001). Conversely, injection of human tau seeds can also enhance neuritic tau pathology in APP knock-in (APP-KI) (He, Guo et al. 2018) and 5xFAD mouse brain (Vergara, Houben et al. 2019). Moreover, transgenic mice expressing both APP and human tau transgenes also show enhanced neurofibrillary tau tangle pathology (Lewis, Dickson et al. 2001, Heraud, Goufak et al. 2014). Despite potential implications for Aβ in promoting tau pathology in these experimental models in mouse brain, spatial relationships between Aβ plaques and tau pathology in human AD is less clear. For example, positive correlations between Aβ and AT8 tau pathology has been previously described using biochemical methods in control and AD frontal cortex, yet no direct correlation between extracellular Aβ plaques and AT8 is observed through immunohistochemical quantification (Malik, Miah et al. 2024). Interestingly, accumulation of intracellular neuronal Aβ can correlate with AT8 ptau in control frontal cortex (Malik, Miah et al. 2024), suggesting that Aβ accumulation early during onset can coincide with AT8 ptau levels. With the convergence of hippocampal Aβ plaque pathology (Thal phase 2 and onwards) (Thal, Rub et al. 2002) and tau during limbic stages of tau pathology (Braak stage III/IV) (Braak and Braak 1991), Aβ and tau may feature pathological interactions that retain spatial proximity at later stages of onset. Indeed, pS396 and AT8 ptau pathology was seen to coincide with the presence of Aβ plaques in AD hippocampus (Furcila, DeFelipe et al. 2018), further implicating spatial intersection of Aβ/tau pathology during AD onset.

Our results here define rs56867267 as a potential cis-eQTL within the MS4A4A locus, where the G allele is potentially associated with elevated *MS4A4A* expression in microglia distal to Aβ plaques. MS4A comprises a family of 18 tetraspanin-like four-pass transmembrane genes, 16 of which comprise a MS4A gene cluster on human Chromosome 11 (Eon Kuek, Leffler et al. 2016, Silva-Gomes, Mapelli et al. 2022). Polymorphisms within the MS4A locus have been implicated in various degenerative disorders, including AD (Ma, Yu et al. 2015), gastric cancer (Sun, Zhang et al. 2018), Progressive Supranuclear Palsy (Sanchez-Ruiz de Gordoa, Erro et al. 2020), and arthritis (Seyhan, Gregory et al. 2020). MS4A4A is highly expressed in microglia, and serves as a potential trafficking component in immune cell types (Cruse, Beaven et al. 2015). Specific polymorphisms at the MS4A4A locus have been specifically linked to AD onset (Harold, Abraham et al. 2009, Hollingworth, Harold et al. 2011, Naj, Jun et al. 2011, Allen, Zou et al. 2012, Ma, Jun et al. 2019). The MS4A4A-associated SNP rs1582763 was previously shown to modulate *MS4A4A* expression, and upregulation of *MS4A4A* increased soluble TREM2 (sTREM2) levels while *MS4A4A* downregulation decreased sTREM2 levels in human macrophages (Deming, Filipello et al. 2019). *Ms4a4a* deletion in 5xFAD mice reduced Aβ levels and featured enhanced expression of proinflammatory markers such as CD86 and Cxcr4, suggesting that MS4A4A may suppress some pro-inflammatory phenotypes in mice (Danhash, Verbeck et al. 2025). In human AD however, variants such as rs1582763 that enhance MS4A4A expression are thought to be protective, potentially promoting interferon response phenotypes while variants such as rs6591561 is linked to reduced *MS4A4A* expression and is associated with increased AD risk, and enhances microglia chemokine pathways (You, Brase et al. 2023). Our results indicate that some MS4A4A eQTL variants, namely rs56867267 may affect *MS4A4A* expression in microglia distal to Aβ plaques. Given that this SNP and its association with *MS4A4A* have not been previously reported, it will be of interest in future genetic studies to determine whether rs56867267 G/G, G/A, or A/A genotypes are associated with AD risk.

Our results also implicate ACE rs4575595-G/G as a cis-eQTL variant that potentially enhances *ACE* expression in excitatory neurons distal to Aβ plaques as well as AT8+ neurons. The ACE gene encoding angiotensin I converting enzyme (ACE1), is required for converting renin-cleaved angI to angII which then mediates vasoconstriction or vasodilation through AT1R/AT2R receptors (Kehoe 2018). In addition to regulating blood pressure through the renin-angiotensin system (RAS), ACE1 has also been shown to degrade Aβ (Hu, Igarashi et al. 2001, Hemming and Selkoe 2005, Oba, Igarashi et al. 2005). However, whether ACE1-mediated Aβ degradation can reduce Aβ plaque deposition *in vivo* remains unclear, as use of ACE1 inhibitors (Eckman, Adams et al. 2006, Hemming, Selkoe et al. 2007) or ACE deletion (Eckman, Adams et al. 2006) failed to modulate Aβ deposition. ACE inhibition enhanced Aβ deposition in Tg2576 mouse brain (Zou, Yamaguchi et al. 2007), suggesting that ACE1 can mediate plaque clearance in some instances. Interestingly, an ACE1 R1279Q (rs4980) variant was identified as a potential enhancer of neurodegenerative features in 5xFAD brain which may be more prominent in female animals (Cuddy, Prokopenko et al. 2020). While effects of modulating of ACE1 expression remain somewhat unclear in AD brain, growing evidence implicates coding variants such as R1279Q in AD pathogenesis.

As the most potent sporadic genetic risk factor in AD, carriers of a single APOE4 allele feature increased risk of AD onset by ∼3 to 4 fold, while two copies of APOE4 can increase AD risk up to ∼15-fold (Corder, Saunders et al. 1993, Saunders, Strittmatter et al. 1993, Roses 1996). APOE4 has been shown to enhance AD-associated Aβ plaque deposition and cerebral amyloid angiopathy (CAA) compared to APOE2 and APOE3 carriers (Serrano-Pozo, Qian et al. 2015). APOE4 carriers are also clinically associated with elevated tau and phosphorylated tau levels during prodromal and mild cognitive impairment (MCI) stages of AD onset (Liu, Tan et al. 2016, Mandecka, Budziszewska et al. 2016). Female APOE4 carriers are also associated with higher AD risk (Farrer, Cupples et al. 1997), as well as enhanced AD pathology (Altmann, Tian et al. 2014) and accelerated cognitive decline (Lin, Choudhury et al. 2015) compared to male carriers. In addition to clinical implications for APOE4 in AD, APOE4 can aggravate neurodegenerative phenotypes in mouse models. For example, APOE4 expression in humanized knock-in PS19 tauopathy mouse models can aggravate tau pathology, atrophy and neuroinflammatory effects compared to APOE3 or APOE2 animals (Shi, Yamada et al. 2017). Specific deletion of APOE4 in astrocytes has also been shown to reduce tau pathology and pathological expression signatures in various cell types in murine PS19/APOE4 knock-in (KI) models (Wang, Xiong et al. 2021), although neuronal APOE4 deletion in similar PS19/APOE KI models can also attenuate tau pathology and gliosis (Koutsodendris, Blumenfeld et al. 2023). APOE4 expression in APP/PS1 mice is also seen to enhance amyloid seeding (Liu, Zhao et al. 2017), and use of genetic exon-swapping strategies has also shown that switching expression of APOE4 to APOE2 isotypes in 5xFAD mouse models can reduce amyloid pathology and gliosis (Golden, Siano et al. 2025). Given that murine microglial APOE expression is enhanced in 5xFAD (Keren-Shaul, Spinrad et al. 2017) as well as EAE, SOD1 and APP/PS1 mouse models of neurodegeneration (Krasemann, Madore et al. 2017), APOE expression is likely regulated and potentially sensitive to proteotoxicity, pathological features and genotype in human brain. Our results indicate that although microglia *APOE* is upregulated in proximity to Aβ plaques in AD hippocampus, this general upregulation is unaffected by APOE genotype. In contrast, we find that APOE4/E4 astrocytes feature reduced *APOE* expression in proximity to Aβ plaques, suggesting that APOE4/E4 astrocytes may comprise mechanisms to downregulate *APOE* expression in the presence of Aβ. In agreement with this notion, APOE expression is specifically reduced in APOE4/E4 in comparison with APOE3/E3 iPSC-derived astrocytes with Aβ exposure (Lin, Seo et al. 2018).

Together, these results establish spatially-defined cell expression atlas, integrating Aβ/AT8 ptau pathology together with genotype-specific expression profiles in 120 AD risk loci. Given the limited size of the MERFISH cohort described in this study, we have integrated our analyses with larger snRNA-seq cohorts; with the rapidly growing number of AD-related datasets available in the field, further comparison and validation with MERFISH analysis described here will be necessary. Moreover, as more loci related to AD risk are being identified, it may also be necessary to include additional risk loci with larger gene panels. With this current study as an example, we anticipate development of future studies that integrate datasets across multiple platforms to enrich spatial and genetic variant-based expression profiles.

## Methods

### Whole genome sequencing data collection

Genomic DNA was extracted using the PureLink Genomic DNA Mini Kit (Thermo Fisher Scientific, Waltham, MA, USA) according to the manufacturer’s instructions. DNA concentration was determined using a Qubit 2.0 Fluorometer (Thermo Fisher Scientific, Waltham, MA, USA). DNA integrity was assessed by electrophoresis on a ∼1% agarose gel, with 50 to 100 ng of DNA loaded per lane. Library preparation was performed using the NEBNext® Ultra™ DNA Library Prep Kit for Illumina® (New England Biolabs, Ipswich, MA, USA) according to the manufacturer’s protocol. Briefly, genomic DNA was fragmented by acoustic shearing using a Covaris S220 instrument (Covaris, Woburn, MA, USA). Fragmented DNA was subsequently purified and subjected to end repair. Following 3′ adenylation, adapters were ligated to the DNA fragments, and libraries were enriched by limited-cycle PCR amplification. Library quality and fragment size distribution were assessed using High Sensitivity D1000 ScreenTape on the Agilent TapeStation System (Agilent Technologies, Santa Clara, CA, USA). Libraries were quantified using a Qubit 2.0 Fluorometer and further quantified by quantitative real-time PCR (Applied Biosystems, Carlsbad, CA, USA) prior to clustering and sequencing on a HiSeq Illumina platform (2×150 bp pair-end reads with targeted sequencing depth at 30x).

### MERFISH data collection

MERFISH data was generated using the commercial MERSCOPE platform by Vizgen. Sample preparation and imaging was performed following the manufacturer’s instructions (MERSCOPE Fresh and Fixed Frozen Tissue Sample Preparation User Guide, Doc. number 91600002). Fresh frozen tissues from human AD and control hippocampus with an RNA integrity number (RIN) > 5 were sectioned into 10 μm-thick slices using a cryostat and mounted on MERSCOPE slides. Sections were then fixed with 4% PFA at 37°C for 30 minutes, followed by permeabilization with 70% ethanol overnight. On the following day, autofluorescence was quenched by exposing sections to UV light for 5 hours using the MERSCOPE Photobleacher instrument. Sections were subsequently stained with antibodies to detect Amyloid-beta (1:500, ab205340, Abcam) and Phospho-Tau (1:100, MN1020, Thermo Fisher Scientific) at room temperature (RT) for 1 hour, followed by incubation with Anti-Mouse Aux 4 (PN 20300101), Anti-Rabbit Aux 5 (PN 20300102), and Anti-Goat Aux 6 (PN 20300103) at RT for an additional hour. A pre-designed 300-gene panel was used for probe hybridization, which was carried out at 37°C for 36 hours in an ACD oven. After hybridization, the sections were washed twice with a formamide buffer at 47°C for 30 minutes each and then embedded in gel. Following gel embedding, the tissue samples were cleared using a clearing solution supplemented with proteinase K for 24-48 hours at 37°C, until no visible tissue remained in the gel. After clearing, the sections were stained with DAPI and PolyT, then fixed with formamide prior to imaging. MERSCOPE images were acquired according to the MERSCOPE Instrument Preparation Guide (Doc. number 91500001). Briefly, the imaging kit was thawed at 37°C for 60 minutes, activated, and loaded into the instrument. The MERSCOPE flow chamber was assembled with the stained tissue sections, fluidics were primed, and the flow chamber was filled with liquid. A 10X low-resolution image was first taken, and based on DAPI staining, a region of interest (ROI) was selected for 60X high-resolution MERSCOPE imaging.

### MERFISH data processing

The MERSCOPE instrument employed built-in processing modules to decode RNA transcript locations and perform preliminary cell segmentation. All postprocessing and segmentation procedures were performed using the Vizgen-postprocessing tool (https://github.com/Vizgen/vizgen-postprocessing, v1.0.2). Final cell segmentation was performed using the CellPose method integrated in the Vizgen-postprocessing tool, using DAPI and PolyT staining images from three z-layers taken at 3 μm, 6 μm and 9 μm. A cell × gene expression matrix was generated based on segmented cell boundaries.

Quality control was performed individually for each sample, and cells with less than 5 genes or less than 20 molecules were filtered out. We further removed cells with volumes less than 100 μm^3 or larger than three times the median volume of all cells. We then applied Scrublet (v0.2.3) (Wolock, Lopez et al. 2019) to identify potential doublets and removed cells with double scores equal to or larger than 0.2. Total counts per cell were normalized over the cell volume (i.e. cell counts/cell volume) for the remaining cells. Cells with normalized total counts within the top and bottom 2% quantile were removed. Finally, expression of each gene was normalized per cell where the sum of all expression values per cell equaled 250. The normalized gene expression values were further log-transformed and then z-scaled with the maximum value being 10. Data normalization and scaling were performed using Scanpy (v1.10.3) (Wolf, Angerer et al. 2018). Log-transformed gene expression values before z-scaling were used for differential expression analysis.

### MERFISH data analysis

QC-passed cells from all samples were integrated together for joint analysis using Scanpy. Raw count matrices of all MERFISH samples were concatenated, followed by normalizing total expression values per cell to 250, further applying log-scaling with a pseudocount of 1 and subsequent z-scaling with a maximum value of 10. Principal component analysis (PCA) was performed on z-scaled values. Harmony (v2.0.0) (Korsunsky, Millard et al. 2019) was used to correct for batch effects and integration of different samples by PCA embedding adjustment. Then the neighborhood graph was computed by Harmony embedding with “n_neighbors” being 10, and UMAP embedding was generated based on the neighborhood graph. Leiden clustering was used to cluster cells with “resolution=2”, and cell clusters were annotated into cell types based on marker gene expression.

To prevent potential false positive signals from biasing downstream analysis, we performed gene filtering in the MERFISH dataset. Infrequently-expressed genes and genes expressed at low levels were removed based on expression profiles from an external snRNA-seq dataset (SEA-AD) (Gabitto, Travaglini et al. 2024). Cells were grouped into major brain cell types according to the original cell type annotation. We built a filter based on gene expression frequency and average read counts for each major cell type. We calculated average read counts among cells with non-zero expression for each gene (as opposed to overall average expression) to avoid removal of genes expressed in a small cell population. Genes expressed in no more than 15% cells, with no more than 1.5 read counts on average among cells with non-zero expression were filtered out. The gene filtering procedure was only applied as a post-filtering step of the differential expression analysis results unless otherwise specified.

### Pathology segmentation and quantification

Aβ plaques were segmented from pathological staining images. For each tissue slide, the MERSCOPE platform generated pathological staining images at 7 z-stacks corresponding to different tissue depths. Images from different z-stacks were processed independently, and combined to ensure consistency across z-stacks and finally to segment Aβ plaques, as explained below. For each stained greyscale image, we applied a Gaussian filter with “sigma=2” using the filters.gaussian function from skimage (van der Walt, Schonberger et al. 2014) to remove high-frequency image noises. This Gaussian filter outputs smoothed per-pixel intensity values in the range from 0 to 1. Our assumption is that background signal intensities of non-tissue regions and tissue regions without Aβ plaques are different, and both are lower than Aβ-stained signals. Based on this assumption and the actual data properties, we proposed a custom thresholding method to separate background and Aβ signals based on pixel intensity distribution, as detailed below. Consistent with our assumption, we observed two sharp peaks with intensities lower than 0.2 from background signal distributions corresponding to the non-tissue region and tissue regions without Aβ plaques, and the distribution of Aβ staining signals featured large variance across samples with Aβ pathology. To differentiate between background signals and Aβ signals based on observation that there are many local small peaks that appear between background and Aβ-positive signals, our thresholding method identifies the first peak with pixel intensities >0.2, together with increased pixel numbers/frequency compared to the last peak as the threshold. The image is binarized with pixel intensities greater than the threshold, thereby representing genuine Aβ signals. Sums from binarized images across different z-stacks were generated, and the aggregated image is further binarized so that only pixels being identified as Aβ signals by at least 3 z-stacks were considered as the final Aβ signals. Aβ plaques were then segmented from aggregated-binarized images using the “label” function from skimage, and Aβ plaques with sizes less than 40 μm^2^ were filtered out.

AT8 ptau stained images were binarized using custom pixel density thresholds to identify AT8 ptau pixels. AT8 ptau pixels are used to quantify tau pathology level in individual cells. For each cell, we approximated the smallest circular area extending from the cell center to cover the cell body using the cell bounding box output by CellPose to estimate the cell radius. The proportion of AT8 ptau pixels were calculated within the circular area covering the cell body, multiplied by 1,000,000 and transformed by log10 with a pseudo_count of 1 to compute the ptau score for each cell.

### Pathology characterization of cells

Cells were annotated based on their proximity to Aβ plaques. For each cell, we calculated the distance from the cell center to the boundary of the nearest Aβ plaque. Cells were further stratified based on their distance to the nearest Aβ plaque within different distance intervals: <10 (including when overlapping with the Aβ plaque), [10, 20), [20, 30), [30, 40), [40, 50], and >50 μm. To simplify the annotation categories, cells were also grouped into two categories including <=50 (Aβ+) and >50 μm (Aβ-).

Tau categorization is based on AT8 ptau scores in neurons. Given that tau pathology is an intracellular pathology enriched in neurons, we classified excitatory and inhibitory neurons into low (ptau score <=1), moderate (ptau score >1 and <5) and high (ptau score >=5) categories. For non-neuron cells, AT8 + (AT8 ptau+) cells were defined by a “tau high” neuron within 30 microns from its cell center, or AT8-if neither “tau high” nor “tau moderate” neurons reside within 30 microns from a non-neuronal cell.

### Mapping spatially-defined MERFISH profiles to snRNA-seq datasets

We trained machine learning models using spatially-defined MERFISH data to identify potential pathology-proximal and distal cells in snRNA-seq datasets (Mathys, Boix et al. 2024). MERFISH cells used for model training were subsampled within each sample to balance the numbers of cells proximal and distal to pathology while maintaining similar distributions of detected gene and transcript counts between the two groups. We tested Extreme Gradient Boosting (XGBoost) and Linear Regression (LR) methods. For XGBoost, ranked and quantile-normalized gene expression values were used as input. We used a grid-search method to optimize model hyperparameters, and used five-fold cross-validation to assess model robustness and reduce the risk of overfitting. The final model used for predicting snRNA-seq cells were trained using all selected MERFISH cells. We normalized snRNA-seq data so that every cell has a total count of 1,000,000, followed by taking logp1p. For the LR method, we used log-normalized gene expression values as input for both MERFISH and snRNA-seq data. The two methods showed high prediction consistencies in all our tasks. We ultimately used and applied XGBoost predictions due to its capability to model non-linear relationships across genes. Pathology-proximal and pathology-distal groups were determined by selecting the top and bottom 20% ranked by the prediction scores in descending order.

### Differential expression analysis

Differential expression (DE) analysis within the MERFISH dataset was performed using the “rank_genes_groups” function with the Wilcoxon rank-sum test from Scanpy. Log-normalized gene expression values were used for DE analysis, and log2 fold-change (L2FC) was calculated using gene expression values after normalizing total expression values to a constant per cell, followed by log2 transformation with pseudo_count of 0.01. Significantly differentially expressed genes comprised DEGs expressed in at least 10% cells in either groups, absolute L2FC values at least 0.2, and adjusted p value less than 0.05 by default if not otherwise specified.

### Variant calling from whole genome sequencing data

Paired-end reads from WGS were mapped to the reference human genome GRCh37 using BWA-MEM aligner (v0.7.17) (Li and Durbin 2009) to facilitate comparison with an external WGS dataset that had also been processed using GRCh37. Other variant calling procedures were performed using modules from GATK (v4.4.0.0) (McKenna, Hanna et al. 2010). Reads from PCR duplicates were marked using the “MarkDuplicates” module. The “BaseRecalibrator” module was used to estimate sequencing-error to further calibrate base qualities of the reads. We subsequently used the “HaplotypeCaller” module to call the variants and infer the haplotypes using local-assembly for each individual. Variants called from different individuals were combined using the “CombineGVCFs” module. Joint genotyping across individuals was then performed using the “GenotypeGVCFs” module. We used the “VariantRecalibrator” to build the statistical model to assess variant quality. Lastly, the “ApplyVQSR” module processed the model produced by “VariantRecalibrator” and marks/filters variants.

### Pathology-dependent eQTL analysis

Pathology-dependent eQTL analysis was performed for cells proximal or distal to pathology independently. For pathology-proximal or distal cells of each cell type from every individual, we calculated pseudo-bulk gene expression profiles by summing up read counts for each gene across cells. For each cell type proximal or distal to pathology, we calculated pseudo-bulk read count matrix of gene × individuals. Size factors of the pseudo-bulk read count matrix were calculated using the weighted trimmed mean of M-values (TMM) method using the “calcNormFactors” function in edgeR (v3.42.4) (Robinson, McCarthy et al. 2010). The count matrix was then normalized to log2 counts per million mapped reads (CPM) using the “voom” function from limma (v3.58.1) (Ritchie, Phipson et al. 2015). To further reduce sensitivity to extreme expression values, we ranked log2(CPM) values and performed quantile normalization across samples for each gene. eQTL analysis was performed using the MatrixEQTL method (v2.3) (Shabalin 2012) considering covariates including sex, age and post mortem interval (PMI). We identified cis-eQTLs within 10kb up/downstream of gene body regions. eQTLs were defined as one variant significantly associated with variations in gene expression of an associated gene (nominal p value < 0.05). eGenes were defined as genes comprising at least one eQTL.

For pathology-dependent eQTL analysis using an external cohort comprising snRNA-seq datasets in 424-individual, we classified pathological groups within the cohort based on neuritic plaque CERAD score into 149 Aβ-individuals (CERAD=0 or 1) and 275 Aβ+ individuals (CERAD = 2 or 3), as well as groupings based on Braak score into 62 low tau pathology individuals (Braak=0, 1 or 2) and 362 high tau pathology individuals (Braak=3, 4, 5 or 6). Generation of pseudo-bulk gene expression profiles and eQTL analysis employed the same pipeline used in processing the MERFISH dataset as described above.

The variants were annotated using the Ensembl Variant Effect Predictor (VEP release 160) (McLaren, Gil et al. 2016). VEP annotates the consequence of each variant on each transcript isoform. For predicted eQTLs between variants and genes, the consequence of each variant on a gene is summarized across transcript isoforms of this gene and represented by the most severe consequence.

## Data availability

MERFISH data generated in this study is hosted in a website for online visualization and exploration (neuro-degeneration.org/admerfish). We downloaded the following datasets from public AD studies. SEA-AD snRNA-seq data of human middle temporal gyrus (Hawrylycz, Kaplan et al. 2024) were downloaded from Synapse (https://adknowledgeportal.synapse.org/Explore/Studies/DetailsPage/StudyDetails?Study=syn26223298). snRNA-seq of human Prefrontal Cortex (PFC) (Mathys, Davila-Velderrain et al. 2019) were downloaded from Synapse (https://www.synapse.org/#!Synapse:syn18485175). snRNA-seq data from 6 human brain regions including hippocampus (Mathys, Boix et al. 2024) were downloaded from Synapse (https://www.synapse.org/Synapse:syn52293442). snRNA-seq data of the human prefrontal cortex (Mathys, Peng et al. 2023) were downloaded from Synapse (https://www.synapse.org/Synapse:syn52293417). snRNA-seq data of human Dorsolateral Prefrontal Cortex (DLFPC) samples from 424 individuals (Fujita, Gao et al. 2024) (https://www.synapse.org/Synapse:syn31512863) and associated variants called from WGS (De Jager, Ma et al. 2018) (https://www.synapse.org/Synapse:syn11724057) were downloaded from Synapse. The reference human genome GRCh37 was downloaded from http://ftp.1000genomes.ebi.ac.uk/vol1/ftp/technical/reference/human_g1k_v37.fasta.gz. The annotated SNP information in dbSNP was downloaded from http://ftp.1000genomes.ebi.ac.uk/vol1/ftp/technical/reference/phase2_mapping_resources/ALL.wgs.dbsnp.build135.snps.sites.vcf.gz.

## Code availability

Analysis codes will be available on Github and other platforms if required before official manuscript acceptance.

## Acknowledgments

We thank all prior (Ke Xu, Andre Batista, Wenqi Yang) and current members of the Huang laboratory, and prior member (Amanda Qiao) of Yip laboratory for project-related discussion and input. Specially thanks to G. Garcia (SBP Histopathology Core) for assistance with tissue processing and data acquisition.

## Author contributions

Project conceptualization and probe design (B.Z., J.P-C., T.Y.H., H.H., G.V.V.), scientific input and discussion (B.Z., J.P-C., T.Y.H., H.H., G.V.V., C.H.S., K.Y.Y.), sample preparation and data acquisition (B.Z., J.P-C., G.V.V.), data processing, analysis and interpretation (C.H.S., W.N.C., K.Y.Y., T.Y.H.), data portal building (C.H.S), manuscript preparation and editing (C.H.S., T.Y.H., K.Y.Y.), funding (T.Y.H.), project supervision (T.Y.H., K.Y.Y.). All authors have read and agreed to the submitted version of the manuscript.

## Competing interests

The authors declare that they have no competing interests.

## Funding

This work was supported by grants from the National Institutes of Health/National Institute on Aging (R01 AG070391, R01 AG056130 and R01 AG085498) to T.Y.H.

## Supplemental Figures

**Figure S1.**
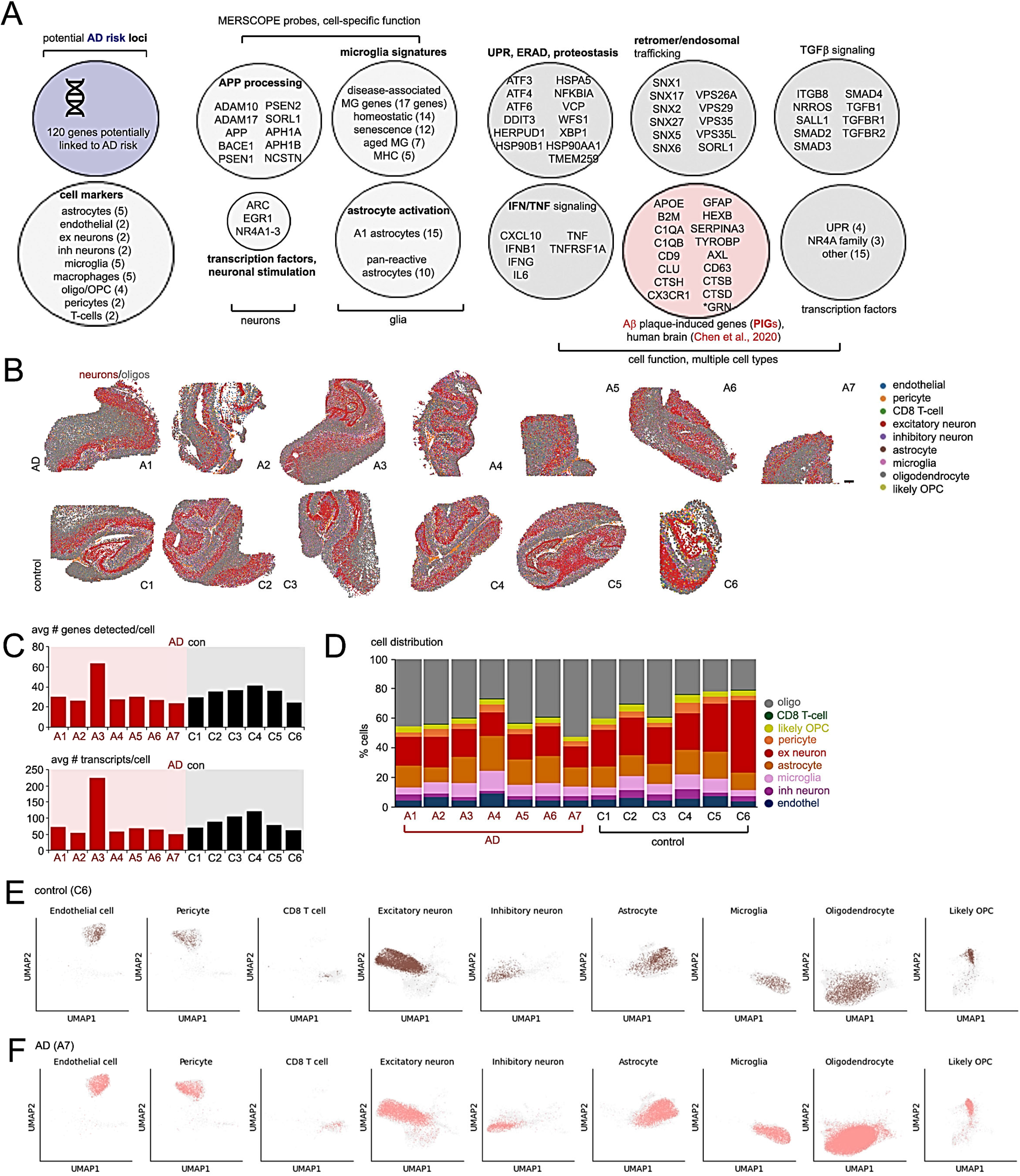
Spatial annotation of cell types and transcripts in control and AD hippocampus. (A) MERSCOPE probe design. A 300-probe panel comprising 120 genes potentially linked to AD risk, marker genes to annotate various cell types, and probes to define various aspects of cell function. A complete annotated list of probes used in the study are listed in Table S3. (B) Annotated cell types in AD (A1-A7) and control (C1-C6) hippocampus. Excitatory neurons are depicted in red, and oligodendrocytes are indicated in gray; color-coded annotations of other cell types are indicated in the adjacent legend. (C) Average number of genes (upper graph) and transcripts (lower graph) detected per cell in control and AD hippocampus. (D) Percentage distribution of annotated cell types in individual control and AD hippocampal sections. Color-coded indicators for annotated cell types are shown in the adjacent legend. (E, F) Spatial distribution of different cell types from the youngest (E) and oldest (F) donors in our cohort. Cells from the corresponding young/old samples were highlighted while cells from other samples are in grey.

**Figure S2.**
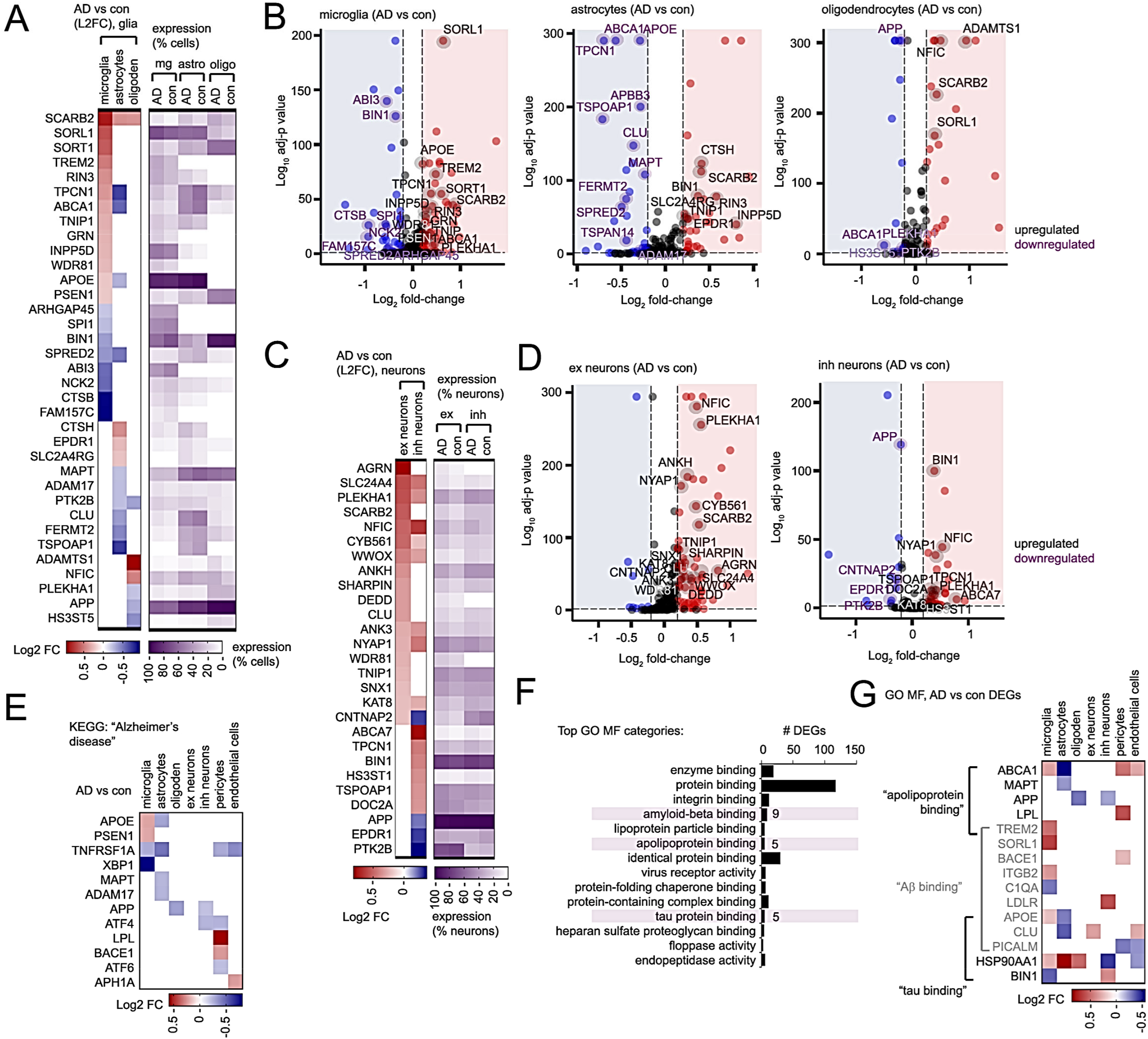
Cell-specific expression in AD and control hippocampus. (A) Red/blue-scaled heatmaps depicting Log2 fold-change in AD vs control DEGs related to AD-risk in glia (left heatmaps). Adjacent purple-scaled heatmaps indicate relative percent expression of genes shown in AD and control microglia (mg), astrocytes (astro), oligodendrocytes (oligo). (B) Volcano plots depicting up and downregulated AD vs control DEGs in microglia (left), astrocytes (middle) and oligodendrocytes (right). Upregulated and downregulated genes related to AD risk are highlighted in black and purple. (C) Red/blue-scaled heatmaps depicting Log2 fold-change in AD vs control DEGs related to AD-risk in neurons (left heatmaps). Adjacent purple-scaled heatmaps indicate relative percent expression of genes shown in AD and control excitatory (“ex”) and inhibitory (“inh”) neurons. (D) Volcano plots depicting up and downregulated AD vs control DEGs in excitatory (ex) neurons (left) and inhibitory (inh) neurons (right). Upregulated and downregulated genes related to AD risk are highlighted in black and purple. (E) Heatmap depicting AD vs control Log2 fold-change of DEGs involved in GO KEGG “Alzheimer’s disease” in the cell types indicated. (F, G) (F) Top GO Molecular Function (MF) categories enriched in AD vs control DEGs in all cell types. (G) Heatmaps depict Log2 fold-change in AD vs control DEGs related to GO MF “apolipoprotein binding”, “Aβ binding” and “tau binding” categories in various hippocampal cell types.

**Figure S3.**
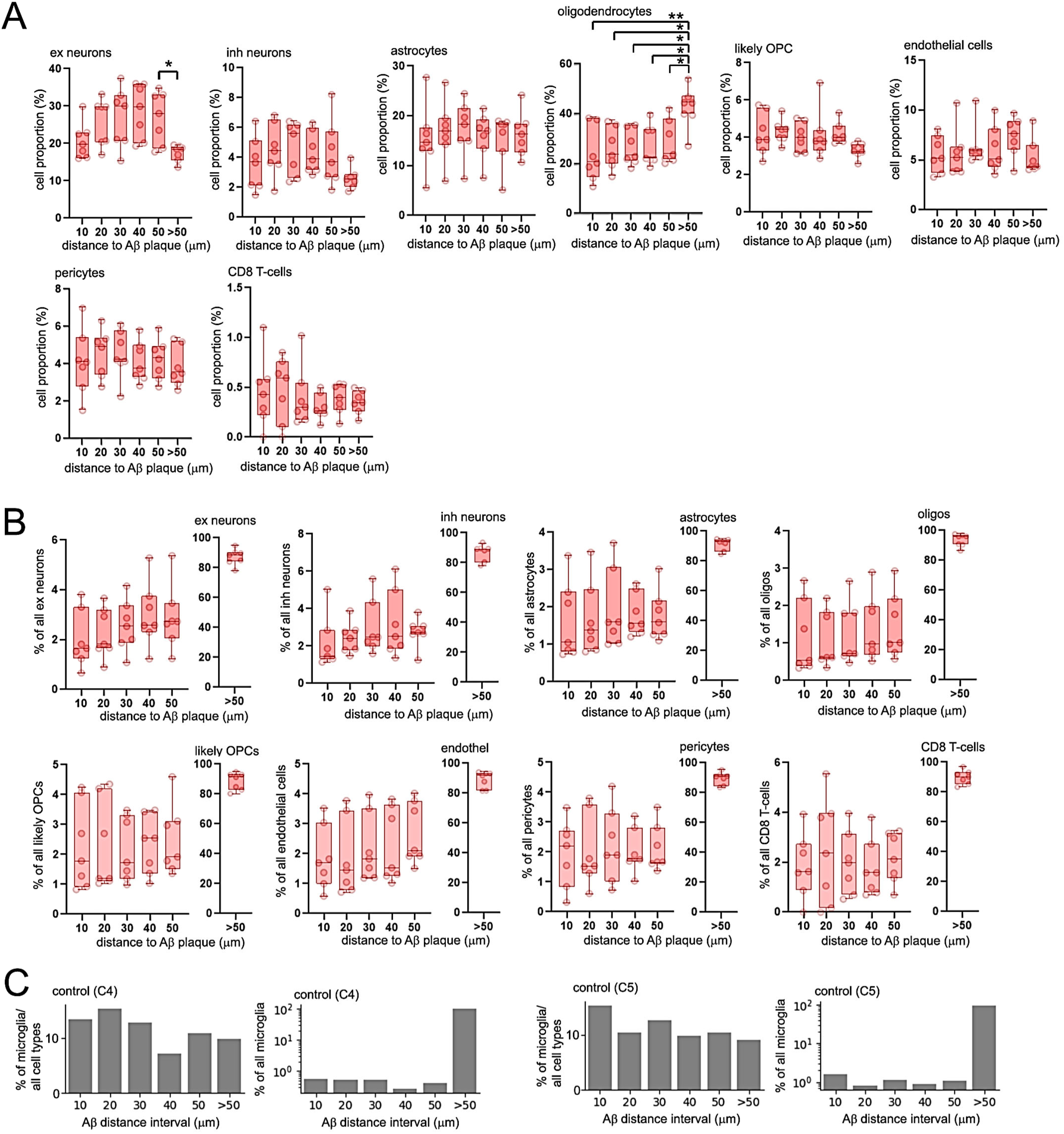
Characterizing cell-specific distribution of cells proximal or distal to Aβ plaques. (A) Box and whisker plots depicting percentage proportion of cells within the distance interval to Aβ plaques in AD hippocampus. Percentages were calculated by normalizing to all cell types within a distance interval. (B) Box and whisker plots depicting percentage proportion of cells within the distance interval to Aβ plaques in AD hippocampus; percentages were calculated by normalizing within all distance intervals for a specific cell type. (C) Percentages of cells within difference distance intervals to Aβ plaques for two control samples (C4 and C5) with Aβ plaques. Percentages were calculated by normalizing within each distance interval across all cell types, and by normalizing within microglia across all distance intervals. Statistical significance was determined by one-way ANOVA (*p<0.05, **p<0.01).

**Figure S4.**
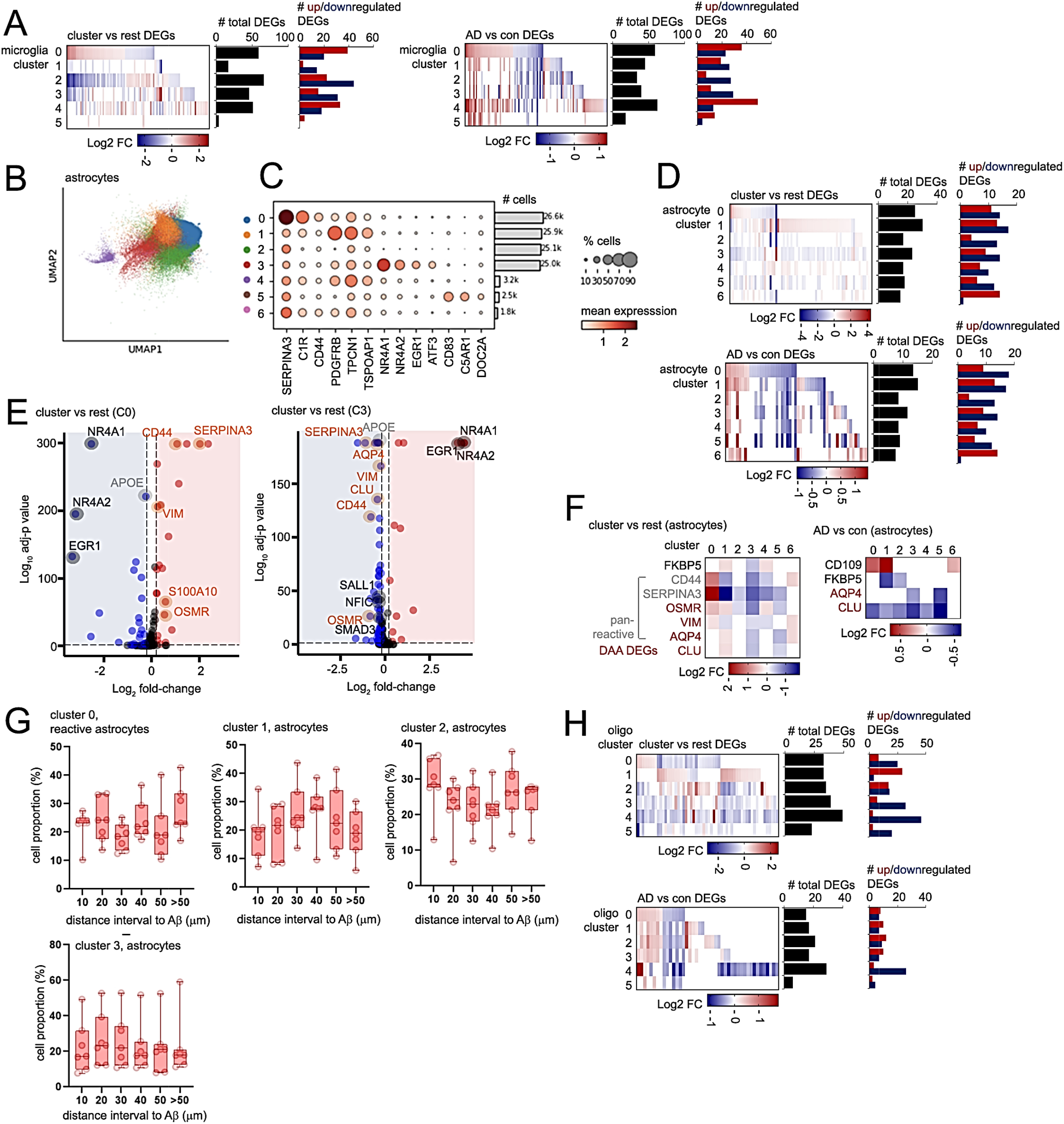
Cluster specific DEG profiles in glia. (A) Heatmap depicting Log2 fold-change cluster specific DEGs (“cluster vs rest”, left heatmap) or AD vs con profiles (right heatmap) in microglia. Adjacent graphs depict number of total (black bars) and up (red) or down-regulated (blue) microglia DEGs. (B) UMAP subcluster representation of 110,137 astrocytes in control and AD hippocampus. A total of 7 astrocyte subclusters (0 to 6) were identified as represented by colors indicated in the legend. (C) Dot plot indicating fraction of cells (%) and level of expression of top markers of clusters. Percentage of cells expressing genes (scaled dot size) or mean expression (red heat scale) are indicated. Bar graph indicates number of cells identified in each cluster. (D) Heatmap depicting Log2 fold-change cluster specific DEGs (“cluster vs rest”, top heatmap) or AD vs con profiles (bottom heatmap) in astrocytes. Adjacent graphs depict number of total (black bars) and up (red) or down-regulated (blue) astrocyte DEGs. (E) Volcano plots for astrocyte clusters 0 (left) and cluster 3 (right) enriched for DAA/transcription factor DEGs in “cluster vs rest” comparisons. Upregulated Disease Associated Astrocyte (DAA) DEGs (orange) or transcription factors (black) are indicated (adj-p<0.05, absolute L2FC ≥ 0.2). (F) Heatmaps depicting Log2 fold-change in astrocyte DAA (red), or pan-reactive (gray) DEGs in astrocyte clusters 0-6 unique to each cluster (“cluster vs rest”, left heatmap) or AD vs control (right heatmap) comparisons. (G) Box and whisker plots indicating percentage of astrocytes identified in different distance intervals surrounding Aβ plaques in clusters 0 to 2 (top), and cluster 3 (bottom) astrocytes. Percentages were calculated as a fraction of all astrocytes within a distance category from all clusters. (H) Heatmap depicting Log2 fold-change cluster specific DEGs (“cluster vs rest”, top heatmap) or AD vs con profiles (bottom heatmap) in oligodendrocytes. Adjacent graphs depict number of total (black bars) and up (red) or down-regulated (blue) oligodendrocyte DEGs. Statistical significance in (G) was determined by one-way ANOVA.

**Figure S5.**
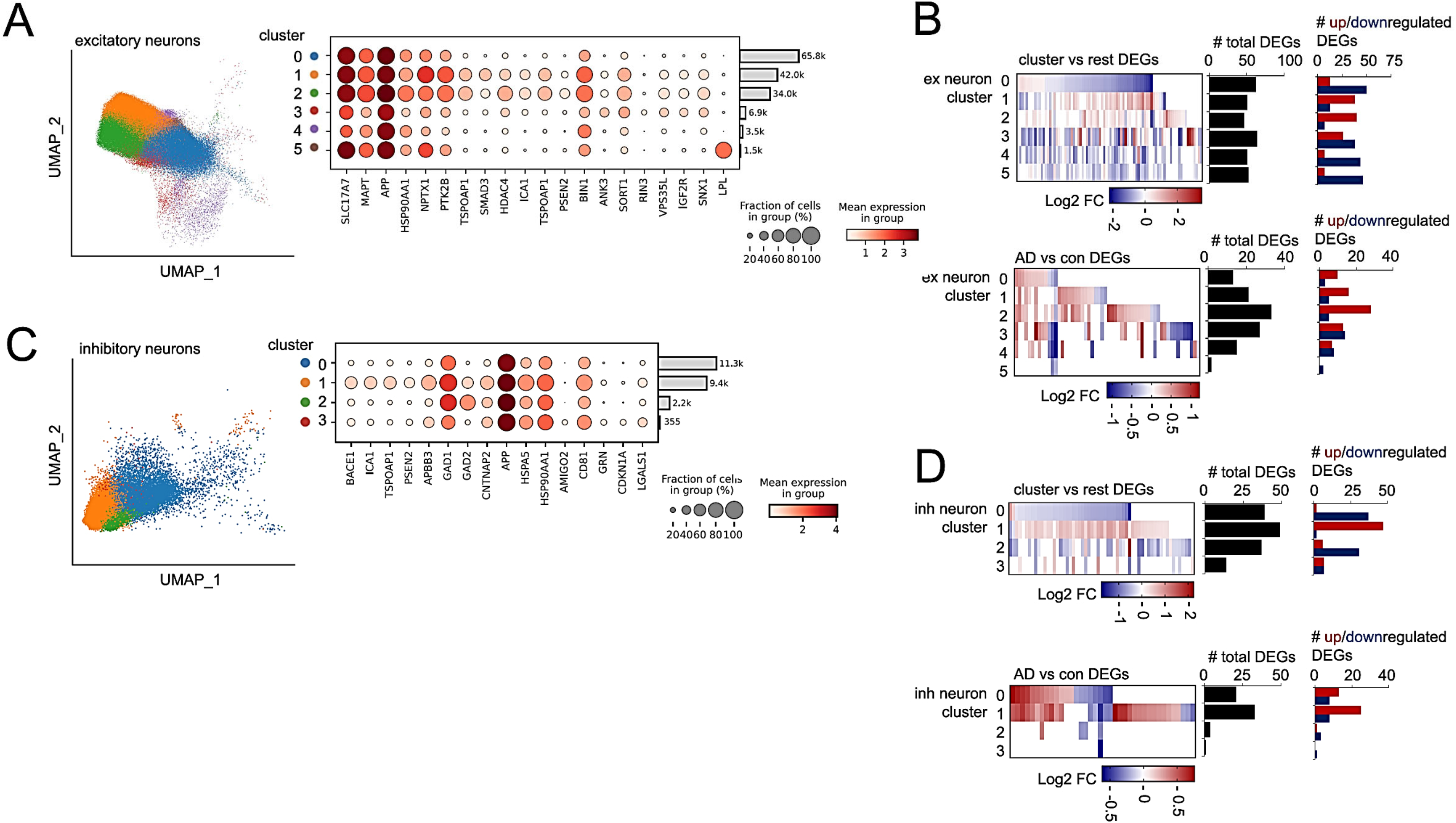
Cluster specific DEG profiles in neurons. (A) UMAP subcluster representation of 153,730 excitatory (ex) neurons in control and AD hippocampus. A total of 6 excitatory neuron subclusters (0 to 5) were identified as represented by colors indicated in the legend. Adjacent dot plot indicating fraction of cells (%) and level of expression of various ex neuron transcripts in clusters indicated. Percentage of cells expressing ex neuron transcripts (scaled dot size) or mean expression (red heat scale) are indicated. Bar graph indicates number of cells identified in each cluster. (B) Heatmap depicting Log2 fold-change of cluster specific DEGs (top) or AD vs con DEGs (bottom heatmap) in ex neurons. Adjacent graphs depict number of total (black bars) and up (red) or down-regulated (blue) oligodendrocyte DEGs. (C) UMAP subcluster representation of 23,324 inhibitory neurons in control and AD hippocampus. A total of 4 inhibitory neuron subclusters (0 to 3) were identified as represented by colors indicated in the legend. Adjacent dot plot indicating fraction of cells (%) and level of expression of various inhibitory neuron transcripts in clusters indicated. Percentage of cells expressing inhibitory neuron transcripts (scaled dot size) or mean expression (red heat scale) are indicated. Bar graph indicates number of cells identified in each cluster. (D) Heatmap depicting Log2 fold-change of cluster specific DEGs (top) or AD vs con DEGs (bottom heatmap) in inhibitory (inh) neurons. Adjacent graphs depict number of total (black bars) and up (red) or down-regulated (blue) inh neuron DEGs.

**Figure S6.**
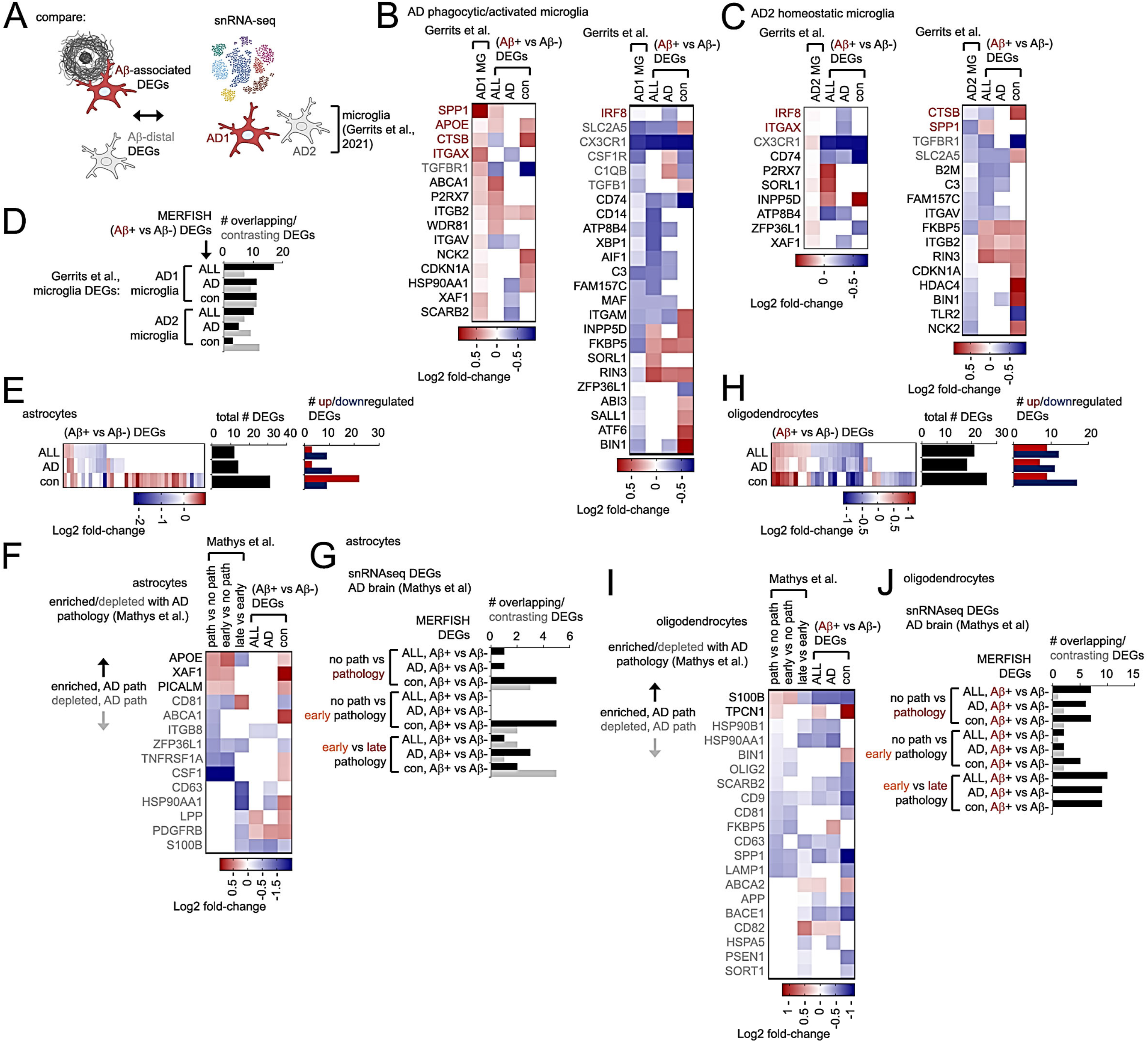
Characterizing Aβ-associated expression profiles in hippocampal glia. (A) Schematic depicting comparison of Aβ-associated microglia expression profiles identified through MERFISH analysis (this study) with snRNA-seq datasets (Gerrits et al.) from human AD brain. (B, C) Comparison of Aβ-proximal expression profiles with activated/homeostatic snRNA-seq microglia profiles in human AD brain (Gerrits et al.). Aβ-associated microglia (Aβ+ vs Aβ-) DEGs identified through MERSCOPE analysis are aligned to phagocytic/activated “AD1” (B) or homeostatic “AD2” microglia (C) as previously described in Gerrits et al. (E-G) Characterizing Aβ-proximal profiles in astrocytes. (E) Heatmap depicting Log2 fold-change of DEGs from Aβ+ vs Aβ-astrocytes in all, AD or control hippocampus. Adjacent graphs depict number of total (black bars) and up (red) or down-regulated (blue) astrocyte DEGs. (F) Heatmap of aligned Log2 fold-change profiles in astrocytes from prefrontal cortex (Mathys et al., 2019) and Aβ+ vs Aβ-DEGs (MERFISH, this study). (G) Graph summarizing overlap between snRNA-seq astrocyte DEGs (Mathys et al., 2019) and MERFISH (this study). Overlapping DEGs are shown in black bars, contrasting DEGs in gray. (H-J) Characterizing Aβ-proximal profiles in oligodendrocytes. (H) Heatmap depicting Aβ-plaque proximal (Aβ+ vs Aβ-) DEGs identified in all, AD or control oligodendrocytes. Adjacent graphs show total number of DEGs (black bars) or up (red) and down-regulated (blue) DEGs. (I) Heatmaps depicting Log2 fold-change in oligodendrocytes, comparing DEGs enriched (black) or depleted (gray) in AD pathology (Mathys et al.) aligned with Aβ+ vs Aβ- MERSCOPE DEGs in all, AD or control oligodendrocytes. (J) Graph summarizing overlap between snRNA-seq oligodendrocyte DEGs (Mathys et al., 2019) and MERSCOPE (this study). Overlapping DEGs with directional Log2 fold-change profiles are shown in black bars, while contrasting DEGs are in gray.

**Figure S7.**
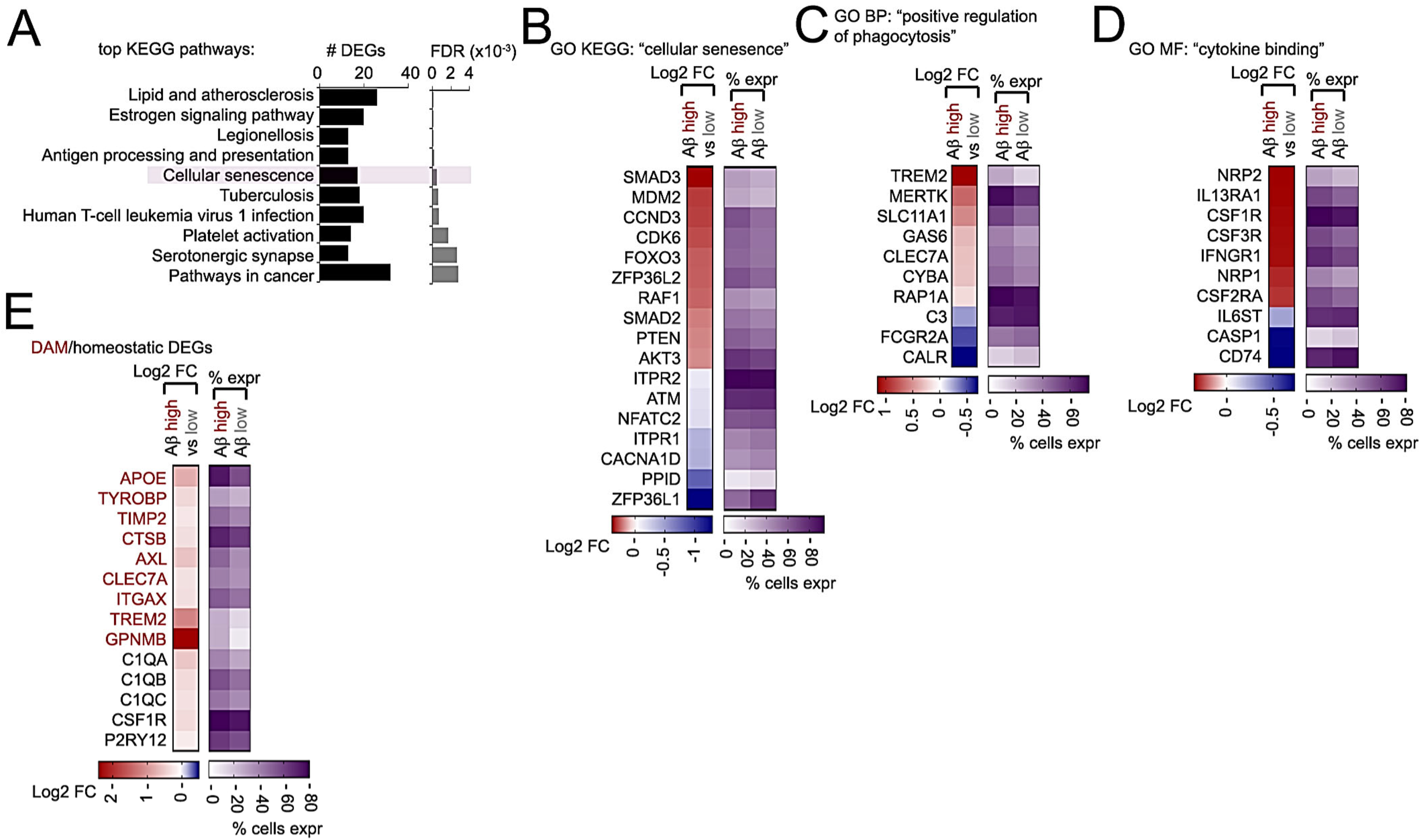
Profiling mapped Aβ-associated MERFISH signatures in snRNA-seq microglia datasets. (A) Top KEGG pathways enriched in Aβ-associated hippocampal microglia DEGs from snRNA-seq analysis (Mathys et al., 2024). (B-E) Heatmaps depicting Log2 fold-change (left) and percentage expression (right heatmaps) for DEGs associated with GO KEGG “cellular senescence” (B), GO BP “positive regulation of phagocytosis” (C), and GO MF: “cytokine binding” categories, in addition to DAM-like and homeostatic microglia DEGs (E).

**Figure S8.**
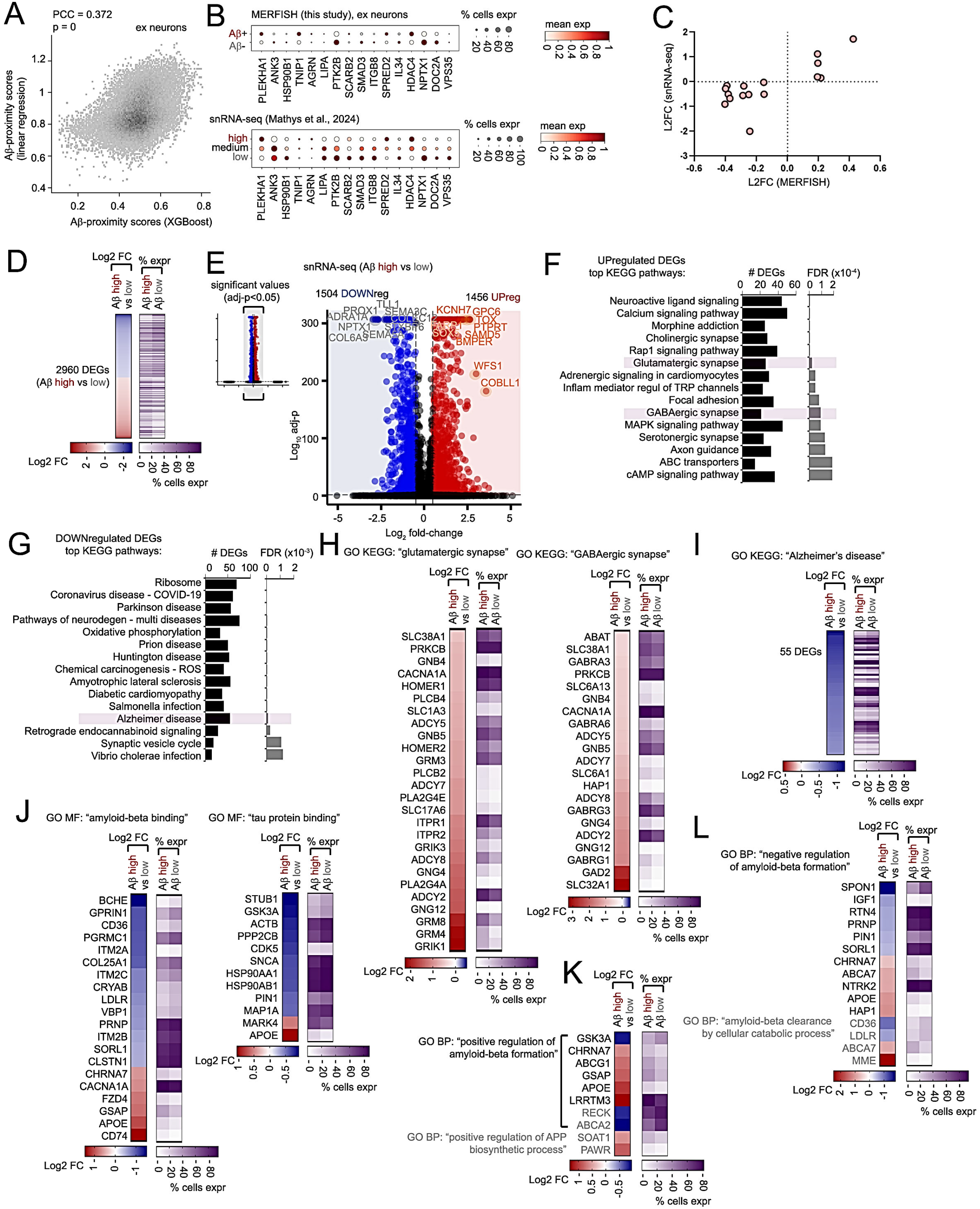
Mapping Aβ-proximal MERFISH excitatory neuron profiles to snRNA-seq datasets. (A-C) Aligning Aβ-proximal transcriptomic MERFISH profiles in excitatory neurons to hippocampal snRNA-seq datasets (Mathys et al., 2024). (A) Scatterplot depicting predicted Aβ-proximity scores by linear regression (LR) (y-axis) and XGBoost (x-axis). Pearson’s correlation coefficient (PCC) and p-value are shown. (B) Dotplots depicting mean and percentage expression of genes enriched or depleted in Aβ-proximal or distal excitatory neurons identified by MERFISH analysis (top dotplot), with similar expression profiles in snRNA-seq datasets with high, medium or low pathology (Mathys et al., 2024; bottom dotplot). (C) Scatterplot depicting L2FC values of genes shown in B in comparing Aβ+ vs Aβ-MERFISH (x-axis) and Aβ-high vs Aβ-low snRNA-seq excitatory neurons (Mathys et al., 2024; y-axis). (D) Heatmaps depicting Log2 fold-change in Aβ high vs low comparisons (left) and percent expression in Aβ high and low cells (right heatmap) of 2,960 DEGs in excitatory neurons mapped for Aβ-proximity in snRNA-seq analysis. (E) Volcano plot of DEGs in Aβ high vs low comparisons using snRNA-seq excitatory neurons (adj-p<0.05, Log2 fold-change >0.5 or <-0.5). Top 10 downregulated (gray) and upregulated (orange) DEGs are highlighted. A volcano plot comprising all DEG data is shown in the inset; non-significant DEGs with Log2 fold-change >10 or <-10 were removed in the full-size volcano plot shown (right). (F, G) Top GO KEGG pathways enriched in upregulated (F) and downregulated Aβ high vs low DEGs (G) mapped from snRNA-seq analysis. (H, I) Heatmaps depicting Log2 fold-change in Aβ high vs low comparisons (left) and percent expression in Aβ high and low cells (right heatmap) of DEGs in GO KEGG “glutamatergic synapse” and “GABAergic synapse” (H) and GO KEGG “Alzheimer’s disease” categories (I). (J-L) Heatmaps depicting Log2 fold-change in Aβ high vs low comparisons (left) and percent expression in Aβ high and low cells (right heatmap) of DEGs in GO MF “amyloid-beta binding” and “tau protein binding” (J) or GO BP “positive regulation of amyloid-beta formation” and “positive regulation of APP biosynthetic process” (K) or GO BP “negative regulation of amyloid-beta formation” and “amyloid-beta clearance by cellular catabolic process” (L).

**Figure S9.**
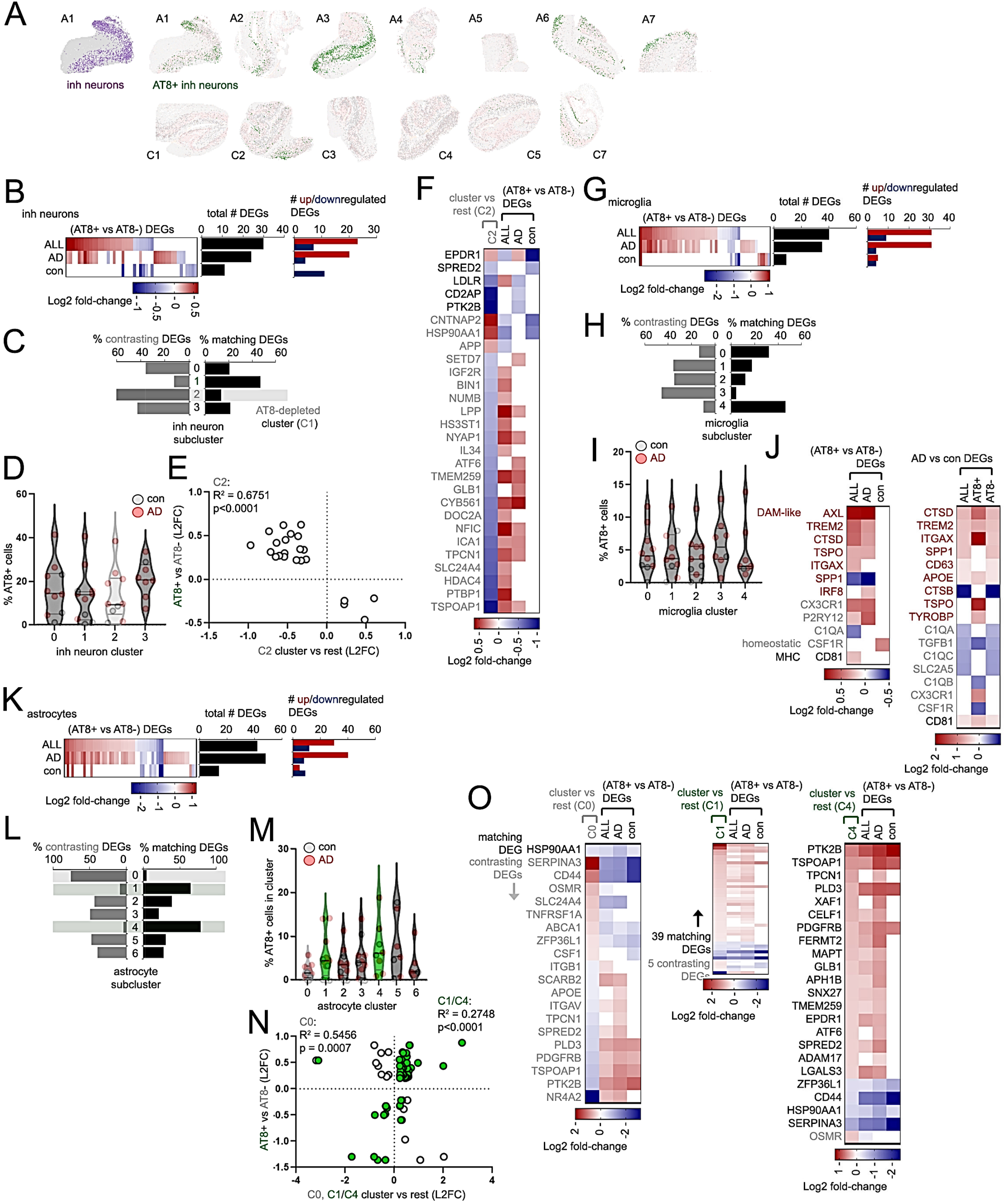
Characterizing expression profiles associated with AT8 ptau in neurons and glia. (A) AT8-enriched inhibitory neurons (green) identified in AD (A1-A7) or control (C1-C6) hippocampus. (B) Heatmap depicting Log2 fold-change of DEGs from AT8+ vs AT8-inhibitory (inh) neurons in all, AD or control hippocampus. Adjacent graphs depict number of total (black bars) and up (red) or down-regulated (blue) inh neuron DEGs. (C) Graph depicting alignment of % matching (black bars) or contrasting (gray bars) inhibitory neuron DEGs from cluster vs rest in clusters 0 through 3 in comparison to AT8+ vs AT8-DEGs. Cluster 2 features a high number of contrasting, and reduced number of matching AT8+ DEGs (potential AT8-depleted cluster). (D) Percentage of AT8+ inhibitory neurons in each inh neuron cluster; cluster 2 as a potential AT8-depleted (light gray) is highlighted. Plots representing percentages for ex neurons in each AD (red) and control (gray) hippocampus are shown. (E) Scatterplot depicting Log2 fold-change (L2FC) in inh neuron cluster 2 (gray) DEGs; cluster vs rest (x-axis) and AT8+ vs AT8-(“ALL” DEGs, y-axis) L2FC values for individual DEGs, and R^2^/p-values for C2 trends are shown. (F) Heatmap showing Log2 fold-change profiles of inh neuron cluster 2 profiles aligned with AT8+ vs AT8-DEGs in all, AD or control inh neurons. (G) Heatmap depicting Log2 fold-change of DEGs from AT8+ vs AT8-microglia in all, AD or control hippocampus. Adjacent graphs depict number of total (black bars) and up (red) or down-regulated (blue) microglia DEGs. (H) Graph depicting alignment of % matching (black bars) or contrasting (gray bars) microglia DEGs in clusters 0 through 4 in comparison to AT8+ vs AT8-DEGs (cluster 5 was excluded due to few DEGs and limited number of cells within this cluster). (I) Percentage of AT8+ microglia in each microglia cluster. Although DAM-like cluster 4 features an increased number of overlapping, and reduced number of contrasting DEGs in (H), no enrichment of AT8+ microglia in cluster 4 is observed. (J) Heatmaps depicting Log2 fold-change in DAM-like, homeostatic or MHC DEGs in AT8+ vs AT8- comparisons in all, AD or control microglia (left), or AD vs control comparisons in all, AT8+ or AT8- microglia (right). (K-O) Characterization of AT8-associated expression profiles in astrocytes. (K) Heatmap depicting Log2 fold-change of DEGs from AT8+ vs AT8-astrocytes in all, AD or control hippocampus. Adjacent graphs depict number of total (black bars) and up (red) or down-regulated (blue) astrocyte DEGs. (L) Graph depicting alignment of % matching (black bars) or contrasting (gray bars) astrocyte DEGs in clusters 0 through 6 in comparison to AT8+ vs AT8- DEGs. Cluster 0 features a high number of contrasting, and reduced number of matching AT8+ DEGs (potential AT8-depleted cluster); clusters 1 and 4 comprise increased number of matching AT8+ DEGs, and reduced number of DEGs with contrasting expression compared to AT8+ astrocytes (potential AT8-enriched clusters). (M) Percentage of AT8+ astrocytes in each astrocyte cluster; cluster 0 as a potential AT8-depleted (light gray), and potential AT8-enriched astrocyte clusters 1 and 4 (green) are highlighted. Only cluster 4 shows enrichment of AT8+ astrocytes. Plots representing percentages for astrocytes in each AD (red) and control (gray) hippocampus are shown. (N) Scatterplot depicting Log2 fold-change (L2FC) in astrocyte cluster 0 (gray) or cluster 1 and 4 (green) DEGs; cluster vs rest (x-axis) and AT8+ vs AT8- (“ALL” DEGs, y-axis) L2FC values for individual DEGs, and R^2^/p-values for trends are shown. (O) Heatmap showing Log2 fold-change profiles of astrocyte cluster 0 (left), 1 (middle) and 4 (right heatmap) aligned with AT8+ vs AT8-astrocyte DEGs. Statistical significance in (E) and (N) were determined by simple linear regression analysis.

**Figure S10.**
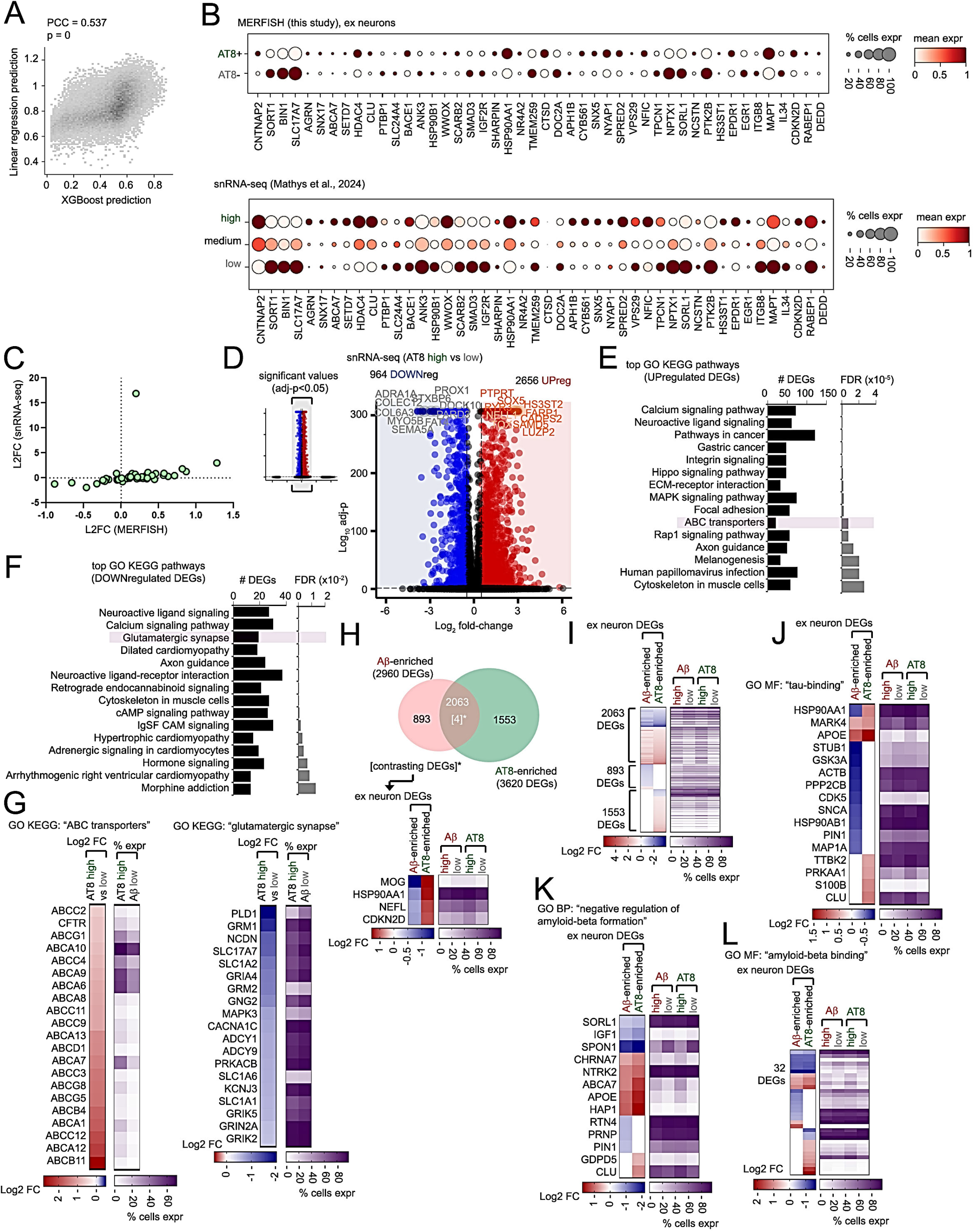
Mapping AT8 ptau MERFISH profiles in snRNA-seq excitatory neuron datasets. (A-C) Aligning AT8 ptau transcriptomic MERFISH profiles in excitatory neurons to hippocampal snRNA-seq datasets (Mathys et al., 2024). (A) Scatterplot depicting AT8 ptau scores of snRNA-seq excitatory neurons using linear regression (y-axis) and XGBoost (x-axis). Pearson’s correlation coefficient (PCC) and p-value are shown. (B) Dotplots depicting mean and percentage expression of genes enriched or depleted in AT8+ (AT8 ptau-enriched) or AT8-(low/no AT8 ptau) excitatory neurons identified by MERFISH analysis (top dotplot), with similar expression profiles in snRNA-seq datasets with high, medium or low pathology (Mathys et al., 2024; bottom dotplot). (C) Scatterplot depicting L2FC of AT8-associated MERFISH genes (x-axis) mapped in snRNA-seq datasets (Mathys et al., 2024; y-axis). (D) Volcano plot of DEGs in AT8 high vs low comparisons in mapped excitatory neurons from snRNA-seq analysis (adj-p<0.05, Log2 fold-change >0.5 or <-0.5). Top 10 downregulated (gray) and upregulated (orange) DEGs are highlighted. A volcano plot comprising all DEG data is shown in the inset; non-significant DEGs with Log2 fold-change >10 or <-10 were removed in the full-size volcano plot shown (right). (E, F) Top GO KEGG pathways enriched in upregulated (E) and downregulated AT8 high vs low DEGs (F) from snRNA-seq AT8 high vs low analysis. (G) Heatmaps depicting Log2 fold-change in AT8 high vs low comparisons (left) and percent expression in AT8 high and low cells (right heatmap) of DEGs in GO KEGG “ABC transporters” and “glutamatergic synapse” pathways as indicated. (H) Venn diagram depicting DEG overlap between Aβ-proximal (left, red) and AT8+ excitatory neurons (right, green). Heatmaps depicting Log2 fold-change in Aβ or AT8 high vs low comparisons (left) and percent expression in Aβ or AT8 high and low cells (right) in contrasting DEGs in Aβ and AT8 comparisons. (I-L) Heatmaps depicting Log2 fold-change in Aβ or AT8 high vs low comparisons (left) and percent expression in Aβ or AT8 high and low cells in all overlapping and unique DEGs (I), or DEGs related to GO MF “tau-binding” (J), GO BP “negative regulation of amyloid-beta formation” (K), or GO MF “amyloid-beta binding” categories (L).

**Figure S11.**
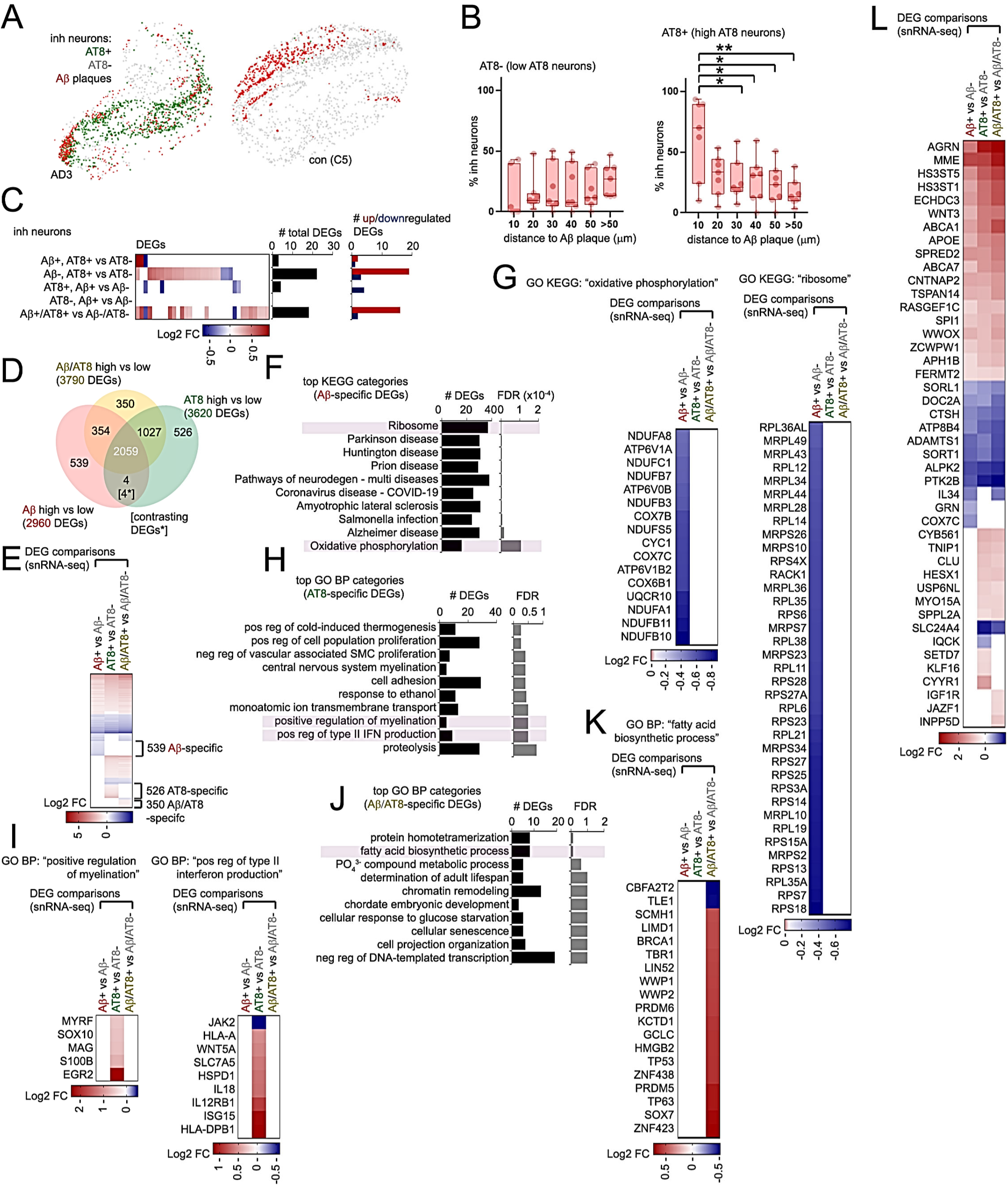
Characterizing spatial profiles associated with Aβ and AT8-tau in neurons. (A-C) Characterizing Aβ/AT8 profiles in inhibitory neurons. (A) Representative hippocampal tissue sections from AD (AD3) and control (C5) individuals. Aβ plaques (red), AT8+ (green) and AT8-inhibitory neurons (gray) are indicated within hippocampal sections. (B) Graphs representing percentages of AT8-(left) and AT8+ inhibitory neurons (right graph) in different distance intervals to Aβ plaques. (C) Heatmap depicting Log2 fold-change in inhibitory (inh) neuron DEGs from the five AT8+ vs AT8-and Aβ+ vs Aβ-comparisons indicated. Adjacent graphs depict number of total (black bars) and up (red) or down-regulated (blue) inh neuron DEGs. (D-L) Characterizing Aβ/AT8-associated excitatory neurons in snRNA-seq datasets mapped using MERFISH profiles. (D) Venn diagram depicting overlap between Aβ high vs low (red), AT8 high vs low (green) or Aβ/AT8 high vs low DEGs in mapped snRNA-seq datasets (Mathys et al., 2024). Contrasting DEGs are indicated in square brackets. (E) Heatmap depicting Log2 fold-change of snRNA-seq DEGs in Aβ, AT8 or combined Aβ/AT8 comparisons indicated. DEGs specific to each comparison are indicated. (F-K) GO analysis of DEGs unique to Aβ, AT8 or combined Aβ/AT8 comparisons. DEGs unique to each comparison from (E) were subjected to GO analysis, and top GO KEGG pathways in Aβ-specific DEGs (F) or top GO BP categories for AT8-specific (H) or Aβ/AT8-specific DEGs (J) are shown. Heatmaps depicting DEG profiles for Aβ-specific GO KEGG “oxidative phosphorylation” or “ribosome” (G), AT8-specific GO BP “positive regulation of myelination” or “positive regulation of type II interferon production” (I), or Aβ/AT8-specific “fatty acid biosynthetic process” categories (K) are shown. (L) Heatmap depicting Log2 fold-change of AD-associated GWAS DEGs for Aβ, AT8 or combined Aβ/AT8 comparisons indicated. Statistical significance in (B) was determined by one-way ANOVA with Tukey’s multiple comparisons (*p<0.05, **p<0.01).

**Figure S12.**
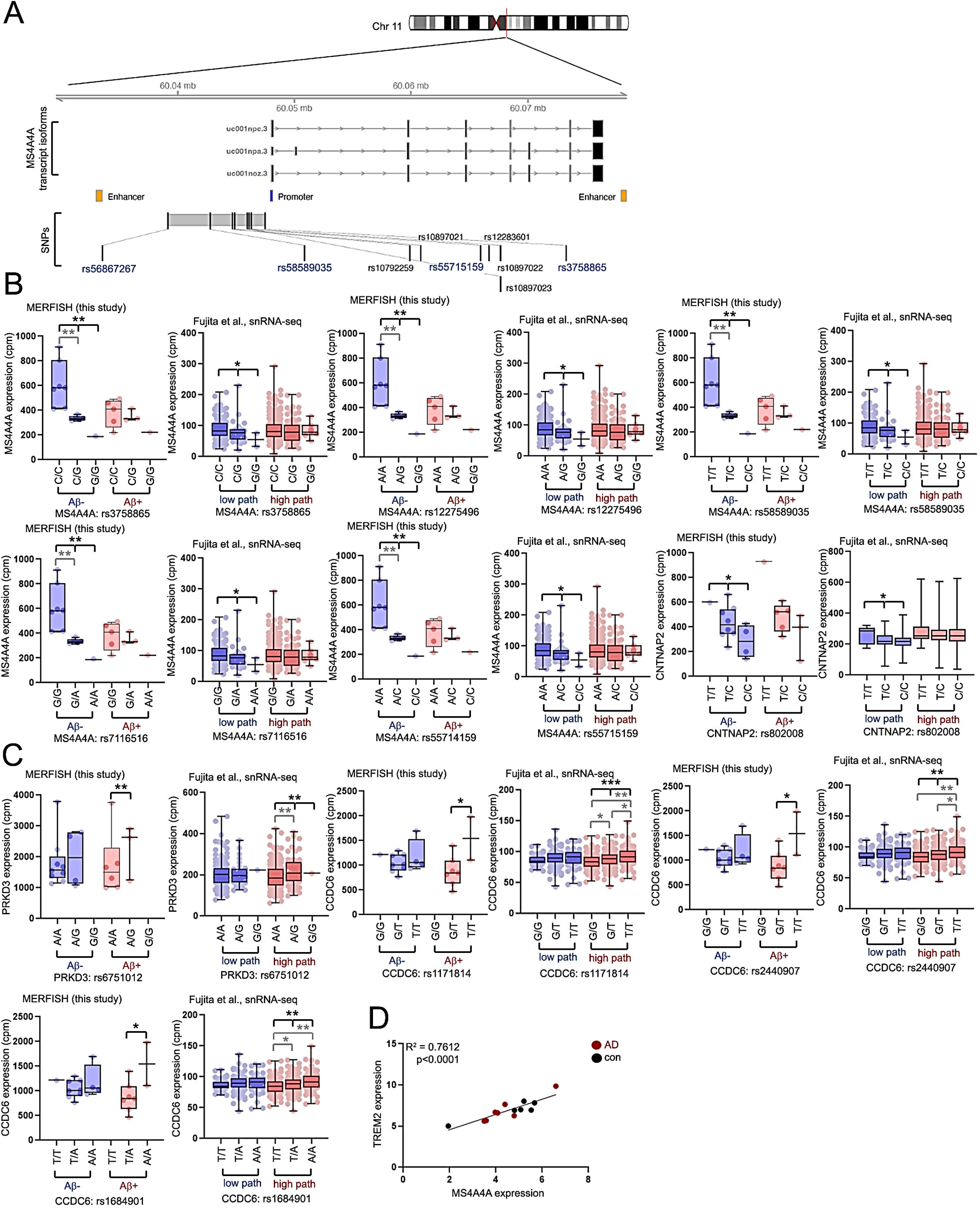
**cis-eQTLs in microglia proximal and distal to Aβ plaques**. (A) Genomic location of the MS4A4A gene locus on Chromosome 11. Regulatory enhancer (orange), promoter (blue) and exons are shown. rs56867267 as well as other SNP variants co-segregated as in a linkage disequilibrium block are shown. SNPs being pathology-dependent eQTLs of MS4A4A are colored in blue. (B, C) Boxplots depicting genotype-dependent microglia expression profiles from eQTLs identified only in microglia distal to (Aβ-) (B) or proximal to Aβ plaques (Aβ+) (C). Genotype-specific expression profiles as characterized by MERFISH analysis in Aβ- and Aβ+ microglia (left graphs) and microglia in low or high pathology categories (right graphs, Fujita et al.) are shown. (D) Correlation between *MS4A4A* and *TREM2* expression in microglia from MERFISH analysis. Individual plots indicate averaged expression values in control (black) and AD (red) hippocampus. Statistical significance in (B) and (C) was determined by a linear model in MatrixEQTL (see method section) and Wilcoxon rank-sum tests for pairwise comparisons between different genotypes. Black indicators represent statistical significance for eQTL analysis, and gray indicators depict statistical significant differences between pairwise genotype analysis (*p<0.05, **p<0.01, ***p<0.001). R2 and p-values in (D) were determined by simple linear regression analysis.

**Figure S13.**
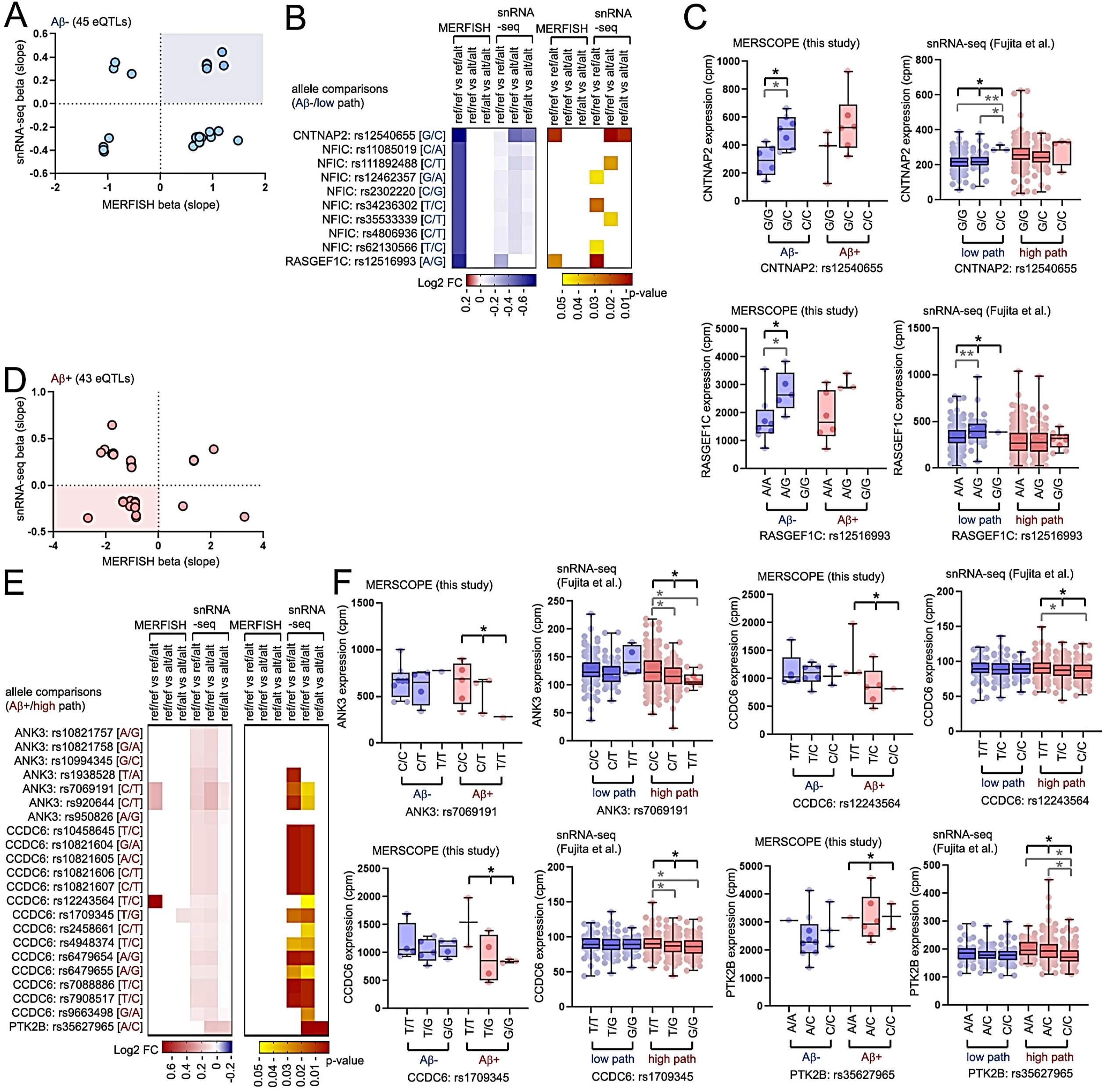
cis-eQTLs responsive to Aβ plaques in microglia. (A-C) Characterizing expression profiles in microglia distal to Aβ plaques. (A) Scatterplot depicting beta (slope) values from snRNA-seq data (Fujita et al., y-axis) and MERFISH data (this study) for eQTLs represented in individual plots. Blue highlight indicates subsequent analysis of eQTLs with positive slope values in both datasets in (B) and (C). (B) Heatmaps depicting Log2 fold-change (left heatmaps) and p-values (right heatmaps) for reference (“ref”) and alternative (“alt”) allele combinations comparisons in MERFISH and snRNA-seq (Fujita et al.) datasets as indicated. (C) Boxplots depicting genotype-dependent microglia expression profiles for transcripts at the loci indicated in microglia distal to (Aβ-) or proximal to Aβ plaques (Aβ+) in eQTLs with positive beta slope values. (D-F) Characterizing expression profiles in microglia proximal to Aβ plaques. (D) Scatterplot depicting beta (slope) values in snRNA-seq datasets (Fujita et al., y-axis) and MERFISH analysis (this study) for eQTLs represented in individual plots. Red highlight indicates subsequent analysis of eQTLs with negative slope values in both datasets in (E) and (F). (E) Heatmaps depicting Log2 fold-change (left heatmaps) and p-values (right heatmaps) for reference (“ref”) and alternative (“alt”) allele combinations comparisons in MERFISH and snRNA-seq (Fujita et al.) datasets as indicated. (F) Boxplots depicting genotype-dependent microglia expression profiles for transcripts at the loci indicated in microglia distal to (Aβ-) or proximal to Aβ plaques (Aβ+) in eQTLs with negative beta slope values. Genotype-specific expression profiles as characterized by MERFISH analysis in Aβ- and Aβ+ microglia (left graphs) and microglia in low or high pathology categories (right graphs, Fujita et al.) are shown. Statistical significance in (C) and (F) was determined by a linear model in MatrixEQTL (see method section) and Wilcoxon rank-sum tests for pairwise comparisons between different genotypes. Black indicators represent statistical significance for eQTL analysis, and gray indicators depict statistical significant differences between allele combinations by pairwise analysis (*p<0.05, **p<0.01).

**Figure S14.**
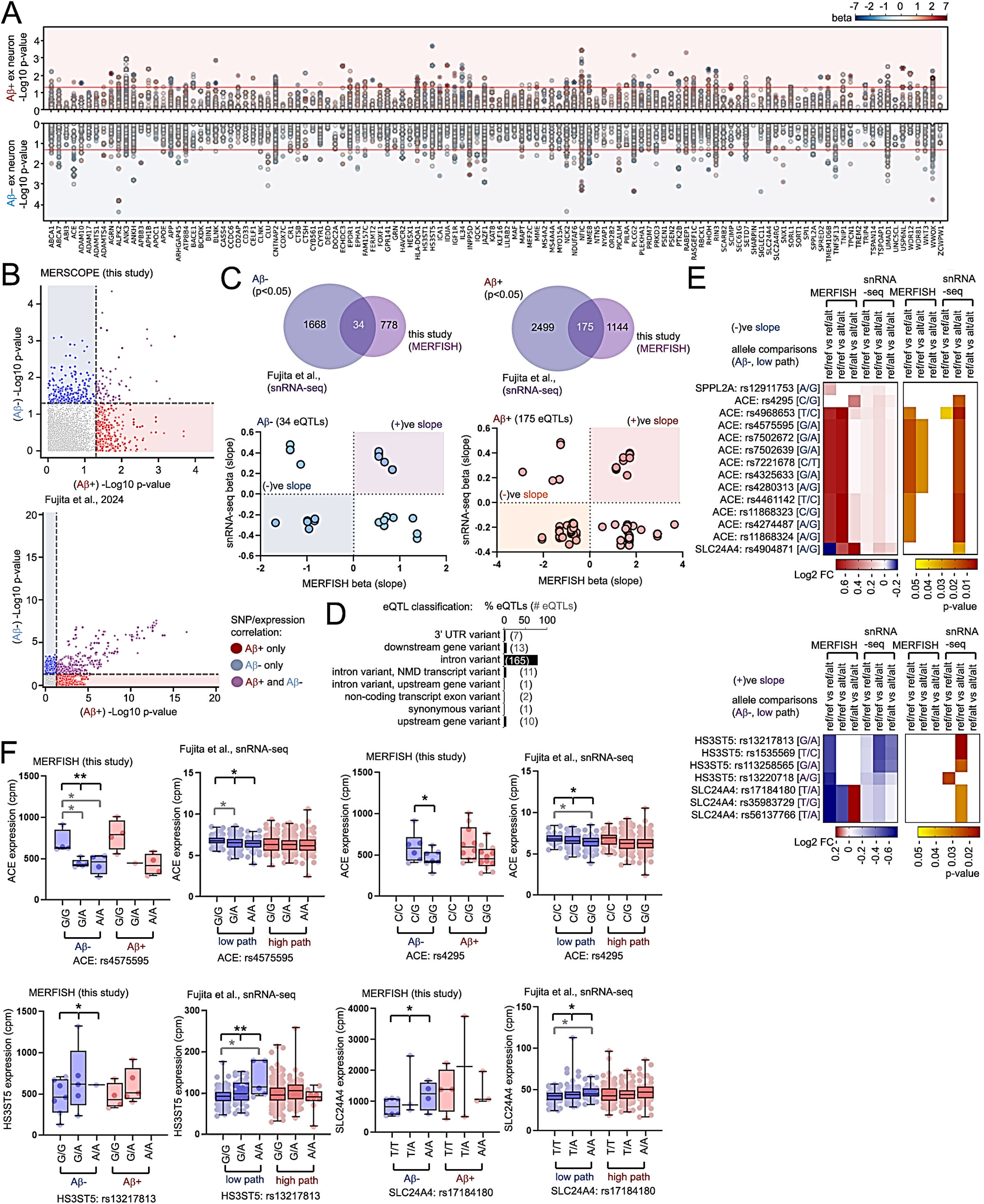
Cis-eQTLs in excitatory neurons proximal and distal to Aβ plaques. (A) Manhattan plot of Aβ+ (top) or Aβ-responsive eQTLs (bottom plots) in excitatory (“ex”) neurons, -log10 p-value and heat-scaled color indicators for beta (slope) are shown. (B) Scatterplot showing -log10(p-value) of eQTLs responsive in either Aβ-or Aβ+ spatial categories in ex neuron MERFISH and snRNA-seq datasets (Fujita et al.). Non-significant hits are shown in gray, Aβ+ (red) or Aβ- (blue) or Aβ+/Aβ- responsive eQTLs (purple) are indicated. (C) Venn diagrams (top panels) depicting overlapping, statistically significant eQTLs identified by snRNA-seq (Fujita et al.) and MERFISH analysis in Aβ-(left) and Aβ+ (right) ex neurons. Scatterplot depicting beta (slope) expression profiles in snRNA-seq datasets (Fujita et al., y-axis) and MERFISH analysis (this study) for eQTLs represented in individual plots. Blue/purple and red/orange highlights indicate eQTLs with matching negative or positive slope values in both datasets in Aβ-and Aβ+ ex neurons. (D) Graph depicting percentage (bars) and number of eQTLs (numbers in brackets) with different effects. (E) Heatmaps depicting Log2 fold-change (left heatmaps) and p-values (right heatmaps) for reference (“ref”) and alternative (“alt”) allele combinations comparisons in MERFISH and snRNA-seq (Fujita et al.) datasets in Aβ- or low pathology with negative beta slope values (top) or positive beta slope values (bottom heatmaps) in ex neurons as indicated. (F) Boxplots depicting genotype-dependent ex neuron expression profiles for ACE rs4575595 and rs4295 with negative beta slope values, or HS3ST5 rs13217813 and SLC24A4 rs17184180 with positive beta slope values distal to (Aβ-) Aβ plaques. Genotype-specific expression profiles as characterized by MERFISH analysis in Aβ- and Aβ+ ex neurons (left graphs) and ex neurons in low or high pathology categories (right graphs, Fujita et al.) are shown. Statistical significance in (F) was determined by a linear model in MatrixEQTL (see method section) and Wilcoxon rank-sum tests for pairwise comparisons between different genotypes. Black indicators represent statistical significance for eQTL analysis, and gray indicators depict statistical significant differences between allele combinations by pairwise analysis (*p<0.05, **p<0.01).

**Figure S15.**
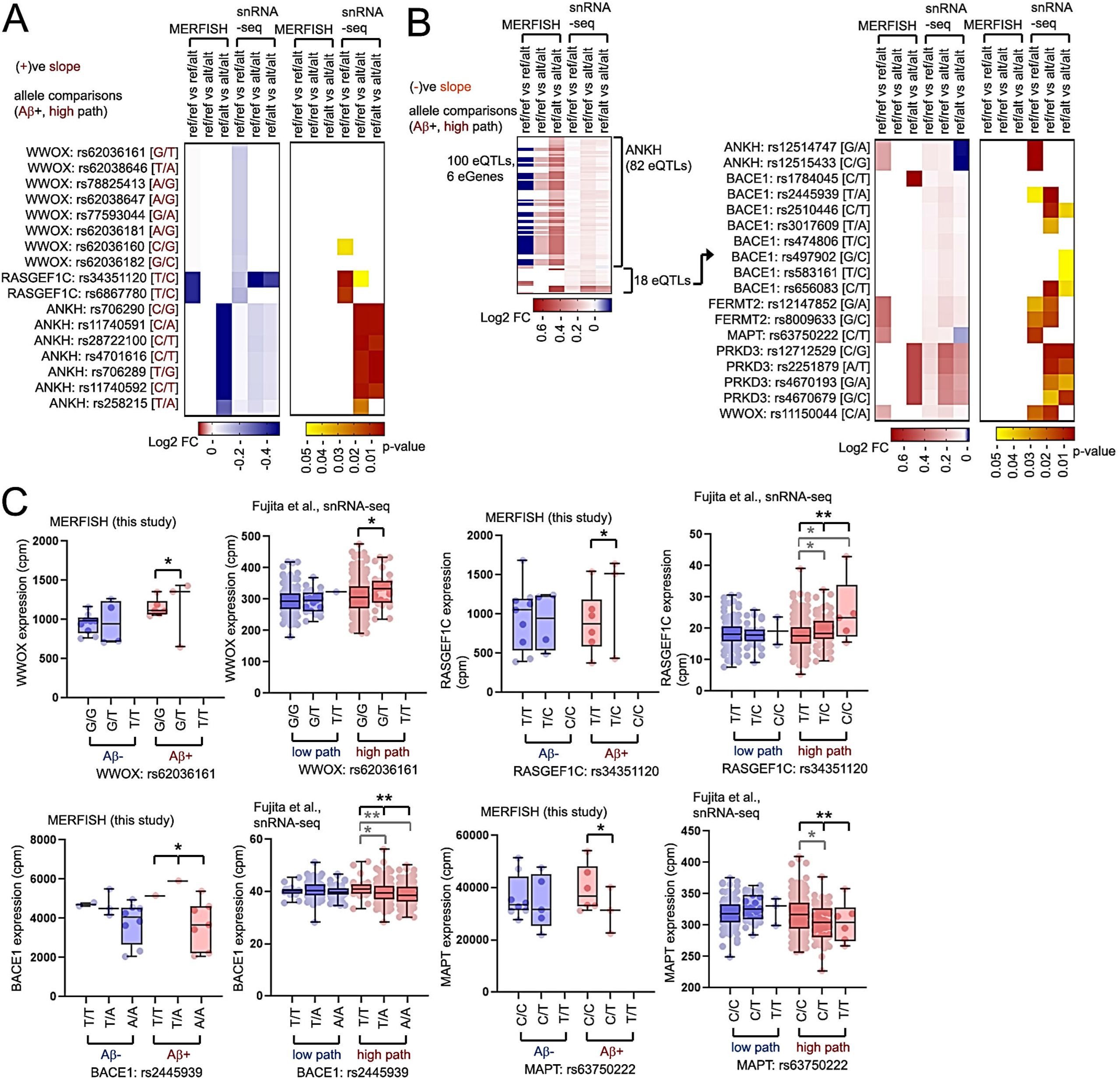
Cis-eQTLs responsive to Aβ plaques in excitatory neurons . (A, B) Heatmaps depicting Log2 fold-change (left heatmaps) and p-values (right heatmaps) for reference (“ref”) and alternative (“alt”) allele combinations comparisons in MERFISH and snRNA-seq (Fujita et al.) datasets in Aβ- or low pathology with positive beta slope values (A) or negative beta slope values (B) in ex neurons as indicated. A total of 82 eQTLs at the ANKH locus were identified, profiles of 18 other eQTLs are shown in (B). (C) Boxplots depicting genotype-dependent ex neuron expression profiles for WWOX rs62036161, RASGEF1C rs34351120, BACE1 rs2445939 and MAPT rs63750222 as indicated. Genotype-specific expression profiles as characterized by MERFISH analysis in Aβ- and Aβ+ ex neurons (left graphs) and ex neurons in low or high pathology categories (right graphs, Fujita et al.) are shown. Statistical significance in (C) was determined by a linear model in MatrixEQTL (see method section) and Wilcoxon rank-sum tests for pairwise comparisons between different genotypes. Black indicators represent statistical significance for eQTL analysis, and gray indicators depict statistical significant differences between allele combinations by pairwise analysis (*p<0.05, **p<0.01).

**Figure S16.**
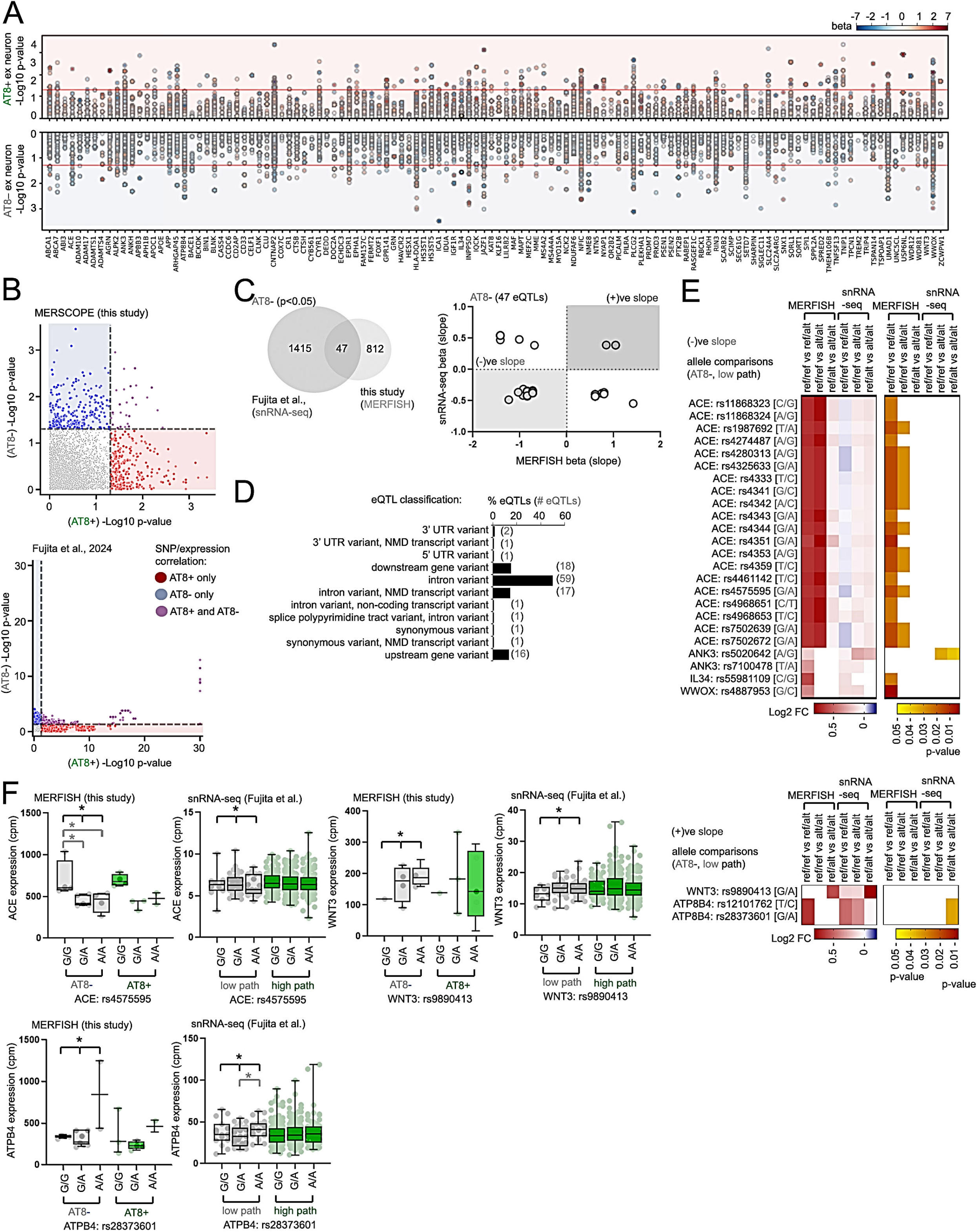
Cis-eQTLs responsive to AT8 pathology in excitatory neurons. (A) Manhattan plot of AT8+ (top) or AT8-eQTLs (bottom plots) results in excitatory (“ex”) neurons, -log10(p-value) and heat-scaled color indicators for beta (slope) is shown. (B) Scatterplot showing -log10(p-value) of eQTLs responsive in AT8-specific spatial categories in ex neuron from MERFISH and snRNA-seq datasets (Fujita et al.). Non-significant hits are shown in gray, AT8+ (red) or AT8-(blue) or AT8+/AT8-responsive eQTLs (purple) are indicated. (C) Venn diagrams (top panel) depicting overlapping, statistically significant eQTLs identified by snRNA-seq (Fujita et al.) and MERFISH analysis in AT8-ex neurons. Scatterplot depicting beta (slope) values in snRNA-seq datasets (Fujita et al., y-axis) and MERFISH analysis (this study) for eQTLs represented in individual plots. Gray highlights indicate eQTLs with matching negative or positive slope values in both datasets in ex neurons. (D) Graph depicting percentage (bars) and number of eQTLs (numbers in brackets) with different effects. (E) Heatmaps depicting Log2 fold-change (left heatmaps) and p-values (right heatmaps) for reference (“ref”) and alternative (“alt”) allele combinations comparisons in MERFISH and snRNA-seq (Fujita et al.) datasets in AT8- or low pathology with negative beta slope values (top) or positive beta slope values (bottom) in ex neurons as indicated. (F) Boxplots depicting SNP-dependent ex neuron expression profiles for ACE (rs4575595), WNT3 (rs9890413), or ATPB4 (rs28373601) in AT8-/low pathology or AT8+/high pathology neurons. Genotype-specific expression profiles as characterized by MERFISH analysis in AT8- and AT8+ ex neurons (left graphs) and ex neurons in low or high pathology categories (right graphs, Fujita et al.) are shown. Statistical significance in (F) was determined by a linear model in MatrixEQTL (see method section) and Wilcoxon rank-sum tests for pairwise comparisons between different genotypes. Black indicators represent statistical significance for eQTL analysis, and gray indicators depict statistical significant differences between allele combinations by pairwise analysis (*p<0.05).

**Figure S17.**
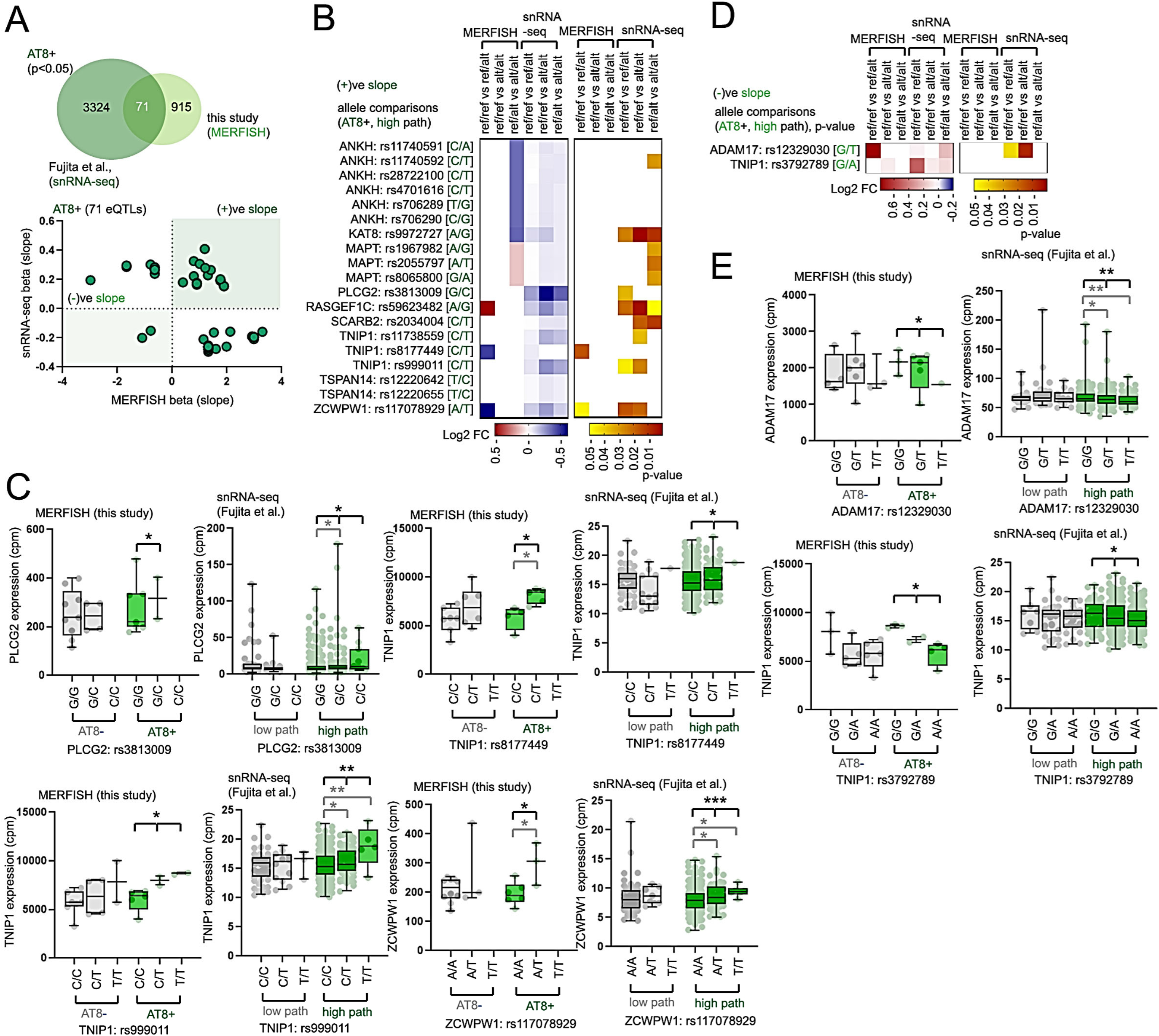
Cis-eQTLs responsive in AT8+ excitatory neurons. (A) Venn diagrams (top panel) depicting overlapping, statistically significant eQTLs identified by snRNA-seq (Fujita et al.) and MERFISH analysis in AT8+ ex neurons. Scatterplot depicting beta (slope) expression profiles in snRNA-seq datasets (Fujita et al., y-axis) and MERFISH analysis (this study) for eQTLs represented in individual plots. Green highlights indicate eQTLs with matching negative or positive slope values in both datasets in AT8+ ex neurons. (B, C) (B) Heatmaps depicting Log2 fold-change (left heatmap) and p-values (right heatmap) for reference (“ref”) and alternative (“alt”) allele combinations comparisons in MERFISH and snRNA-seq (Fujita et al.) datasets in AT8+ or high pathology with positive beta slope values in ex neurons as indicated. (C) Boxplots depicting genotype-dependent ex neuron expression profiles for representative eQTLs identified in (B). Genotype-specific expression profiles as characterized by MERFISH analysis in AT8- and AT8+ ex neurons (left graphs) and ex neurons in low or high pathology categories (right graphs, Fujita et al.) are shown. (D) Heatmaps depicting Log2 fold-change (left heatmap) and p-values (right heatmap) for reference and alternative allele combinations comparisons in MERFISH and snRNA-seq (Fujita et al.) datasets in AT8+ or high pathology with negative beta slope values in ex neurons as indicated. (E) Boxplots depicting genotype-dependent ex neuron expression profiles for ADAM17 (rs12329030) or TNIP1 (rs3792789) identified in (D). Genotype-specific expression profiles as characterized by MERFISH analysis in AT8- and AT8+ ex neurons (left graphs) and ex neurons in low or high pathology categories (right graphs, Fujita et al.) are shown. Statistical significance in (C) and (E) was determined by a linear model in MatrixEQTL (see method section) and Wilcoxon rank-sum tests for pairwise comparisons between different genotypes. Black indicators represent statistical significance for eQTL analysis, and gray indicators depict statistical significant differences between allele combinations by pairwise analysis (*p<0.05, **p<0.01, ***p<0.001).

**Figure S18.**
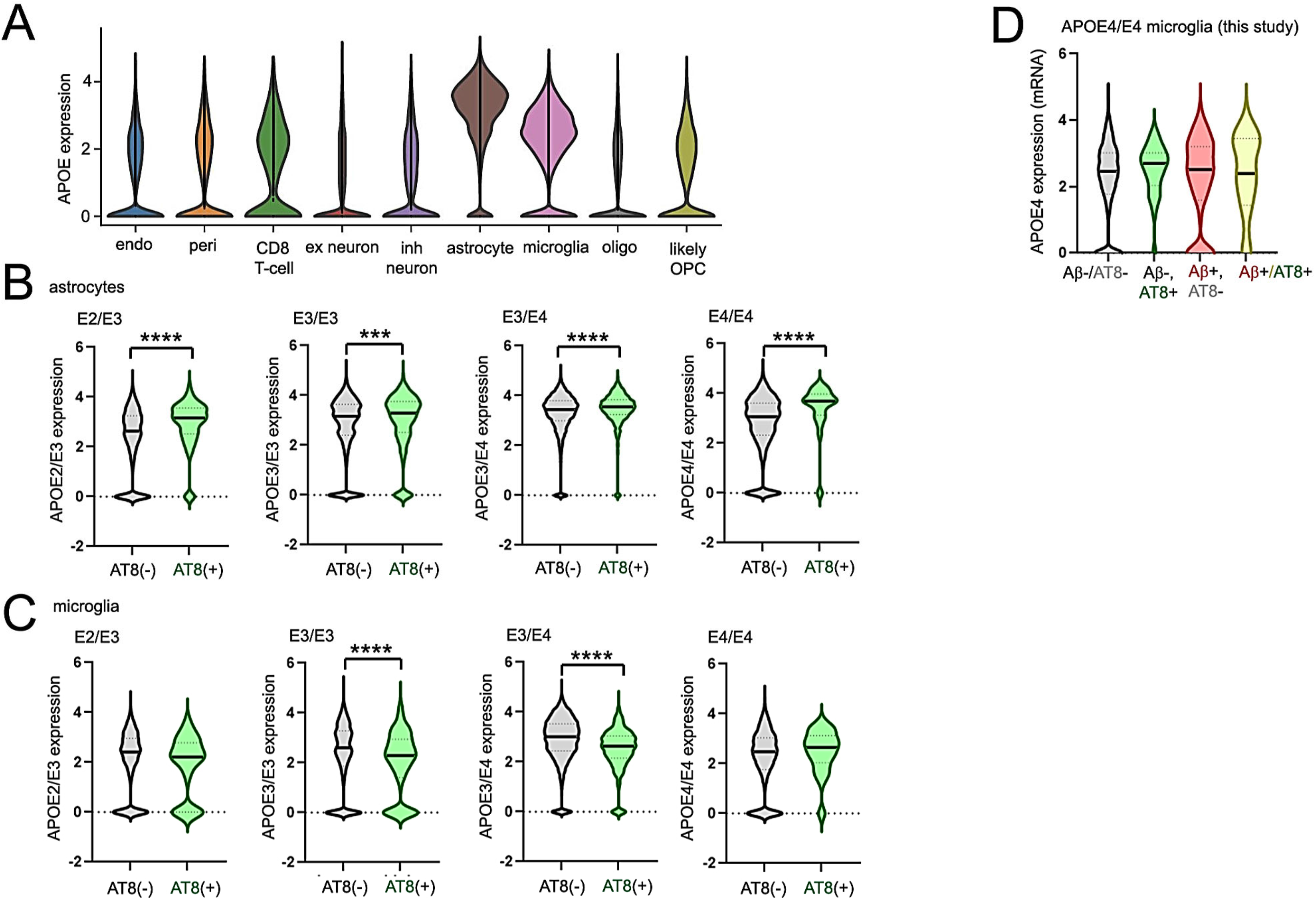
Characterization of astrocyte *APOE* expression in AD brain. (A) Violin plot showing *APOE* mRNA expression in varying cell types in the MERSCOPE dataset. (B, C) Violin plot depicting *APOE* expression in APOE2/E3, E3/E3, E3/E4 or E4/E4 hippocampus in astrocytes (B) or microglia (C) distal to (AT8-) or proximal (AT8+) to AT8+ neurons. (D) *APOE* expression quantified in APOE4/E4 microglia in proximity to Aβ or AT8+ neurons as indicated from MERFISH analysis. Statistical significance was determined by unpaired Student’s t-test in (B, C), or one-way ANOVA in (D) (***p<0.001, ****p<0.0001).

## Inventory of supplemental tables

Table S1 – List of hippocampal control and AD tissues used in the study, donor information

Table S2 – List of 120 GWAS AD genes and description of variants in AD from published studies

Table S3 – Description of MERFISH probes

Table S4 – AD vs con DEGs in various cell types, GO analysis of AD vs con DEGs

Table S5 – DEG analysis (cluster vs rest; AD vs con) of transcriptional clusters in glia and neurons Table S6 – DEG analysis of glia and neurons in proximity/distal to Aβ plaques

Table S7 – snRNA-seq (Mathys et al., 2024) microglia Aβ+ vs Aβ-DEGs derived from mapped Aβ-proximal MERFISH profiles

Table S8 - DEG analysis of AT8+ vs AT8-neurons, or glia in proximity/distal to AT8+ neurons

Table S9 - snRNA-seq (Mathys et al., 2024) excitatory neuron Aβ+, AT8+ and combined Aβ/AT8+ DEGs derived from MERFISH profiles

Table S10 - DEG analysis of Aβ/AT8+ vs Aβ/AT8-neurons or glia

Table S11 – eQTL analysis in Aβ-proximal/distal microglia, overlapping eQTLs in MERFISH and snRNA-seq (Fujita et al.) datasets

Table S12 - eQTL analysis in Aβ-proximal/distal excitatory neurons, overlapping eQTLs in MERFISH and snRNA-seq (Fujita et al.) datasets

Table S13 - eQTL analysis in AT8+ excitatory neurons, overlapping eQTLs in MERFISH and snRNA-seq (Fujita et al.) datasets

## Notes

### Competing Interest Statement

The authors have declared no competing interest.

